# Evolutionarily informed gene sets reveal conserved and lineage-modified transcriptional programs during vertebrate forebrain evolution

**DOI:** 10.64898/2026.03.24.713832

**Authors:** Haowen He, Jeffrey T. Streelman, Peng Qiu

## Abstract

The vertebrate forebrain exhibits striking diversity in anatomical architecture, yet is built from deeply conserved gene regulatory programs and cell types that underlie shared neural functions and behaviors. Understanding how these conserved cellular programs are maintained and modified across ∼500 million years of vertebrate evolution requires systematic cross-species single-cell comparisons, a challenge compounded by complex gene evolutionary histories. Here we present scGENUS, a multispecies comparative framework that derives evolutionarily informed gene sets from a global homology graph and represents cells in a shared, interpretable gene-set feature space. Applying this framework to forebrain profiles from eleven vertebrate species, from sea lamprey to human, we construct a unified cross-vertebrate cell atlas. We find that conserved transcriptional programs define stable cell-type identities across vertebrates, with evolutionary divergence occurring predominantly within, rather than between, cell types. Gene-set conservation scales with evolutionary age, whereas lineage-modified programs reflect coordinated clade-level remodeling. Radial glia exhibit a conserved fate bifurcation into neurogenic and gliogenic trajectories with lineage-dependent modulation of transcriptional dynamics. Finally, human neuropsychiatric GWAS signals map onto conserved neural substrates across vertebrates. Together, our results demonstrate that vertebrate forebrain evolution proceeds through lineage-specific tuning of deeply conserved transcriptional programs embedded within stable cellular architectures.

## INTRODUCTION

Vertebrates share a remarkable capacity for complex cognition, driven by forebrain circuits that regulate behaviors ranging from spatial navigation and sensory integration to social decision-making^1,2^. Yet, the anatomical substrates underlying these functions have undergone distinct evolutionary expansions, exhibiting markedly divergent architectures across lineages: the six-layered neocortex is a uniquely mammalian structure, whereas the pallium of sauropsids (birds and reptiles) mainly comprises the dorsal ventricular ridge (DVR), which is organized into discrete nuclear rather than laminar subdivisions, and that of ray-finned fishes is shaped by a unique outward folding known as eversion^3,4^. Consequently, establishing homologies between these forebrain territories has been historically challenging, leading to conflicting models of vertebrate brain evolution that remain unresolved^5^. Comparative molecular analyses, however, demonstrate that the evolution of cell-type identity is largely decoupled from tissue topology; despite divergent anatomical configurations, underlying cell populations defined by shared transcriptional programs remain deeply conserved^6–9^. Determining the extent to which these conserved cell types and gene programs are maintained, modified, and redeployed across divergent forebrain architectures is therefore central to reconstructing vertebrate forebrain evolution.

Recent whole-brain single-cell and spatial transcriptomic atlases across diverse vertebrates have advanced our understanding of forebrain organization and cellular diversity. These studies extend beyond canonical mammalian models to non-model lineages, revealing conserved cell-type composition and ancestral features such as the persistence of neurogenic radial glia and widespread adult neurogenesis, which are regionally restricted in the mammalian brain^10^. Despite these advances, a systematic cellular comparison across the vertebrate tree of life remains lacking, as most existing cross-species analyses are restricted in phylogenetic scope, often focused on mammals or limited to pairwise comparisons within specific lineages.

Extending comparative single-cell analysis across ∼500 million years of vertebrate evolution poses a substantial computational challenge. Complex gene evolutionary histories, shaped by pervasive duplication, lineage-specific expansion, and subfunctionalization, complicate cross-species comparisons that rely on one-to-one gene correspondence^11–13^. Recent approaches address this by aligning cells in shared latent spaces while incorporating homology information through sequence similarity (SAMap), protein language model embeddings (SATURN), or graph neural network representations (CAME)^13–15^. In practice, these strategies effectively construct shared cross-species cell embeddings, while gene-level contributions are incorporated or recovered implicitly within the learned representations rather than serving as explicit features for integration. As a result, disentangling the specific gene families and regulatory mechanisms that underlie cell-type correspondence across species can be less direct, limiting biological interpretability.

Here we present scGENUS (single-cell Graph-Enabled Network Unifying Species), a comparative framework for cross-species single-cell analysis across phylogenetically distant vertebrates that captures complex many-to-many gene homology relationships beyond strict one-to-one orthology. scGENUS constructs a global homology graph from reciprocal protein sequence similarity, partitions it into evolutionarily informed gene sets, and represents single-cell transcriptomes in an interpretable gene-set feature space. By shifting the emphasis from cell-level embedding alignment alone to evolutionarily informed feature engineering, scGENUS complements existing integration approaches and enhances biological interpretability in cross-species single-cell analyses.

We apply scGENUS to single-cell forebrain transcriptomes from eleven vertebrate species and establish, to our knowledge, the most comprehensive comparative vertebrate forebrain atlas to date. Our results demonstrate that core cell-type identities are broadly conserved across vertebrates, with substantial evolutionary divergence arising primarily through lineage-specific modulation of transcriptional programs within cell types rather than through the widespread emergence of novel cell types. We further uncover a conserved radial glial bifurcation into neurogenic and gliogenic trajectories with lineage-dependent variation in transcriptional dynamics. Finally, this gene-set framework enables cross-species mapping of human neuropsychiatric genetic risk onto conserved neural cell populations. Collectively, this work establishes evolutionarily informed feature engineering as a scalable approach for understanding how conserved cell types and transcriptional programs evolve across deep phylogenetic time.

## RESULTS

### scGENUS enables evolutionarily informed integration of single-cell data across vertebrates

Comparative integration of single-cell transcriptomes across deeply diverged vertebrate species faces two central challenges: first, complex gene evolutionary histories shaped by duplication and divergence preclude simple one-to-one gene correspondence; second, existing cross-species integration approaches often rely on latent cell embeddings that obscure the gene-level features underlying cell-type correspondence, limiting biological interpretability. To overcome these limitations, we developed scGENUS, a graph-based framework that explicitly incorporates evolutionary homology into feature construction prior to data integration. By constructing a global gene homology graph across species and deriving evolutionarily informed gene sets, scGENUS transforms single-cell transcriptomes into an interpretable gene-set feature space that preserves conserved core programs alongside lineage-specific variation, while remaining compatible with standard single-cell analysis workflows. This design enables scalable and interpretable integration of single-cell data across deep phylogenetic distances, providing a unified foundation for comparative analyses of vertebrate forebrain cell types (Fig. 1).

**Figure 1.**
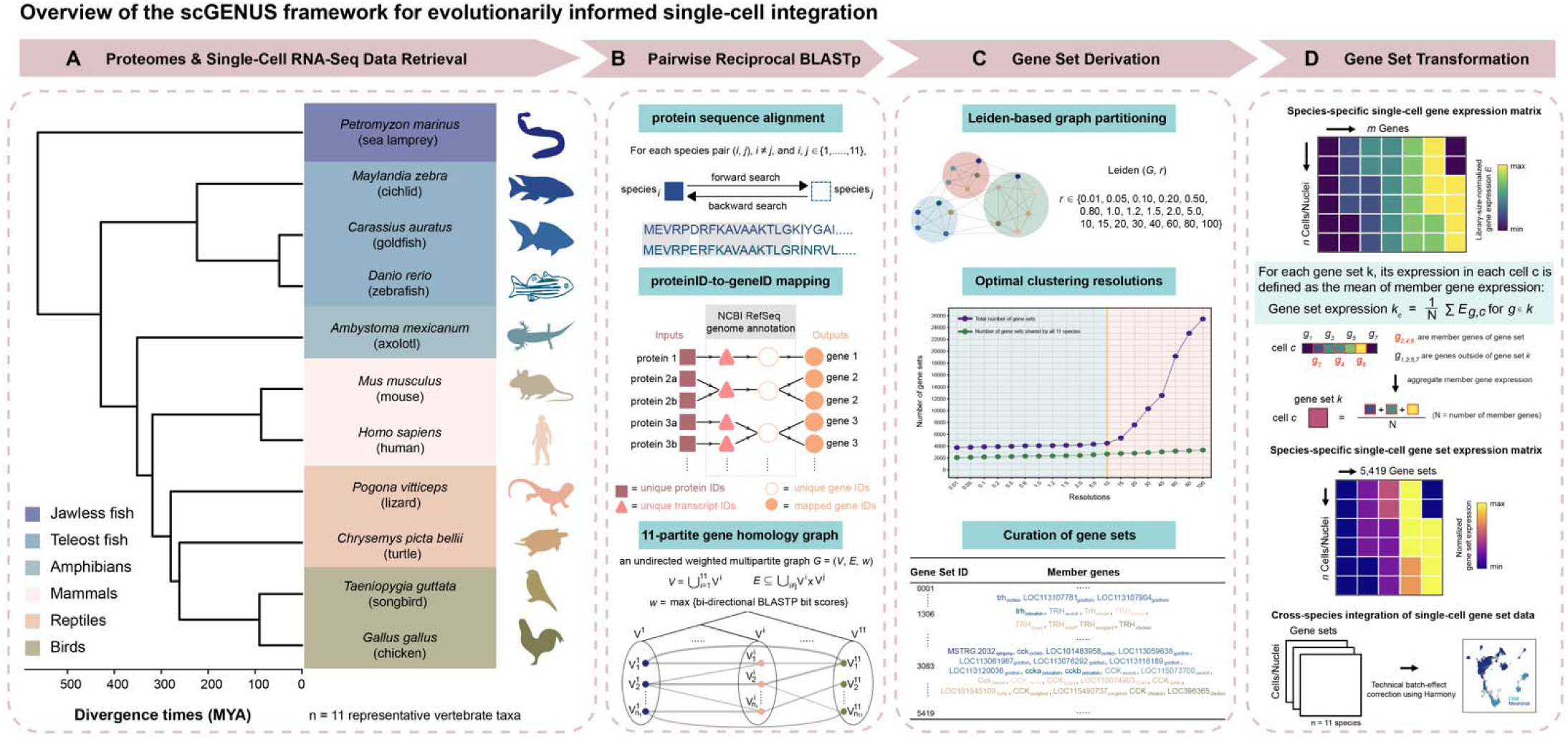
Overview of the scGENUS framework for evolutionarily informed integration of single-cell forebrain data across vertebrates. (A) Curation of single-cell and single-nucleus RNA-sequencing datasets from telencephalic or telencephalon-equivalent forebrain regions across eleven representative vertebrate species spanning approximately 500 million years of evolution, together with corresponding reference proteomes and gene annotations. (B) Construction of a global gene homology graph by performing pairwise reciprocal protein sequence alignments (BLASTp) between all species pairs. Protein sequences are mapped to gene identifiers, and homologous genes across species are connected in an undirected, weighted multipartite graph, with edge weights defined by the maximum of the reciprocal BLASTp bit scores to yield a symmetric measure of sequence similarity. (C) Derivation of evolutionarily informed gene sets by partitioning the gene homology graph using Leiden community detection across a range of clustering resolutions. This multiresolution strategy yields gene sets of varying sizes and degrees of evolutionary conservation across taxa, capturing homologous gene groups at different levels of granularity. Resolutions with sufficient cross-species representation are retained, and redundant gene sets recovered across retained resolutions are collapsed to generate the final collection of gene sets (Supplementary Data File 1). (D) Transformation of single-cell transcriptomes from gene-level to gene-set-level expression by aggregating gene expression values using the mean across genes within each gene set. Gene-set expression matrices are then integrated across species using batch-effect correction in gene-set space, enabling joint visualization and comparative analysis of cell populations across vertebrates.

As a foundation for cross-species analysis of the vertebrate forebrain, we curated single-cell and single-nucleus transcriptomic datasets together with protein sequences and gene annotations from reference genomes for eleven representative vertebrate species spanning approximately 500 million years of evolution (Fig. 1A), including jawless fish (*Petromyzon marinus*^16^), teleosts (*Mchenga conophorus*^17,18^, *Carassius auratus*^19^, and *Danio rerio*^20^), amphibians (*Ambystoma mexicanum*^21^), reptiles (*Pogona vitticeps*^22^ and *Chrysemys picta bellii*^23^), birds (*Taeniopygia guttata*^24^ and *Gallus gallus*^25^), and mammals (*Mus musculus*^26^ and *Homo sapiens*^27^). These datasets were derived from telencephalic or telencephalon-equivalent forebrain regions and collectively provide a broadly sampled and evolutionarily diverse resource for systematic comparison of vertebrate forebrain cell types across deep phylogenetic distances.

Building on these inputs, scGENUS operates in three conceptual stages: constructing a global gene homology graph across species, deriving evolutionarily informed gene sets, and transforming single-cell transcriptomes into a shared gene-set feature space for downstream analysis (Fig. 1).

First, to capture complex homology relationships beyond strict one-to-one orthology, we performed pairwise reciprocal protein sequence alignments using BLASTp between all species pairs and constructed an undirected, weighted multipartite gene homology graph (Fig. 1B). In this graph, nodes represent genes from individual species, and edges connect homologous gene pairs. Edge weights were defined as the maximum of the reciprocal BLASTp bit scores, yielding a symmetric measure of sequence similarity. This graph structure represents many-to-many homology relationships across species, in which multiple genes from one species can be linked to multiple homologous genes in another, reflecting gene duplication and divergence over vertebrate evolution.

Second, to identify evolutionarily coherent gene sets, we partitioned the global homology graph using Leiden community detection across a range of clustering resolutions (Fig. 1C). This multiresolution strategy yielded gene sets of varying sizes, capturing both broadly conserved homology communities and more lineage-restricted groups. We retained resolutions for which at least 50% of resulting communities contained genes from all eleven species to ensure sufficient cross-species representation. Redundant gene sets recovered across resolutions were then collapsed, resulting in a final collection of 5,419 evolutionarily informed gene sets shared across subsets of vertebrate taxa. Gene-set structure showed limited sensitivity to protein length and alignment coverage, and remained stable under alternative edge-weight definitions, stricter BLASTp thresholds, and the retained range of Leiden resolutions, while the final collection of gene sets showed broad cross-species representation and significantly greater Gene Ontology (GO) Biological Process coherence than size-matched random gene groups in both human and mouse (Figs. S1-5).

Finally, single-cell transcriptomes from each species were transformed from gene-level expression matrices into gene-set expression matrices by averaging, within each species, the expression values of all genes assigned to each gene set (Fig. 1D), so that lineage-specific gene expansions contribute through their mean expression rather than increased feature weight. This feature engineering step yields a reduced and biologically interpretable representation of cellular transcriptional profiles while remaining compatible with standard single-cell analysis workflows, as demonstrated by gene-set-based low-dimensional embeddings that recapitulate major cell-type structure (Fig. S6). Gene-set expression matrices from all species were subsequently integrated in gene-set space using Harmony, reducing unwanted technical variation while enabling joint visualization and comparison of cell populations across species. Potentially technical gene sets, including mitochondrial and ribosomal gene sets, had minimal influence on the integrated representation (Fig. S7).

Together, this framework enables scGENUS to integrate single-cell datasets across deep phylogenetic distances using evolutionarily informed gene-set features, providing a unified foundation for comparative analyses of gene programs, cell-type identity, developmental trajectories of cellular differentiation, and evolutionary divergences in the vertebrate forebrain.

### A shared transcriptional atlas of the vertebrate forebrain reveals conserved cell-type identities and lineage-specific divergence

Using the scGENUS gene-set feature space, we integrated forebrain single-cell and single-nucleus transcriptomes from eleven vertebrate species into a shared low-dimensional embedding comprising a total of 300,905 cells. In the integrated UMAP space, cells formed well-defined clusters corresponding to major cell types across species, encompassing glutamatergic and GABAergic neurons; radial glia and astroglial lineage cells (astrocytes and ependymal cells); oligodendrocyte-lineage cells spanning oligodendrocyte precursor cells (OPCs) and mature myelinating oligodendrocytes; immune-associated populations, such as microglia and peripheral immune cells; and non-neural populations, including vascular cells (Fig. 2A). Importantly, the cell-type labels shown here were obtained from the original studies and were not used during integration, indicating that the gene-set-based feature space preserves biologically meaningful cell-type structure in an unsupervised manner across deep phylogenetic distances. Benchmarking against established cross-species integration methods and negative controls further showed that scGENUS achieved the highest overall score across combined measures of cross-species mixing and biological conservation (Fig. S8).

**Figure 2.**
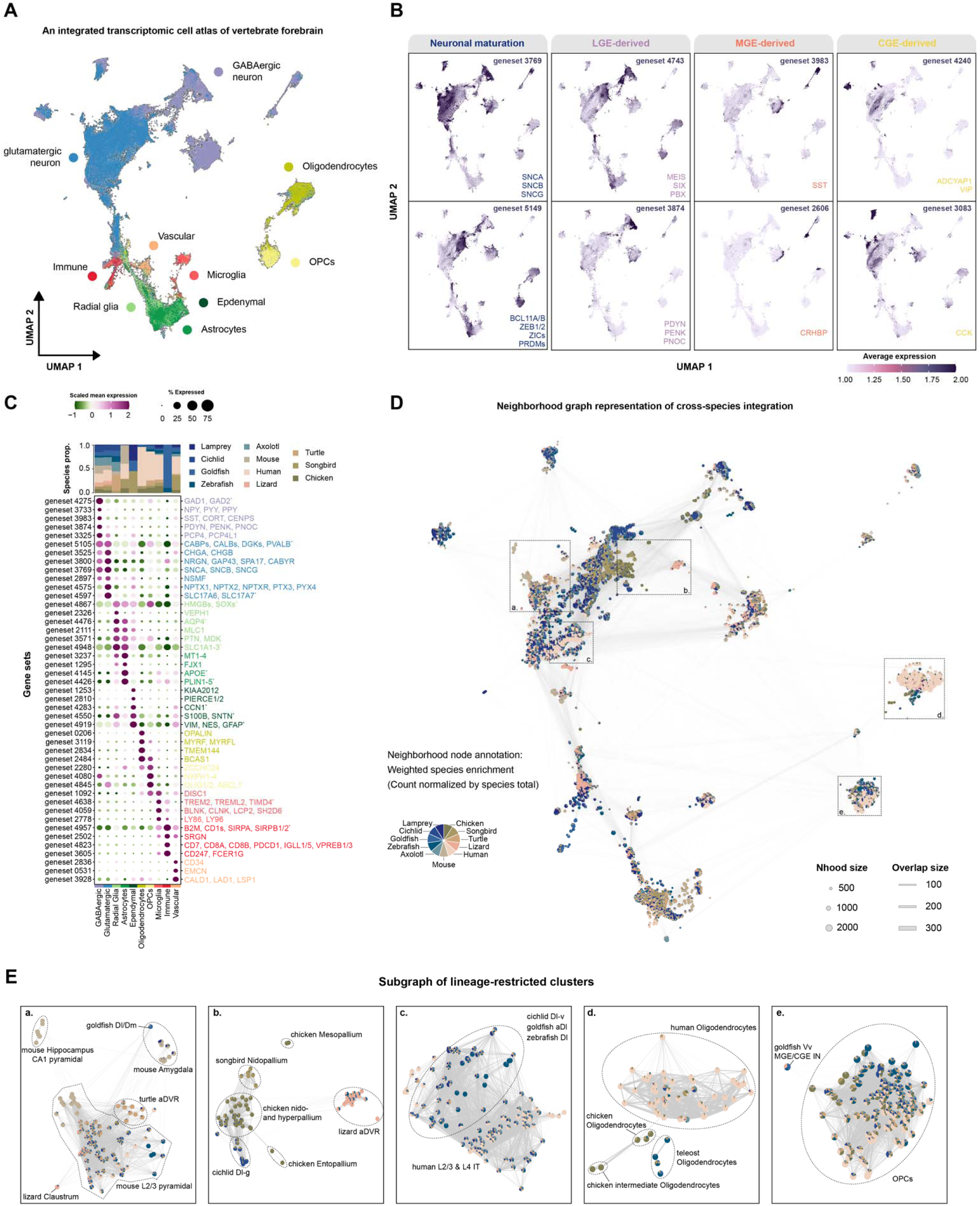
Cross-species integration with scGENUS reveals conserved cell-type identities and lineage-specific divergence across the vertebrate forebrain. (A) UMAP visualization of the integrated vertebrate forebrain atlas generated using the scGENUS gene-set feature space (300,905 cells across eleven species). Major cell types are indicated, including glutamatergic and GABAergic neurons, radial glia, astrocytes, ependymal cells, oligodendrocyte precursor cells (OPCs), oligodendrocytes, microglia, immune and vascular populations (see Supplementary Data File 3). (B) Representative scGENUS gene sets projected onto the integrated vertebrate forebrain atlas. Gene sets associated with mature neuronal states, neuronal development, and LGE-, MGE-, and CGE-derived interneuron programs exhibit structured expression patterns across species. (C) Gene-set-level differential expression identifies cell-type-specific gene programs. Dot plot showing differentially expressed scGENUS gene sets across major forebrain cell types. Gene set identifiers are indicated on the left; representative human gene names from each gene set are shown on the right to facilitate annotation. Asterisks (*) denote large gene sets for which only representative neuronal genes are displayed. Dot size represents the fraction of cells expressing the gene set within each cell type, and color indicates scaled mean expression. Stacked bar plots above each column show species composition within each cell type (see Supplementary Data File 2). (D) A neighborhood graph representation of the integrated vertebrate forebrain atlas. Nodes represent local transcriptional neighborhoods defined in the scGENUS gene-set space; node size reflects the number of cells within each neighborhood, and edges indicate the number of shared cells between adjacent neighborhoods. Pie charts within nodes depict species composition normalized to global species frequencies (see Supplementary Data File 4). (E) Subgraphs highlighting species-biased neighborhoods. Neighborhoods with reduced local species mixing index are shown (see Supplementary Fig. S10 for the full distribution of local species mixing indices across neighborhoods); systematic characterization of strongly species-biased neighborhoods is provided in Fig. S11.

Representative scGENUS gene sets exhibited structured expression patterns across neuronal populations corresponding to distinct neuronal states and subtypes within the integrated space. Several gene sets were associated with neuronal maturation programs conserved across vertebrates (Fig. 2B). In particular, gene set 3768, composed exclusively of synuclein family genes (SNCA, SNCB, SNCG), showed broad and elevated expression across a large fraction of forebrain neurons. Synucleins are predominantly localized to presynaptic terminals of mature neurons, indicating that this gene set marks mature neuronal states in the vertebrate forebrain^28–30^. In contrast, gene set 5149 comprised zinc-finger and homeobox transcription factors with established roles in neural progenitor maintenance and neuronal differentiation, including ZIC (ZIC1-5), ZEB (ZEB1/2), and PRDM (PRDM8, PRDM13, PRDM16) family genes, as well as BCL11A and BCL11B, key regulators of projection neuron development and specification^31–35^. Together, these gene sets delineate complementary transcriptional programs reflecting progenitor-associated and mature differentiated neuronal states within the integrated space. Extending to interneuron diversity, the integrated gene-set space also resolved major subtypes of forebrain interneurons defined by conserved developmental origins. Gene sets corresponding to lateral, medial, and caudal ganglionic eminence programs (LGE, MGE, and CGE, respectively) segregated interneurons into consistent domains across species (Fig. 2B).

Rather than performing differential expression analysis at the level of individual genes, which is restricted to the limited set of genes with one-to-one orthologs across all eleven species, we conducted differential expression in the evolutionarily informed gene-set space defined by scGENUS. Because scGENUS gene sets are derived from a global homology graph, they group evolutionarily related genes, including paralogous and lineage-specific duplications, into shared gene-set features that can be compared across species. This differential expression analysis identified gene sets exhibiting cell-type-restricted expression patterns across the integrated vertebrate forebrain atlas (Fig. 2C). Neuronal gene sets captured conserved programs characteristic of GABAergic and glutamatergic neurons, including canonical markers such as GAD1/2 and SLC17A6/7. Distinct gene sets selectively marked astroglial (e.g., GFAP, AQP4), oligodendrocyte-lineage (e.g., MYRF, ZCCHC24), microglial and immune (e.g., TREM2, SRGN), and vascular (e.g., CD34) populations. Cell-type-specific signals were robustly detected at the level of gene sets despite species differences in underlying gene composition. Moreover, gene-set-level differential expression further enabled identification of species-specific programs within otherwise conserved cell types (Fig. S9). For example, in oligodendrocytes, an OPALIN gene set containing only human and mouse homologs, consistent with its previously reported mammalian specificity, showed preferential expression in human^36^. In radial glia, a WDR73 gene set represented across all eleven species showed selectively elevated expression in cichlid but not in other species, consistent with prior work implicating this gene in lineage-specific social behavior^17^.

Species composition within each cell type was evaluated using a k-nearest neighbor graph constructed with Milo^37^ in the gene-set space to define local neighborhoods of transcriptionally similar cells. For each neighborhood, species enrichment or depletion was quantified relative to global species frequencies, thereby accounting for unequal representation across the dataset. Most neighborhoods contained cells from multiple species, often spanning all major vertebrate lineages, indicating extensive cross-species mixing within major cell types (Fig. 2D). We next computed a local species mixing index using Shannon entropy of species identities within each Milo-defined neighborhood, normalized to global species proportions. Most neighborhoods exhibited high entropy, indicating that cells from multiple vertebrate lineages were intermingled within the same cell types (Fig. S10). A subset showed reduced species mixing (median entropy < 0.55), corresponding to localized enrichment of specific species (Fig. 2E). To systematically characterize these strongly species-biased neighborhoods, we further examined neighborhoods within the low-entropy tail (Fig. S11). Strongly species-biased neighborhoods comprised 533 of 2,260 neighborhoods (23.6%) and contained 14.6% of all cells, and their prevalence varied substantially across species and cell types. These neighborhoods were particularly enriched among oligodendrocytes but were much less frequent among radial glia, indicating that broad cross-species mixing coexists with localized lineage-associated transcriptional divergence. For example, among excitatory neurons, mouse hippocampal CA1 pyramidal cells formed a distinct cluster. As the principal output neurons of the hippocampus, CA1 pyramidal cells comprise heterogeneous subpopulations with distinct molecular and functional properties and established roles in cognitive processing^38,39^. Mouse basolateral amygdala neurons localized in close proximity to goldfish Dl/Dm populations, in agreement with previous pairwise cross-species transcriptomic comparison of goldfish and mouse telencephalon^19^. In addition, avian nidopallial and hyperpallial populations clustered together and showed connectivity to non-avian reptile aDVR populations, mirroring recently reported transcriptomic similarity between these regions and their preferential resemblance to reptilian DVR over mammalian isocortical cell types^9^. Notably, these populations also mapped near cichlid Dl-g, a teleost forebrain region homologous to mammalian retrohippocampal and adjacent retrosplenial/cingulate areas^18^, aligning with the recent finding that selected avian pallial cell types are more closely related to mammalian retrohippocampal regions than to the isocortex^9^. Mesopallial and entopallial populations, by contrast, clustered nearby but remained more distinct within the embedding, reflecting their distinct developmental trajectories relative to the nido-and hyperpallium^9^. Similarly, human oligodendrocytes formed a partially distinct cluster while remaining connected to their counterparts from other species, consistent with gene-set-level differential expression identifying human-enriched transcriptional features in oligodendrocytes.

### Evolutionary cell state divergence and phylogenetic organization of vertebrate forebrain cell types

Cell types are considered “evolutionary units” that retain core regulatory complexes while exhibiting quasi-independent evolutionary divergence across species, reflecting both shared ancestry and lineage-specific cell state evolution^40–42^. To quantify the magnitude of evolutionary divergence within individual cell types relative to differences between cell types, we compared transcriptional distances in principal component (PCA) representations derived from the scGENUS gene-set space (Fig. 3A). For each cell type, we calculated pairwise L1 distances between species-specific centroids, representing evolutionary cell state shifts within that cell type, and contrasted these with L1 distances between centroids of distinct cell types, representing inter-cell-type divergence. Across all eleven vertebrate species, within-cell-type distances were largely separated from between-cell-type distances (Fig. 3B), indicating that evolutionary divergence within a given cell type is generally smaller than divergence between distinct cell types and supporting conservation of major cell-type identities across vertebrates. However, the magnitude of within-cell-type shifts varied substantially across cell types. Oligodendrocytes exhibited comparatively elevated interspecies distances, driven primarily by divergence between amphibians and amniotes and between teleosts and tetrapods. These shifts align with evolutionary remodeling of oligodendrocyte regulatory programs and central nervous system myelin composition, including the emergence of PDGFRA-positive progenitors in tetrapods and transitions from P0-associated programs in teleosts and amphibians toward PLP-dominant programs in amniotes^43–45^. Among neuronal populations, the jawless vertebrate sea lamprey displayed consistently larger distances relative to other vertebrates, reflecting deep evolutionary separation and pronounced transcriptional divergence. Sublineage-restricted analyses across major vertebrate groups (mammals, reptiles, birds, and teleosts; Fig. S12) revealed a similar overall separation between within-and between-cell-type distances. Among these groups, teleosts exhibited comparatively greater within-cell-type divergence. Within teleosts, divergence was heterogeneous across cell types. Radial glia and glutamatergic neurons exhibited the largest interspecies distances, and substantial divergence was observed between cichlid and goldfish across multiple cell types (Fig. 3C). Collectively, these results show that evolutionary cell-state divergence varies across vertebrate forebrain cell types while generally remaining smaller than transcriptional differences between major cell-type identities.

**Figure 3.**
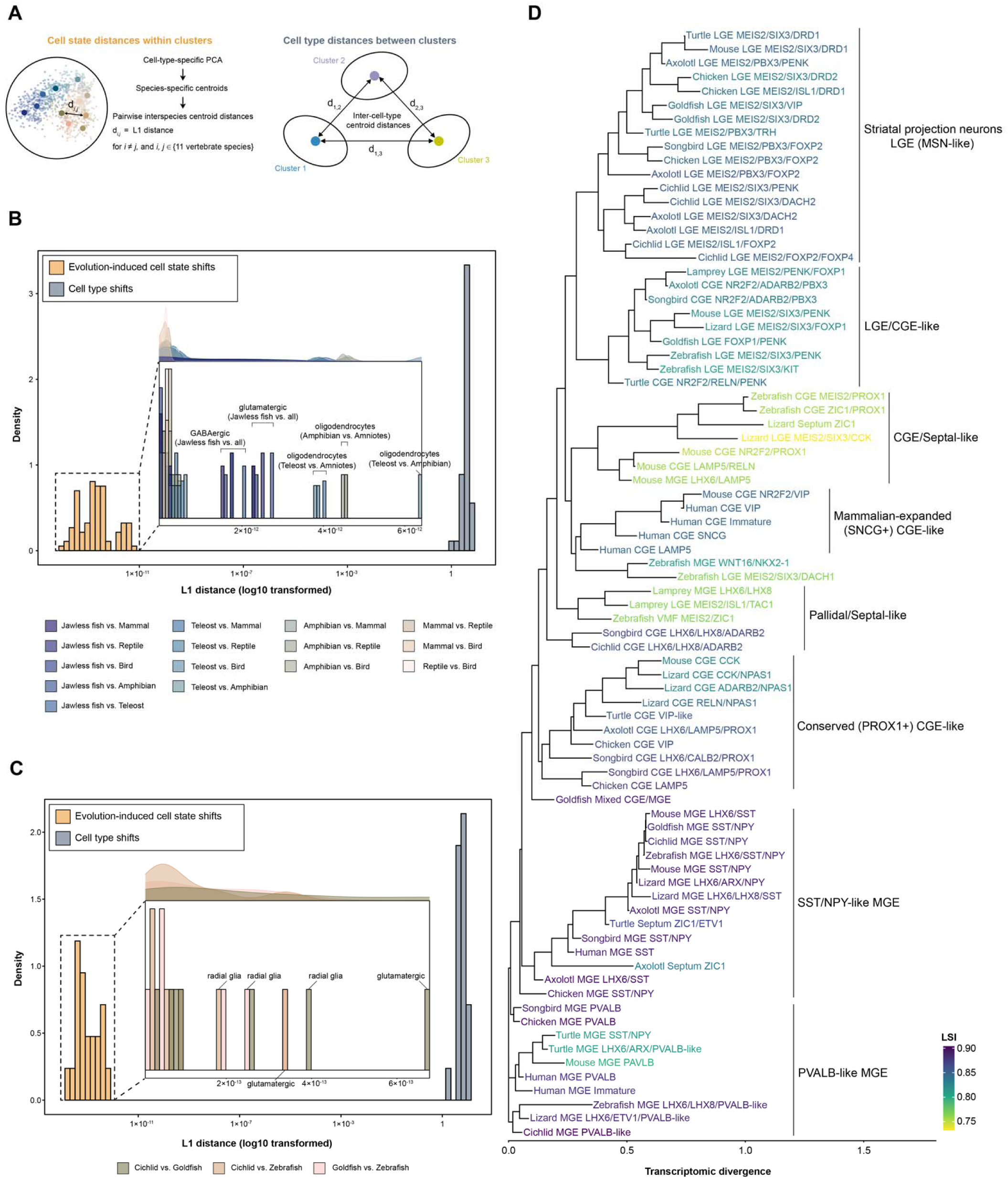
Evolution-induced cell state shifts and phylogenetic organization of GABAergic interneuron subtypes. (A) Strategy for quantifying evolution-induced cell state shifts relative to cell type shifts. Species-specific centroid coordinates were computed for each cell type in PCA representations derived from scGENUS gene-set expression. Pairwise L1 distances between centroids of different species within the same cell type quantified evolutionary cell state shifts and were compared to L1 distances between distinct cell types, representing cell type divergence. (B) Histogram of log10-transformed L1 distances between centroids of different species within each cell type (evolution-induced cell state shifts) versus pairwise L1 distances between centroids of distinct cell types (cell type shifts) across eleven vertebrate species. Comparisons contributing to elevated within-cell-type distances are indicated in the inset. (C) Histogram of log10-transformed L1 distances as in (B), restricted to teleost species. (D) Unrooted, distance-based phylogeny of GABAergic interneuron subtypes inferred from transcriptomic similarity in the integrated PCA space derived from scGENUS gene-set expression (first 50 principal components). Subtypes from eleven vertebrate species are shown at the tips. Branch lengths reflect transcriptomic divergence within the integrated space. Major developmental lineages (LGE-, MGE-, and CGE-derived) are indicated. Tip color represents the leaf stability index (LSI) calculated from 1,000 bootstrap resampling, with values ranging from 0 (low stability) to 1 (high stability).

To examine the evolutionary organization of vertebrate forebrain cell types, we inferred unrooted, distance-based phylogenies for major cell types, including GABAergic and glutamatergic neurons, as well as glial and non-neural populations, from transcriptomic similarity in the integrated PCA space derived from scGENUS gene-set expression, using the first 50 principal components as phylogenetic features (Fig. 3D; Fig. S13). In the phylogeny reconstructed for GABAergic interneurons, interneuron subtypes from across eleven vertebrate species clustered by developmental lineage and molecular identity despite substantial variation in branch lengths, indicating that subtype identity provides a dominant organizational signal over species-level divergence. Comparison of pre-and post-integration representations further showed stronger organization by developmental origin across species (Fig. S14). LGE-, MGE-, and CGE-derived populations formed distinct cross-species clades consistent with their conserved developmental origins. Striatal projection neuron-like subtypes originating from LGE grouped across vertebrates and retained canonical striatal lineage markers, including MEIS2, SIX3, ISL1, FOXP2, and BCL11B, along with differentiated D1-and D2-type medium spiny neuron programs defining the principal projection neurons of the striatum^46,47^. Similarly, MGE-derived SST/NPY-like and PVALB-like interneurons formed coherent clades spanning vertebrate lineages. Within CGE-related populations, both conserved and lineage-partitioned clades were observed, with non-human CGE subtypes clustering separately from human CGE subtypes, which formed a distinct clade characterized by longer branch lengths, consistent with the expanded representation and cortical recruitment of CGE-derived interneurons reported in primates^48,49^. To assess topological robustness, we performed bootstrap resampling of cells within each subtype (1,000 iterations) and quantified tip stability using a leaf stability index (LSI) derived from bootstrap tree variation. LSI values ranged from 0 (low stability) to 1 (high stability) and were concordant within major developmental clades. MGE-derived clades exhibited high stability, whereas mixed-lineage clades, including pallidal/septal-like, CGE/septal-like, and LGE/CGE-like populations, showed reduced stability. These less stable clades were enriched for expression of the neural developmental transcription factors ZIC1 and NR2F2, along with co-expression of LHX6 and LHX8, markers associated with early-born MGE-derived interneuron programs^50^. Overall, tree topology preserved major developmental lineage relationships, whereas branch-length variation reflected lineage-specific transcriptional divergence within the integrated space. Similar phylogenetic organization was observed for glial and non-neural populations (Fig. S13B, C), indicating that cell-type phylogenetic structure is broadly conserved across vertebrates despite extensive evolutionary diversification of cell state programs. Within neuronal populations, we confirmed strong conservation of inhibitory neuron lineages across vertebrates, whereas excitatory neuron populations diverged more substantially over evolution^9,18,21,23–25^ (Fig. S13A).

### Gene-set expression variation across vertebrates reveals lineage-specific divergence

To characterize the magnitude of gene-set expression conservation across vertebrate forebrain cell types, we computed pairwise Spearman correlations of gene-set expression profiles across species (Fig. 4A). For each gene set, species-specific expression vectors were generated across ten major cell types using pseudobulk aggregation. A gene-set expression conservation score, ranging from -1 (highly divergent) to 1 (highly conserved), was defined as the mean pairwise Spearman correlation coefficient across the eleven species, providing a quantitative measure of cross-species expression concordance. The distribution of gene-set expression conservation scores across 5,419 gene sets was strongly right-skewed (Fig. 4B). More than 72% of gene sets exhibited conservation scores above 0.5, including 51.3% classified as conserved (between 0.5 and 0.75) and 21.3% classified as highly conserved (greater than 0.75). In contrast, only 2.5% of gene sets had negative conservation scores. Similar distributions were observed when gene sets were stratified by size (number of genes per set; Fig. S15A), indicating that conservation patterns are largely independent of gene-set size. These results demonstrate widespread preservation of gene-set-level expression programs across vertebrate forebrain cell types. Ranking gene sets by conservation score revealed a continuum ranging from broadly conserved canonical neuronal identity gene sets to phylogenetically restricted gene sets (Fig. 4C). Highly conserved gene sets were enriched for genes defining neuronal identity, including neurotransmitters, synaptic signaling components, genes involved in neuronal homeostasis, and neuropeptides, reflecting preservation of fundamental neuronal identity across vertebrates. In contrast, divergent gene sets displayed restricted phylogenetic distributions and species-biased enrichment patterns, suggesting lineage-specific remodeling of gene-set expression.

**Figure 4.**
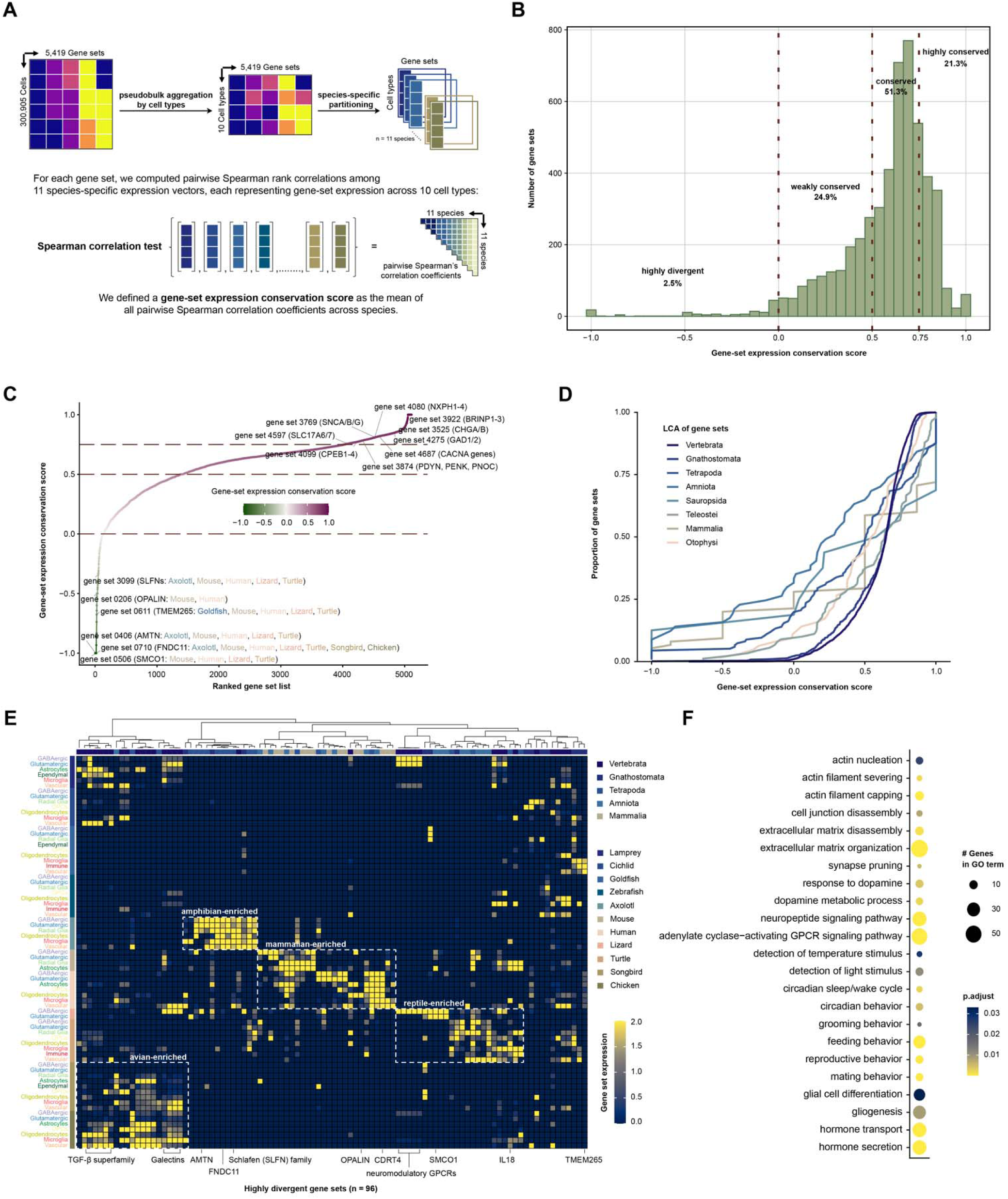
Evolutionary conservation and lineage-specific divergence of gene-set expression across vertebrates. (A) Schematic illustrating calculation of the gene-set expression conservation score. For each gene set, species-specific pseudobulk expression vectors were computed across ten major cell types. Pairwise Spearman correlation coefficients were calculated between species, and the mean pairwise correlation was defined as the gene-set expression conservation score. (B) Distribution of gene-set expression conservation scores across 5,419 gene sets. Scores range from -1 (highly divergent) to 1 (highly conserved). Dashed vertical lines at 0, 0.5, and 0.75 indicate classification thresholds: highly conserved (score 0.75), conserved (0.5 score 0.75), weakly conserved (0 score 0.5), and highly divergent (score 0). (C) Gene sets ranked by conservation score, illustrating a continuum from highly conserved to highly divergent programs, with representative gene sets highlighted. (D) Empirical cumulative distribution functions (ECDFs) of conservation scores grouped by evolutionary age, defined by the last common ancestor (LCA) of the species represented within each gene set. (E) Heatmap of the 96 highly divergent gene sets. Rows represent cell type-species combinations, and columns represent gene sets; values indicate normalized gene-set expression. Row annotations denote species identity, and column annotations indicate the inferred evolutionary age of each gene set. The hierarchical dendrogram reflects similarity relationships among gene sets based on expression profiles. (F) Biological pathway enrichment results of the 96 highly divergent gene sets (human orthologs). Circle size indicates the number of genes associated with each GO term, and color represents FDR-adjusted *p*-values.

Evolutionarily older gene sets are generally expected to exhibit greater cross-species conservation than more recently emerged gene sets. We therefore examined whether gene-set expression conservation varies with evolutionary age. For each gene set, we inferred an evolutionary age, operationally defined here as the most recent common ancestor (LCA) of the species represented within that set, and compared the empirical cumulative distributions of conservation scores across phylogenetic levels (Fig. 4D). Gene sets tracing to deeper evolutionary nodes showed a rightward shift toward high conservation scores, indicating that a larger proportion of ancient gene sets exhibit strong cross-species concordance. In contrast, gene sets associated with more restricted clades, including Tetrapoda, Amniota, and Mammalia, displayed broader distributions with a higher proportion of lower conservation scores, including negative values indicative of divergent expression. This age-dependent pattern indicates that evolutionary depth is a strong predictor of cross-species expression stability. Although only a small fraction of gene sets from deep evolutionary nodes displayed divergent expression, these ancient gene sets were disproportionately represented among the most highly divergent gene sets (Fig. 4E). Among the 96 highly divergent gene sets, evolutionary origins spanned multiple phylogenetic depths, including deep vertebrate nodes as well as Tetrapoda, Amniota, and Mammalia, indicating that expression divergence affects both deeply conserved and more recently emerged gene sets rather than being confined to evolutionarily young gene sets.

Examination of lineage-specific gene-set expression patterns revealed coherent blocks of enrichment corresponding to major vertebrate branches (Fig. 4E). Distinct avian-, amphibian-, mammalian-, and reptilian-enriched clusters were observed, each characterized by coordinated up-regulation across multiple cell types within a lineage. Teleosts and sea lamprey exhibited comparatively fewer enriched divergent gene sets. Notably, divergence was predominantly lineage-wide rather than cell-type-restricted, suggesting coordinated remodeling of gene-set expression at the species or clade level. Gene ontology enrichment analysis of the 96 highly divergent gene sets, performed using human orthologs, revealed overrepresentation of pathways related to cytoskeletal organization and extracellular matrix remodeling, synaptic pruning, neuromodulatory signaling, and behavior-and hormone-associated processes (Fig. 4F). By comparison, highly conserved gene sets were enriched for core neuronal signaling and synaptic transmission pathways (Fig. S15B). Together, these results indicate that evolutionary divergence preferentially affects regulatory and modulatory programs, whereas fundamental neuronal identity and synaptic machinery remain comparatively stable across vertebrates.

### Conserved fate bifurcation of radial glia-like cells into neurogenic and gliogenic trajectories

Radial glia are central to vertebrate forebrain development, serving as multipotent progenitors that generate the neurons and glial cell types populating the brain^51^. In several vertebrate lineages, where regenerative neurogenesis remains widespread, they also persist as a source of new neurons in adulthood^52^. Radial glia can occupy distinct functional states, including quiescence, proliferation, and lineage commitment^53^. In single-cell transcriptomic data, these states form a continuum of differentiation that can be reconstructed through pseudotime inference^54^.

To infer developmental potential of radial glia-like populations across species, we employed CellRank 2 with the CytoTRACE kernel (Fig. 5A), which estimates transition probabilities based on neighborhood connectivity and CytoTRACE-derived differentiation scores without predefined starting states^55^. The inferred vector field revealed coherent directional flows emerging from two spatially distinct radial glia-like regions that aligned preferentially with neurogenic and gliogenic branches, respectively (Fig. 5A). In addition, a discrete quiescent-like radial glia domain exhibited outward-directed transitions, and a distinct origin was observed within the microglial population. To corroborate these putative root states, we examined the expression of canonical gene sets associated with distinct radial glia functional states. The two radial glia-like origin regions were differentially enriched for gene set 5240 (NOTCH1-4; neurogenic competence^56^) and gene set 4595 (FABP7; astrocytic lineage^57^), consistent with their alignment to neurogenic and gliogenic branches (Fig. 5B). A separate quiescent-like radial glia domain was enriched for gene set 4598 (HES1; quiescence maintenance^58^), whereas the independent microglial origin corresponded to enrichment of gene set 4638 (TREM1/2; microglial innate immune receptors^59^). Putative terminal states were enriched for neuronal differentiation markers (for example, NeuroD family genes^60^) and ependymal markers (for example, keratin family genes^61^), validating the inferred branch identities. Notably, gene sets enriched for NFI transcription factors, key regulators of radial glia differentiation, were elevated within both radial glia origin domains and persisted in early progenitor populations^62^. In addition, integrin -enriched gene set (ITGBs) was localized within radial glia domains, particularly neurogenic radial glia, whereas TGF ligand gene set (TGFB1-3) was prominent in microglia (Fig. S16A). This pattern aligns with reported roles for integrin-mediated activation of TGF signaling in microglia development^63^.

**Figure 5.**
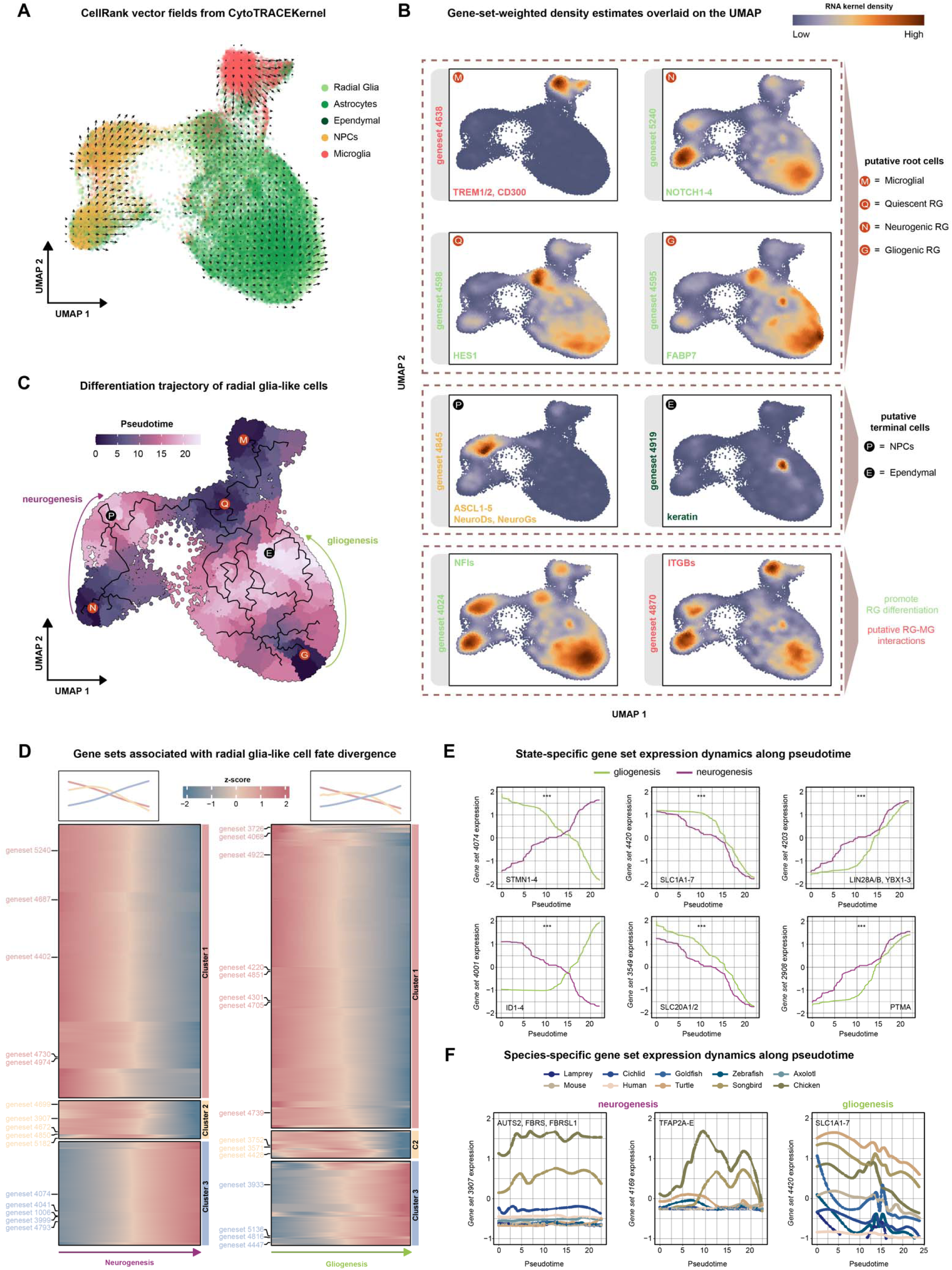
Trajectory inference and pseudotime alignment reveal state-and species-specific gene-set dynamics during radial glia differentiation. (A) CellRank 2 projected vector fields showing inferred cell-cell transition dynamics across radial glia-like, neural progenitor, astrocytic, ependymal, and microglial populations using CytoTRACE-derived developmental potential. Arrows indicate the predicted direction of state transitions in the UMAP embedding. (B) Gene-set-weighted density estimates projected onto the UMAP embedding. Regions of high density correspond to gene set 4638 (TREM1/2; microglia), gene set 4598 (HES1; quiescent-like radial glia), gene set 5240 (NOTCH1-4; neurogenic radial glia), and gene set 4595 (FABP7; gliogenic radial glia). Neuronal progenitor markers (gene set 4845; ASCL1-5; NeuroD/NeuroG) and ependymal markers (gene set 4919; keratin family genes) localize to distinct clusters. Gene sets containing NFI transcription factors and ITGB family genes also show elevated signal in radial glia-like populations. (C) Monocle3 pseudotime alignment of radial glia-like, neural progenitor, astrocytic, ependymal, and microglial populations, initialized from CellRank-inferred starting states. Cells are colored by pseudotime. (D) Gene sets clustered by pseudotime-dependent expression dynamics along neurogenic (left) and gliogenic (right) branches. Heatmaps display z-scored gene-set expression across pseudotime. Insets depict representative module expression patterns. (E) State-dependent gene-set expression dynamics along pseudotime. Curves show smoothed expression for representative gene sets in neurogenic (purple) and gliogenic (green) branches. Asterisks indicate significant differences between branches identified using a natural spline linear regression model (Bonferroni-adjusted *p*-value < 0.0001). (F) Species-specific gene-set expression dynamics along pseudotime within neurogenic and gliogenic branches. Lines represent smoothed expression dynamics for each species across pseudotime (see Supplementary Data File 5).

Using the radial glia root states identified by CellRank as the starting points, we applied Monocle3^64^ to order cells along pseudotime. This analysis resolved a fate bifurcation of radial glia-like cells into neurogenic and gliogenic trajectories, consistent with the directional flows inferred from CellRank (Fig. 5C). Milo neighborhood analysis of the radial glia subset further showed broad cross-species mixing along both branches, indicating that these trajectories reflect shared cell-state transitions rather than species-driven segregation (Fig. S16D,E). Clustering gene sets by their pseudotemporal expression dynamics identified modules associated with fate divergence along neurogenic and gliogenic branches (Fig. 5D). Along the neurogenic trajectory, gene sets exhibited temporally distinct activation patterns and were enriched for synapse assembly, axonogenesis, neuronal projection guidance, Wnt/Notch signaling, and neural precursor proliferation (Fig. S16B). In contrast, gene sets activated along the gliogenic trajectory were enriched for glial differentiation, intermediate filament organization, lipid metabolic processes, and stress-responsive pathways, including NF-B signaling (Fig. S16C). Taken together, these gene-set dynamics delineate distinct transcriptional programs underlying radial glia fate specification.

Next, we tested for state-dependent differences in gene-set expression dynamics between neurogenic and gliogenic branches (Fig. 5E) using a previously established natural spline linear regression model^65^. This analysis identified 428 gene sets exhibiting significant differences in pseudotime-dependent expression between the two branches. Notably, several of these gene sets showed reciprocal behavior across the neurogenic and gliogenic branches. Gene set 4074 (stathmins, STMN1-4), implicated in neural progenitor maintenance and neuronal differentiation^66^, increased progressively along the neurogenic trajectory while declining during gliogenesis. In contrast, gene set 4001 (ID1-4), linked to ependymal maturation^67,68^, showed the opposite pattern, rising during gliogenesis and decreasing along the neurogenic branch. Other gene sets displayed shared activation with branch-dependent differences in magnitude. Gene set 4203 (LIN28A/B, YBX1-3), which promotes neural progenitor proliferation and neurogenic progression^69,70^, increased along both branches but reached higher levels during neurogenesis. Conversely, gene set 4420 (SLC1A family), encoding astrocytic glutamate transporters such as EAAT1/2 (SLC1A3/2), declined along both branches but showed a more gradual reduction during gliogenesis. These results indicate that radial glia fate divergence involves both reciprocal switching of branch-biased gene sets and quantitative modulation of shared regulatory programs. We subsequently examined species-specific variation in gene-set dynamics within each branch (Fig. 5F). Along the neurogenic branch, gene sets 3907 (AUTS2/FBRS family) and 4169 (TFAP2 family), contributing to neuronal specification and maturation^71,72^, displayed stronger activation in the avian lineage relative to other vertebrates. Along the gliogenic branch, gene set 4420 (SLC1A family) exhibited conserved branch-specific dynamics across species but differed in magnitude: amniotes and cichlids showed broadly similar patterns, whereas goldfish, zebrafish, and sea lamprey exhibited more divergent dynamics. These findings suggest that, although the overall fate bifurcation framework of radial glia differentiation is conserved, the magnitude and temporal dynamics of individual gene sets vary in a lineage-dependent manner across vertebrates.

### Translational mapping of cognition-and disease-associated gene sets to conserved neuronal cell types

Complex social and cognitive behaviors are supported by evolutionarily conserved neural circuits across vertebrates, raising the possibility that transcriptional programs implicated in human neuropsychiatric traits may map onto conserved neuronal cell types, as suggested by previous translational work in songbirds and teleost models^73–75^. To assess this, we applied the MAGMA.Celltyping gene-set enrichment framework^76–78^ to human GWAS summary statistics, which maps SNP-level associations to genes and evaluates whether genes within predefined scGENUS gene sets show significant enrichment of GWAS signal. Multiple gene sets exhibited significant enrichment across neuropsychiatric disorders, including ADHD, schizophrenia (SCZ), bipolar disorder (BPD), autism spectrum disorder (ASD), and major depressive disorder (MDD), as well as cognitive traits such as fluid intelligence, prospective memory, and processing speed (Fig. 6A). Functional enrichment analysis of gene sets showing significant GWAS association revealed overrepresentation of biological processes highly relevant to neuropsychiatric pathology, including synapse assembly, postsynaptic organization, axon guidance, regulation of neurogenesis, and Wnt/MAPK signaling (Fig. S17A), indicating that human neuropsychiatric risk preferentially converges on transcriptional programs governing neural development and synaptic organization.

**Figure 6.**
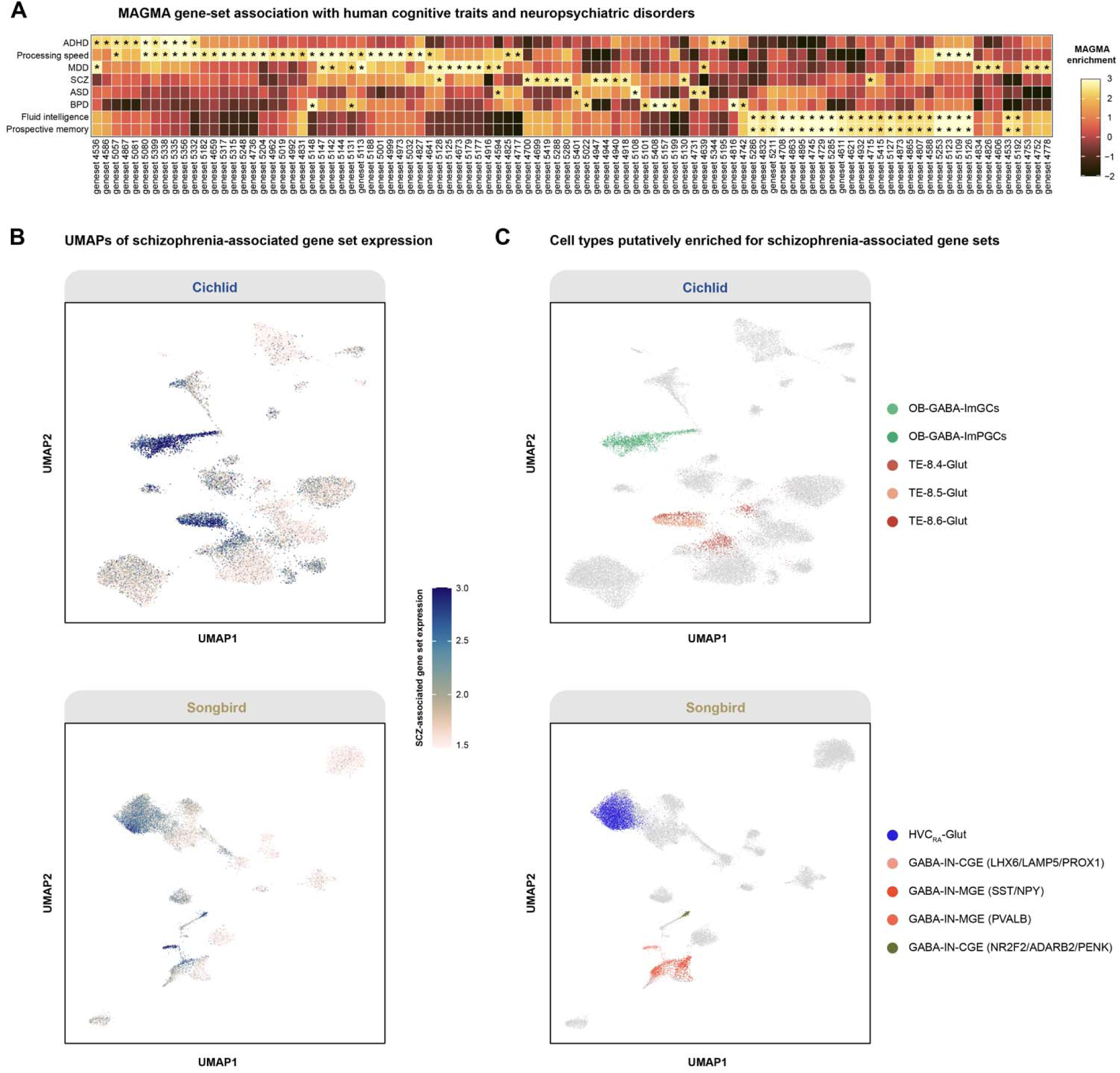
Translational mapping of neuropsychiatric-associated gene sets across vertebrate neuronal populations. (A) MAGMA.Celltyping gene-set enrichment analysis. Heatmap shows enrichment of scGENUS gene sets across neuropsychiatric disorders (ADHD, SCZ, BPD, ASD, MDD) and cognitive traits (fluid intelligence, prospective memory, processing speed). Color scale represents standardized effect size (MAGMA enrichment coefficient / ). Asterisks indicate statistical significance (*p*-value < 0.01) (see Supplementary Data File 6). (B) Projection of SCZ-associated gene-set expression onto single-cell datasets from cichlid and songbird forebrains. Module scores are overlaid on scGENUS-derived UMAP embeddings. (C) scGENUS-derived UMAP embeddings of cichlid and songbird forebrains colored to highlight putative SCZ-associated cell types identified in (B).

Before projecting across species, we first examined the distribution of trait-associated gene-set expression in the human prefrontal cortex dataset. Projection of SCZ-and BPD-associated gene-set expression onto the scGENUS-derived UMAP embedding revealed distinct patterns of enrichment across cell types (Fig. S17B-D). Both SCZ-and BPD-associated gene sets were preferentially enriched in excitatory intratelencephalic neurons (L4 and L5 IT), MGE-derived interneurons (PVALB^+^ and SST^+^), CGE-derived interneurons (LAMP5^+^), and OPCs. SCZ-associated gene sets additionally showed enrichment in astroglial populations, whereas BPD-associated gene sets showed stronger enrichment in L2/3 IT excitatory neurons. These distributions are consistent with prior chromatin accessibility-based analyses demonstrating GWAS signal enrichment across cortical cell types^27^. We next asked whether these programs map onto homologous neuronal populations in non-human species. Because genetic risk for schizophrenia has been linked to neural circuit mechanisms regulating social interaction and vocal communication, we projected SCZ-associated gene sets onto single-cell datasets from cichlid and songbird forebrains, two species in which social and communicative behaviors are mediated by well-characterized neural circuits (Fig. 6B). In cichlid, SCZ-associated gene sets were enriched in olfactory bulb GABAergic granule and periglomerular interneurons, as well as glutamatergic populations, including the 8.4-Glut cluster implicated in social and reproductive behaviors^17^ (Fig. 6C). In songbird, enrichment was observed in MGE-derived (PVALB^+^ and SST^+^) and CGE-derived (LAMP5^+^ and NR2F2^+^) interneurons, as well as HVC projection neurons targeting the robust nucleus of the arcopallium (RA), a key component of the vocal learning circuit that drives song production (Fig. 6C). These findings do not imply conservation of psychiatric disease states. Rather, they demonstrate that human neuropsychiatric risk signals converge on conserved neuronal substrates regulating complex social behaviors across vertebrates.

## DISCUSSION

Cellular diversity evolves through changes in gene expression programs that shape cell identity, functional specialization, and developmental trajectories across lineages. Although comparative single-cell studies increasingly suggest that core cell identities are conserved while their regulatory programs are modified, systematically resolving these conserved and lineage-dependent transcriptional architectures across species remains technically challenging. As a result, how vertebrate forebrain evolution generates and reshapes cellular diversity and cell state transitions through modifications of transcriptional programs remains incompletely understood. scGENUS, an evolutionarily informed comparative framework, enables integration of single-cell transcriptomic data across phylogenetically distant species by capturing complex many-to-many homology relationships through evolutionarily informed gene sets and representing cells in a shared, interpretable gene-set feature space. Applied to forebrain datasets from eleven vertebrate species spanning ∼500 million years of evolution, this framework reveals a common transcriptional organization in which major neuronal, glial, and non-neural cell identities are preserved across lineages. We observe broad conservation of major forebrain cell-type identities alongside substantial lineage-specific modulation of transcriptional programs within this shared cellular framework. Most gene sets show conserved expression patterns across vertebrate forebrain cell types, with conservation generally increasing with evolutionary age and divergence often concentrated at the clade level. We further identify a conserved radial glial bifurcation into neurogenic and gliogenic trajectories that is maintained across vertebrates but exhibits lineage-dependent differences in transcriptional dynamics. Finally, we demonstrate that the gene-set representation enables mapping of human neuropsychiatric genetic risk onto conserved neural cell populations in a cross-species context.

Cross-species single-cell integration across deep phylogenetic distances is fundamentally constrained by complex gene evolutionary histories, including extensive duplication, lineage-specific expansion, and uneven genome annotation, which complicate strategies that rely on strict one-to-one ortholog correspondence. scGENUS addresses this limitation by redefining the unit of comparison from individual genes to evolutionarily informed gene sets derived from a global homology graph constructed from reciprocal protein sequence similarity. By incorporating many-to-many homology relationships during feature construction prior to integration, scGENUS represents cells in a shared gene-set feature space in which gene-set expression profiles define cellular states, preserving a direct and interpretable link between integrated cell types and the homologous genes that compose each gene set. This design confers several advantages. First, it reduces dependence on shared one-to-one ortholog intersections, enabling robust comparison even when gene content differs across lineages. Second, because gene sets are defined from the homology graph rather than inferred implicitly within latent embeddings, gene-set expression can be directly compared across taxa, preserving interpretability and comparability at the level of gene families and transcriptional programs. Third, the gene-set framework supports downstream evolutionary analyses, including differential gene-set expression, developmental trajectory inference, phylogenetic reconstruction, and GWAS enrichment, within a unified and scalable structure. In this way, scGENUS complements existing latent cell-embedding-based integration strategies while restoring direct evolutionary interpretability. Quantitative benchmarking further showed that scGENUS achieves strong overall performance relative to representative cross-species integration approaches while providing a favorable balance between cross-species mixing and preservation of biological structure (Fig. S8; Supplementary Results). scGENUS also significantly outperforms a one-to-one ortholog baseline and negative controls based on size-matched random gene sets and a rewired homology graph, supporting a contribution of evolutionarily informed feature construction beyond generic feature aggregation or batch correction. In our implementation, scGENUS additionally required neither coarse cell-type annotations during integration nor GPU acceleration.

These advantages arise from treating homology as a graph-based feature-construction problem rather than requiring uniquely resolved orthology assignments. Accordingly, scGENUS gene sets represent sequence-homology communities in a multispecies sequence-similarity graph rather than formal orthology groups. Their composition was highly stable across protein length, alignment coverage, alternative edge-weight definitions, stricter BLASTp thresholds, and the retained range of Leiden resolutions, and the resulting gene sets showed significantly greater functional coherence than size-matched random groups (Figs. S1-S5). Mean aggregation across same-species homologs prevents lineage-specific gene expansions from receiving disproportionate feature weight, but may reduce paralog-specific resolution when homologs have diverged in expression, as expected from the established roles of paralog subfunctionalization and neofunctionalization in transcriptome evolution^13,79,80^. Empirically, expression among same-species homologs was generally concordant across cell types, although a minority showed substantial expression-domain divergence (Fig. S19; Supplementary Results). Gene-set-level differences therefore represent aggregate signals rather than evidence of coordinated changes across every constituent homolog. Because constituent gene identities are retained, individual homologs can be traced back and examined in the original gene-level expression data when finer resolution is needed.

The cross-vertebrate atlas reveals a conserved transcriptional organization of the forebrain in which major neuronal, glial, and non-neural cell types align across lineages despite extensive evolutionary divergence. Unsupervised integration across ∼500 million years preserves coherent separation of annotated cell types, indicating that core forebrain identities are anchored by deeply conserved transcriptional programs. Centroid-based transcriptional distance analysis further demonstrates that divergence between cell types exceeds divergence within cell types across species, suggesting that established cell identities constitute relatively stable evolutionary units. Nevertheless, the extent of within-type divergence varies across lineages and cell types. Oligodendrocytes and radial glia exhibit pronounced clade-specific shifts, and teleost lineages show elevated remodeling in select neuronal populations, reflecting lineage-dependent modulation of conserved programs. Transcriptome-derived phylogenies further reinforce this pattern: developmental origins, particularly among GABAergic interneurons, cluster by subtypes rather than by species, in agreement with recent phylogenetic reconstructions that use principal components as evolutionary characters to recover robust cell-type clades across distantly related species^81^. Together, these findings suggest that quantitative modification of conserved cell identities is a prominent feature of vertebrate forebrain evolution, while not excluding lineage-specific cellular innovations that may be underrepresented in a homology-based feature space. Strongly species-biased transcriptional neighborhoods further demonstrate that localized lineage-specific divergence persists within this broadly conserved organization (Fig. S11; Supplementary Results).

The gene-set representation provides a quantitative view of how transcriptional programs are conserved and modified across vertebrates. Most evolutionarily informed gene sets exhibit conserved expression across major cell types, and the degree of conservation scales with inferred evolutionary age, here approximated by the phylogenetic depth of species representation rather than formal reconstruction of gene origin: older gene sets tend to maintain stable cell-type-specific expression profiles, whereas younger or lineage-restricted sets show greater variability. Despite heterogeneity among source datasets, conservation scores remained highly stable in leave-one-species-out analyses, indicating that the overall conservation landscape is robust to exclusion of individual species datasets (Fig. S18; Supplementary Results). Conserved gene sets are enriched for canonical neuronal identity and core cellular functions, while more divergent sets are disproportionately associated with regulatory, modulatory, and structural processes. Notably, divergence often occurs at the level of entire clades rather than being restricted to particular cell types, indicating coordinated remodeling of transcriptional programs within evolutionary lineages.

Radial glia, which generate both neuronal and glial lineages and in some vertebrates remain a principal source of adult neurogenesis, provide a powerful context for examining how conserved developmental programs are modified across evolution. Trajectory inference identifies a conserved bifurcation in which radial glia-like populations diverge into neurogenic and gliogenic trajectories across species, supported by gene-set signatures associated with neurogenic commitment, gliogenic differentiation, and progenitor quiescence. Broad cross-species mixing along both branches supports their interpretation as shared cell-state trajectories rather than species-driven segregation, although species-level pseudotemporal differences may also reflect age, developmental stage, and sampling heterogeneity. Within this conserved fate bifurcation, gene-set dynamics reveal divergence at two levels. First, neurogenic and gliogenic branches are distinguished by differential activation of branch-associated gene sets. Second, within each branch, gene-set expression profiles vary in magnitude and pseudotemporal progression across species. Thus, although the overall bifurcation is conserved, the deployment of branch-specific programs is modulated in a species-dependent manner. These patterns suggest that evolutionary differences in neuronal and glial production arise primarily through quantitative tuning of conserved developmental trajectories.

scGENUS also provides a framework for linking conserved cellular programs to human neuropsychiatric genetics in an evolutionary context. Gene sets enriched for neuropsychiatric GWAS signals map onto conserved neuronal and glial populations, indicating that genetic risk for human brain disorders intersects with deeply conserved cellular substrates. Because these transcriptional programs are shared across vertebrates, cross-species comparisons can reveal how similar cellular programs operate in species with biological capacities absent in humans. For example, axolotls regenerate complex tissues including limbs and spinal cord through injury-induced cellular reprogramming^82^, whereas cichlids maintain lifelong tooth replacement through a persistent successional dental lamina^83^. Mapping conserved transcriptional modules across such species provides a comparative framework for identifying molecular programs associated with regenerative capacity. These evolutionary comparisons therefore create a reciprocal bridge: human neuropsychiatric genetics can help interpret conserved neural substrates of complex social behaviors across vertebrates, while regenerative vertebrate models can reveal how related molecular programs operate in contexts inaccessible in humans.

Although developed here in the context of vertebrate forebrain evolution, scGENUS-derived gene sets may also provide a useful representation for comparative analyses in other biological systems. Because these gene sets are derived from global homology relationships defined across entire species proteomes, rather than from a forebrain-restricted reference, the same representation is not inherently limited to forebrain comparisons and may also be applicable in other biological contexts, where the most informative gene sets will depend on the biology of the organ system under study. In addition, the granularity achievable in these broader applications will ultimately depend on the resolution and comparability of the underlying datasets. Finer cell-subtype resolution can be essential for identifying species differences that may be obscured at broader levels of annotation^84–86^. scGENUS can support such analyses when comparable subtype annotations are available, with finer-scale comparisons resolving conserved subtype-level organization across species (Fig. S20; Supplementary Results). At a complementary level, cell-type identity is shaped and maintained by combinations of transcription factors acting at cis-regulatory elements. As single-cell multiomic datasets become increasingly available, this framework could also be extended to open chromatin states, offering clearer resolution of cell-type specificity and the upstream regulatory basis of gene-set expression changes. In this context, chromatin accessibility can be aggregated into gene-set activity, extending the shared gene-set feature space from transcriptomic to regulatory variation. Evolutionarily informed gene-set representations may thus support direct comparative analysis of cis-regulatory programs, providing a basis for understanding how conserved cellular programs are differentially regulated across species.

## ACKNOWLEDGEMENTS

We thank Dr. George W. Gruenhagen for thoughtful comments on early drafts of the manuscript. We also thank Caitlin M. Sullivan and Siddhi Vivekkumar Ranbhor for valuable discussions during the early stages of the project. This work was supported by NIH R01GM144560 and R35GM162583 to J.T.S.

## CONTRIBUTIONS

P.Q. and H.H. conceived the study and developed the methodology. H.H. performed the analyses and generated the visualizations. H.H. wrote the initial draft of the manuscript. P.Q. and J.T.S. reviewed and edited the manuscript. P.Q. and J.T.S. supervised the project.

## METHODS

### Overview of the scGENUS framework

To enable comparative analysis of single-cell transcriptomes across the eleven phylogenetically distant vertebrate species analyzed in this study, we developed the scGENUS framework, a graph-based approach that transforms gene-level expression profiles into an evolutionarily informed gene-set feature space. The framework operates on multispecies single-cell expression matrices together with corresponding protein sequences and gene annotations derived from reference genomes.

Let

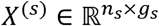

denote the gene expression matrix for species *s*, where *n_s_* represents the number of cells and *g_s_* represents the number of genes in that species. The set of input expression matrices is therefore

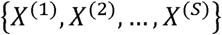

where *S* = 11 corresponds to the eleven vertebrate species included in this study. The scGENUS workflow consists of three sequential steps:

1. Construction of a global gene homology graph across species
2. Identification of homologous gene communities (gene sets)
3. Transformation of gene expression matrices into gene-set expression matrices

### Construction of the global gene homology graph

Pairwise protein sequence alignments were performed between all species pairs using BLASTp^87^. Protein sequences were obtained from NCBI RefSeq proteomes, and protein identifiers were mapped to their corresponding gene identifiers using genome annotations prior to graph construction. For each pair of proteins from different species, BLASTp searches were performed in both directions. When a pair of proteins shared multiple High Scoring Pairs (HSPs), which represent local regions of sequence similarity, the HSP with the highest bit score was used to represent the similarity between those proteins. Only alignments with E-value <10^-6^ were retained, while all other BLAST parameters were left at their default settings.

Based on these alignments, we constructed an undirected weighted multipartite graph

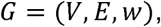

where nodes in *V* represent genes, edges *E* represent homology relationships between genes across species, and the weight function *w* quantifies sequence similarity between homologous genes. The node set

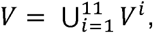

is the union of genes from the eleven species, where *V^i^* denotes the set of genes from species *i*. Edges connect genes whose encoded proteins exhibit sequence similarity in BLASTp alignments. Formally, edges satisfy

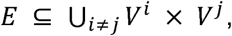

where *i* and *j* denote genes belonging to different species. To reduce weak or asymmetric matches, only reciprocal alignments were retained. Edge weights quantify sequence similarity between homologous genes and were defined as

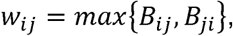

where *B_ij_* and *B_ij_* denote BLASTp bit scores obtained from alignments identified in reciprocal directions between genes *i* and *j*. The maximum reciprocal bit score was used as a pragmatic symmetric edge weight that retains the strongest sequence-similarity evidence observed for a reciprocal gene pair after protein-level alignments were mapped to gene identifiers. This choice was motivated in part by the reciprocal BLAST graph formulation used in SAMap^13^, which similarly retains reciprocal sequence-similarity relationships and summarizes multiple local alignments using the highest-scoring HSP. Sensitivity of gene-set construction to alternative reciprocal edge-weight definitions was evaluated separately (Fig. S2). This graph formulation naturally accommodates many-to-many homology relationships arising from gene duplication and divergence during vertebrate evolution.

### Identification of homologous gene sets

To identify groups of homologous genes, the global gene homology graph *G* = (*V, E, w*) was partitioned using the Leiden community detection algorithm^88^. Community detection was applied directly to the weighted homology graph. In this graph, nodes correspond to genes and edge weights reflect sequence similarity derived from reciprocal BLASTp alignments. The Leiden algorithm partitions the graph into densely connected subgraphs by optimizing a modularity-based objective function. In this context, each detected community represents a group of genes that share stronger sequence similarity relationships with one another than with genes outside the community, thereby identifying homologous gene groups across species.

Because homology relationships span multiple evolutionary scales, community detection was performed across a range of resolution parameters to capture gene groups at different evolutionary scales. Lower resolution values identify larger gene communities corresponding to broadly conserved gene families, whereas higher resolution values partition these families into smaller subgroups reflecting more specific evolutionary relationships. Resolution parameters ranged from 0.01 to 100, sampled approximately on a logarithmic scale.

To ensure sufficient cross-species representation, resolutions for which fewer than 50% of gene sets contained genes from all eleven species were discarded. Redundant gene sets identified across retained resolutions were then collapsed. The resulting communities were defined as evolutionarily informed gene sets, which serve as the fundamental features for downstream analyses. Each gene set may contain multiple genes from the same species as well as homologs from multiple species, reflecting gene duplication and lineage-specific expansion. These gene sets provide the basis for transforming gene-level expression matrices into a shared gene-set feature space for cross-species integration.

Robustness of gene-set construction to protein alignment coverage, BLASTp E-value threshold, reciprocal edge-weight definition, Leiden resolution, and cross-species filtering criteria was evaluated systematically (Figs. S1-S4), and the structural and functional characteristics of the final gene-set collection were additionally characterized (Fig. S5).

### Transformation of gene-level expression to gene-set expression

Following identification of homologous gene sets, gene-level expression matrices were transformed into gene-set expression matrices to enable cross-species comparison in a shared feature space. For each species *s*, let

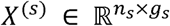

denote the gene expression matrix, where *n_s_* is the number of cells and *g_s_* is the number of genes for that species. Let *G_j_* denote the set of genes belonging to gene set *j*, as identified from the homology graph partitioning. For each cell *c* and gene set *j*, gene-set expression values were computed as the mean expression of all genes in *G_j_*:

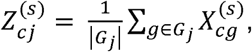

where 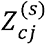 represents the expression value of gene set *j* in cell *c*, and |*G_j_*| denotes the number of genes contained in gene set *j*. If a gene set contained genes that were absent from the expression matrix of species *s*, aggregation was performed using only the subset of genes from *G_j_* present in that species. Mean aggregation was used so that gene sets containing lineage-specific gene expansions did not receive greater feature weight solely because a species contained more homologous copies. This strategy can, however, attenuate paralog-specific expression differences when same-species homologs have diverged in expression. We therefore separately evaluated expression concordance among same-species homologs within scGENUS gene sets (Fig. S19).

This transformation maps gene-level expression profiles into a shared gene-set feature space, where each feature corresponds to a homologous gene group identified from the global homology graph. Because gene sets may contain multiple homologs from the same species as well as homologs across species, this representation accommodates lineage-specific gene duplication and differential gene content while preserving the structure of conserved transcriptional programs. The resulting gene-set expression matrices

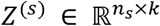

(where *k* denotes the total number of gene sets) were used for all downstream analyses, including cross-species integration, dimensionality reduction, cell-type compositional analysis, differential gene-set expression testing, developmental trajectory inference, and comparative evolutionary analyses. The mapping between each gene set and its constituent genes was retained, allowing gene-set-level signals to be traced back to individual homologs in the original gene-level expression matrices when finer interpretation was required.

### Single-cell transcriptomic dataset retrieval

Single-cell and single-nucleus RNA sequencing datasets from eleven vertebrate species were assembled from previously published studies. Datasets for cichlid (*Mchenga conophorus*^17^), goldfish (*Carassius auratus*^19^), axolotl (*Ambystoma mexicanum*^21^), turtle (*Chrysemys picta bellii*^23^), songbird (*Taeniopygia guttata*^24^) and mouse (*Mus musculus*^26^) were obtained from a published cross-species forebrain atlas^18^. For these datasets, preprocessing and quality-control procedures followed those described in the original study. Briefly, cells were restricted to telencephalic or telencephalon-equivalent forebrain regions as defined by the source annotations, and quality filtering was performed according to the criteria reported in the corresponding publications. Additional datasets were collected for sea lamprey (*Petromyzon marinus*^16^), zebrafish (*Danio rerio*^20^), human (*Homo sapiens*^27^), lizard (*Pogona vitticeps*^22^) and chicken (*Gallus gallus*^25^). For datasets generated as whole-brain atlases (sea lamprey and lizard), only cells annotated in the original studies as belonging to the telencephalon were retained. The human dataset comprised both prefrontal cortex (PFC) and primary visual cortex; only PFC cells were included in the present analysis. To better align developmental stages across species, the human dataset was further restricted to postnatal samples. For the lizard dataset, telencephalic cell-type annotations were available only for neuronal populations; glial and other non-neural cell types were therefore excluded. Where available, neuronal cell annotations were further refined by retaining cell types with clear anatomical annotations for glutamatergic neurons and developmental origin annotations for GABAergic interneurons as reported in the original datasets or accompanying supplementary materials. All gene-level expression matrices were normalized by library size using default parameters prior to downstream processing. These normalized gene-level matrices were subsequently transformed into gene-set expression matrices for cross-species analysis. Detailed dataset-and sample-level metadata, including source study, sequencing modality and platform/chemistry, biological replicates, anatomical region, developmental stage, sex where available, retained cell numbers, and major cell-type composition, are provided in Supplementary Data File 7.

### Retrieval of protein sequences and genome annotations

To construct the gene homology graph and quantify sequence similarity across species, we retrieved reference proteomes and genome annotations for each species included in the analysis. Protein sequences were downloaded from NCBI RefSeq using the genome assemblies referenced in the corresponding single-cell studies when available. These included *Maylandia zebra* (GCF_000238955.4), *Carassius auratus* (GCF_003368295.1), *Danio rerio* (GCF_000002035.6), *Mus musculus* (GCF_000001635.27), *Homo sapiens* (GCF_000001405.40), *Pogona vitticeps* (GCF_900067755.1), *Chrysemys picta bellii* (GCF_000241765.3), *Taeniopygia guttata* (GCF_003957565.1) and *Gallus gallus* (GCF_000002315.5). For *Petromyzon marinus*, protein sequences and gene annotations were obtained from the custom genome annotation provided in the original study and made available through Zenodo^89^. For *Ambystoma mexicanum*, the proteome and gene annotations were obtained from the v6.0 axolotl genome assembly (AmexT_v47^90^), which is not available through the NCBI RefSeq repository. Protein identifiers were mapped to gene identifiers using the corresponding genome annotation files prior to homology graph construction.

### Cross-species integration and dimensionality reduction in gene-set feature space

Gene-set expression matrices derived from each species were concatenated and stored as a Seurat^91^ object for cross-species analysis. Mitochondrial-and ribosomal-associated gene sets were retained in the full scGENUS feature catalog rather than being explicitly removed prior to integration or downstream analyses; their potential influence was evaluated separately (Fig. S7). Data normalization was performed using SCTransform^92^, which models gene-set expression values using a regularized negative binomial regression model while correcting for technical confounding factors such as sequencing depth. Principal component analysis (PCA) was performed on the concatenated gene-set expression matrix to obtain a low-dimensional representation of cellular transcriptional programs. The top 50 principal components were retained for downstream integration. To account for dataset-specific technical variation arising from different sequencing platforms, experimental batches, and sampling differences across studies, batch correction was performed using Harmony^93^. Harmony was applied to the principal component representation of the combined gene-set expression matrix using the batch annotations provided in the original datasets as covariates. This procedure generated an integrated embedding that preserves shared biological structure while reducing batch-associated variation.

For visualization, two-dimensional embeddings were computed using Uniform Manifold Approximation and Projection (UMAP) applied to the Harmony-corrected principal components. These embeddings were used to visualize cross-species relationships among cell populations in the shared gene-set feature space. UMAP parameters were set as follows: the minimum distance between embedded points was set to 0.1 (min.dist = 0.1), the number of neighboring points used in local approximations was set to 50 (n.neighbors = 50), and cosine distance was used as the similarity metric. Cell-type annotations used for coloring UMAP visualizations were taken from the original studies and retained for downstream comparative analyses.

### Assessment of dataset heterogeneity and integration robustness

To assess the impact of dataset heterogeneity, within-species batch mixing was quantified using graph iLISI before and after Harmony correction, comparing the pre-Harmony PCA and Harmony-corrected PCA representations for datasets containing multiple batches. Robustness of gene-set conservation estimates to individual species datasets was additionally evaluated using a leave-one-species-out analysis. Conservation scores were recalculated after excluding each species in turn and compared with scores obtained from the complete eleven-species dataset (Fig. S18).

### Benchmarking of cross-species integration methods

Cross-species integration performance was benchmarked against SATURN, CAMEX, SAMap, and a one-to-one ortholog representation integrated using Harmony (Harmony O2O). Where required by the respective implementation, coarse cell-type annotations were provided during model training. Two negative controls were additionally evaluated: size-matched random gene sets and gene sets derived from a rewired homology graph, each processed using the same downstream integration workflow as scGENUS. Integration performance was evaluated using metrics adapted from the scIB framework. Batch-correction performance was quantified using batch silhouette score, graph iLISI, and graph connectivity, whereas biological conservation was evaluated using cell-type silhouette score, isolated-label preservation, and graph cLISI. Metrics were oriented such that higher values indicate better performance and were summarized into batch-correction, biological-conservation, and overall aggregate scores. Detailed benchmark results are provided in Fig. S8 and Supplementary Data File 8.

### Cell-type compositional analysis across species

To quantify differences in cell-type composition across species, we performed neighborhood-level compositional analysis using Milo^37^. A k-nearest neighbor (KNN) graph was constructed in the Harmony-corrected principal component representation of the gene-set expression space using the top 50 principal components with 50 nearest neighbors. Overlapping cellular neighborhoods were then defined using the makeNhoods function in Milo, which samples representative cells and constructs neighborhoods based on their local connectivity in the KNN graph. For each neighborhood, the distribution of cells originating from each species was quantified. To account for unequal representation of species across the dataset and prevent datasets with larger cell numbers from biasing compositional estimates, species counts within each neighborhood were normalized relative to global species frequencies across the dataset, yielding weighted species enrichment values. Species composition within each neighborhood was visualized as pie charts in the neighborhood graph representation. To quantify the degree of species mixing within each neighborhood, Shannon entropy of species labels was calculated from the normalized species proportions. Entropy values range from 0 (single-species dominance) to 1 (maximal cross-species mixing). Neighborhoods were further grouped using k-means clustering (k = 20) applied to the neighborhood graph representation. For each cluster of neighborhoods, the median entropy was calculated to summarize the degree of local species mixing. Clusters with median entropy values below 0.55 were interpreted as exhibiting species-enriched cellular composition and were highlighted in downstream analyses. To separately characterize the strongest species-biased neighborhoods at the individual neighborhood level, we examined the distribution of normalized species-mixing entropy and used the local minimum separating the low-entropy mode from the broader distribution (*h* < 0.29) as a stringent threshold. The prevalence and cellular composition of these neighborhoods were summarized across species and major cell types, and their neighborhood sizes were compared with those of the remaining neighborhoods (Fig. S11).

### Differential gene-set expression analysis

To identify marker gene sets for each major cell type, differential gene-set expression analysis was performed using the FindAllMarkers function implemented in Seurat. Tests were conducted on the gene-set expression matrices derived from the scGENUS framework. Only positive markers were retained (only.pos = TRUE). A minimum log_2_ fold change threshold of 0.25 (logfc.threshold = 0.25) and a minimum detection rate of 5% of cells (min.pct = 0.05) were required to ensure that marker gene sets were robustly detected in the corresponding cell populations. Statistical significance was evaluated using the Wilcoxon rank-sum test, and gene sets with Bonferroni-adjusted *p*-values < 0.05 were retained as cell-type-specific markers.

To identify gene sets exhibiting species-specific expression patterns, differential expression analysis was performed within each annotated cell type. Gene-set expression values were compared between species using the FindMarkers function in Seurat, where each species was compared against all remaining species within the same cell type. Statistical testing was performed using the Wilcoxon rank-sum test, and multiple-testing correction was applied using the Bonferroni method implemented in Seurat. Gene sets with absolute log_2_ fold change greater than 0.25 and adjusted *p*-values < 0.05 were considered significantly differentially expressed.

### PCA distance-based comparative analysis of cell types

To quantify evolutionary divergence of transcriptional states across species, we performed a centroid-based distance analysis following approaches previously described by Chari *et al*^94^. and Kaplan *et al*^95^. For each cell type, cells from all species were subsetted and normalized using SCTransform, after which 2,000 highly variable gene sets were identified and principal component analysis (PCA) was performed. For each species within a given cell type, a centroid vector was calculated as the mean position of all cells belonging to that species in the PCA-reduced space. Evolutionary divergence within each cell type was quantified by computing pairwise L1 distances between species-specific centroids, representing transcriptional differences between species within the same cell type. The number of principal components used for each cell type was determined by identifying the minimum number of leading principal components required to explain 20% of the variance in the data. If more than 100 principal components were required to reach this threshold, the analysis was limited to the top 100 principal components. Distances calculated between centroids of the same cell type across different species represent evolutionary divergence of transcriptional states within that cell type. For comparison, we also calculated distances between centroids of different cell types using the Harmony-integrated PCA space, providing a reference scale for divergence between distinct cell identities.

### Phylogenetic reconstruction of cell-type relationships

To examine the evolutionary organization of vertebrate forebrain cell types, we reconstructed distance-based phylogenetic trees from transcriptomic similarity in the shared Harmony-corrected principal component space derived from scGENUS gene-set expression. Analyses were performed separately for the four major cell classes, GABAergic neurons, glutamatergic neurons, glial cells, and non-neural cells, with phylogenies inferred from the annotated subtypes within each class. For each annotated cell subtype, a centroid vector was calculated as the mean position of all cells belonging to that subtype in the first 50 principal components.

Pairwise similarity between cell-type centroids was computed using Pearson correlation, and distances were defined as 1 − *ρ*, where *ρ* represents the Pearson correlation coefficient between centroid vectors. An unrooted neighbor-joining tree was then inferred from the resulting distance matrix using the *ape*^96^ package in R. To assess the robustness of tree topology, we performed bootstrap resampling of cells within each cell subtype. In each bootstrap iteration, cells belonging to each subtype were sampled with replacement, subtype centroids were recalculated, and a new correlation-based distance matrix was computed. A neighbor-joining tree was then reconstructed from this bootstrap distance matrix. This procedure was repeated for 1,000 bootstrap iterations, generating a distribution of bootstrap trees representing sampling variation in the data. Tip stability was quantified using the leaf stability index (LSI) implemented in the *Rogue*^97^ package in R. LSI was computed from the set of bootstrap trees and measures how consistently each cell-type tip occupies a similar phylogenetic position across bootstrap replicates. LSI values range from 0 (low stability) to 1 (high stability). Phylogenetic trees were visualized with branch lengths representing transcriptomic divergence between cell-type centroids.

### Gene-set expression conservation analysis

To quantify the degree of cross-species conservation in gene-set expression programs, we computed pairwise Spearman correlations of gene-set expression profiles across species. For each species, gene-set expression values were aggregated into pseudobulk profiles across the ten major cell types by averaging gene-set expression across all cells belonging to each cell type. For each gene set, this procedure produced a species-specific expression vector describing its relative expression across the ten cell types. Pairwise Spearman correlation coefficients were then calculated between all species pairs using these vectors. A gene-set expression conservation score was defined as the mean of the pairwise Spearman correlation coefficients across species comparisons, yielding a score ranging from -1 (highly divergent expression) to 1 (highly conserved expression). To investigate the relationship between expression conservation and evolutionary origin, we assigned an evolutionary age to each gene set based on the most recent common ancestor (LCA) of the species represented within that set. Phylogenetic relationships among species were represented using a reference vertebrate species tree containing the eleven species included in this study, and LCA assignments were inferred using the getMRCA function implemented in the *ape*^96^ package in R.

### Paralog expression concordance analysis

For each gene-set–species combination containing at least two same-species homologs, gene-level expression profiles were summarized across cell-type pseudobulks and pairwise Spearman correlations were calculated among homologs. The median pairwise correlation was used to summarize expression concordance for each gene-set–species combination. For low-concordance cases, expression-domain divergence was quantified as *dT* = 1 - *N_both_*/*N_either_*, where *N_both_* denotes the number of cell types in which both homologs were expressed and *N_either_* is the number in which at least one paralog is expressed. For discordant cases with an appropriate single-copy outgroup, the closest phylogenetically relevant single-copy homolog was used as a proxy for the ancestral expression domain to identify putative domain partitioning, loss, or gain.

### Learning vector fields using the CellRank2 CytoTRACE kernel

To estimate developmental potential and infer directed transitions between cellular states, we applied the CytoTRACE kernel implemented in CellRank2^55^. This approach estimates cellular plasticity directly from transcriptional diversity and does not require specification of a predefined root cell for trajectory inference. CytoTRACE scores were computed for each cell from the scGENUS-derived gene-set expression matrix using the compute_cytotrace function in the CytoTRACE Kernel. A k-nearest neighbor (KNN) graph capturing transcriptomic similarity between cells was constructed and combined with CytoTRACE scores to compute a directed cell-cell transition matrix. The resulting transition probabilities were projected onto the low-dimensional embedding to visualize differentiation dynamics as vector fields.

### Trajectory inference and pseudotime analysis

To gain additional insights into gene-set expression variations during cell differentiation, we performed pseudotime analysis using Monocle3^64^ to reconstruct single-cell trajectories. The normalized gene-set expression matrix, cell-type annotations, and low-dimensional embeddings from the Seurat object were imported into Monocle3 using the as.cell_data_set function. Trajectories were inferred using the learn_graph function, which fits a principal graph on the UMAP embedding to recover differentiation paths. Cells were then ordered along the trajectory using the order_cells function, with root nodes selected based on developmental potential estimated from the CellRank2 CytoTRACE kernel. Analysis of specific trajectory branches was performed using the choose_cells function. Within each branch, gene sets exhibiting dynamic expression along pseudotime were identified using the graph_test function, which evaluates gene-set expression autocorrelation along the trajectory using the Moran’s I statistic. To identify gene sets significantly associated with pseudotime, we filtered the output of the graph_test function using a threshold of *q*-value < 0.01, and further refined the selection by retaining the top 500 gene sets ranked by Moran’s I. Pseudotime-dependent gene-set expression patterns were clustered using k-means and visualized with the ClusterGVis package (v0.1.2) in R.

### State-dependent regression analysis of pseudotime dynamics

To test for state-dependent differences in gene-set expression dynamics between neurogenic and gliogenic branches, we applied a natural spline linear regression model along pseudotime as previously described by Kanton *et al*^65^. For each gene set, expression values were modeled as a function of pseudotime using a natural spline regression (degrees of freedom = 6). A null model was first fitted assuming a shared expression trajectory across both branches. An alternative model allowing branch-specific regression curves was then fitted. The residual variance of the two models was compared using an F-test to determine whether gene-set expression dynamics differed significantly between the neurogenic and gliogenic branches. *p*-values were adjusted for multiple testing using the Bonferroni correction, and gene sets with adjusted *p*-values < 0.0001 were considered significantly state-dependent.

### Species-specific regression analysis of pseudotime dynamics

To test for species-specific differences in gene-set expression dynamics along pseudotime, we applied a generalized linear regression model implemented in Monocle3^64^ using the fit_models function. Gene-set expression was fitted with the formula ∼species*pseudotime, where the interaction term captures species-specific differences in expression dynamics along the trajectory. *p*-values were adjusted using the Bonferroni correction, and gene sets with adjusted *p*-values < 0.0001 were considered significant.

### MAGMA gene-set enrichment analysis

To assess enrichment of GWAS signals in scGENUS-derived gene sets, we performed gene-set enrichment analysis using the MAGMA.Celltyping^98^ R package, which implements the MAGMA^99^ framework. SNPs from GWAS summary statistics were first mapped to genes using the map_snps_to_genes function, assigning SNPs to gene regions defined as 35 kb upstream and 10 kb downstream of each gene. Gene-set enrichment was subsequently evaluated using MAGMA’s linear regression model through the calculate_geneset_enrichment function, which tests whether genes belonging to a given gene set show stronger GWAS associations than genes outside the set. For each scGENUS gene set, human orthologs were used as representative genes, as GWAS summary statistics are derived from human populations. Gene sets with *p*-values < 0.01 were considered significantly associated with the trait. The strength of GWAS enrichment was quantified using the standardized effect size (*β*/SE). The GWAS data used in this study can be downloaded from the following links:

- fluid intelligence^100^ (phenocode 20016), processing speed^100^ (phenocode 20023) and prospective memory^100^ (phenocode 20018): https://pan.ukbb.broadinstitute.org/downloads/;
- ASD^101^: https://figshare.com/articles/dataset/asd2019/14671989;
- MDD^102^: https://datashare.ed.ac.uk/handle/10283/3203;
- BPD^103^: https://figshare.com/articles/dataset/bip2021_noUKBB/22564402;
- ADHD^104^: https://figshare.com/articles/dataset/adhd2022/22564390;
- SCZ^105^: https://figshare.com/articles/dataset/cdg2018-bip-scz/14672019.

### Gene ontology enrichment analysis

Gene ontology (GO) enrichment analysis was performed to characterize the functional annotations of genes within each gene set. For each scGENUS gene set, human orthologs were used for GO annotations, as GO databases are primarily curated for human genes. Over-representation analysis was conducted using the genORA function implemented in the *GeneKitr*^106^ package. This method tests whether genes within a given set are significantly overrepresented in predefined GO categories (biological processes) relative to the genomic background. Enrichment terms with Benjamini-Hochberg adjusted q-value < 0.01 were considered statistically significant. To reduce redundancy among enriched GO terms, semantic similarity analysis was applied to cluster related GO categories and summarize representative functional annotations.

## DATA AVAILABILITY

All data needed to evaluate the conclusions in this paper are present in the paper and/or the Supplementary Materials. A GitHub repository (https://github.com/Haowen-He/scGENUS) is provided containing select input, intermediate and output files (‘scGENUS/Supplementary-data/’) sufficient to reproduce the analyses.

## CODE ABAILABILTY

All custom scripts for data analyses are available in our GitHub repository (https://github.com/Haowen-He/scGENUS) and Zenodo (https://doi.org/10.5281/zenodo.19198180).

## REFERENCES

1. O’Connell, L. A. & Hofmann, H. A. Evolution of a Vertebrate Social Decision-Making Network. Science 336, 1154–1157 (2012).

2. O’Connell, L. A. & Hofmann, H. A. The Vertebrate mesolimbic reward system and social behavior network: A comparative synthesis. J. Comp. Neurol. 519, 3599–3639 (2011).

3. Briscoe, S. D. & Ragsdale, C. W. Homology, neocortex, and the evolution of developmental mechanisms. Science 362, 190–193 (2018).

4. Wullimann, M. F. & Mueller, T. Teleostean and mammalian forebrains contrasted: Evidence from genes to behavior. J. Comp. Neurol. 475, 143–162 (2004).

5. Pessoa, L., Medina, L., Hof, P. R. & Desfilis, E. Neural architecture of the vertebrate brain: implications for the interaction between emotion and cognition. Neurosci. Biobehav. Rev. 107, 296–312 (2019).

6. Cajal, S. R. Y., Azoulay, D. L., Swanson, N. & Swanson, L. W. Histology Of The Nervous System: Of Man And Vertebrates. (Oxford University Press New York, NY, 1995). doi:10.1093/oso/9780195074017.001.0001.

7. Dugas-Ford, J., Rowell, J. J. & Ragsdale, C. W. Cell-type homologies and the origins of the neocortex. Proc. Natl. Acad. Sci. 109, 16974–16979 (2012).

8. Fu-Bao-Qian, H. et al. Cross-species single-cell transcriptomics reveals neuronal similarities and heterogeneity in amniote pallium: *Zool*. Res. 46, 193–208 (2025).

9. Zaremba, B. et al. Developmental origins and evolution of pallial cell types and structures in birds. Science 387, eadp5182 (2025).

10. Weissman, T. Neurogenic Radial Glial Cells in Reptile, Rodent and Human: from Mitosis to Migration. Cereb. Cortex 13, 550–559 (2003).

11. Nehrt, N. L., Clark, W. T., Radivojac, P. & Hahn, M. W. Testing the Ortholog Conjecture with Comparative Functional Genomic Data from Mammals. PLoS Comput. Biol. 7, e1002073 (2011).

12. Prince, V. E. & Pickett, F. B. Splitting pairs: the diverging fates of duplicated genes. Nat. Rev. Genet. 3, 827–837 (2002).

13. Tarashansky, A. J. et al. Mapping single-cell atlases throughout Metazoa unravels cell type evolution. eLife 10, e66747 (2021).

14. Rosen, Y. et al. Toward universal cell embeddings: integrating single-cell RNA-seq datasets across species with SATURN. Nat. Methods 21, 1492–1500 (2024).

15. Liu, X., Shen, Q. & Zhang, S. Cross-species cell-type assignment from single-cell RNA-seq data by a heterogeneous graph neural network. Genome Res. 33, 96–111 (2023).

16. Lamanna, F. et al. A lamprey neural cell type atlas illuminates the origins of the vertebrate brain. *Nat*. Ecol. Evol. 7, 1714–1728 (2023).

17. Johnson, Z. V. et al. Cellular profiling of a recently-evolved social behavior in cichlid fishes. Nat. Commun. 14, 4891 (2023).

18. Hegarty, B. E., Gruenhagen, G. W., Johnson, Z. V., Baker, C. M. & Streelman, J. T. Spatially resolved cell atlas of the teleost telencephalon and deep homology of the vertebrate forebrain. *Commun*. Biol. 7, 612 (2024).

19. Tibi, M. et al. A telencephalon cell type atlas for goldfish reveals diversity in the evolution of spatial structure and cell types. Sci. Adv. 9, eadh7693 (2023).

20. Anneser, L., Satou, C., Hotz, H.-R. & Friedrich, R. W. Molecular organization of neuronal cell types and neuromodulatory systems in the zebrafish telencephalon. Curr. Biol. 34, 298–312.e4 (2024).

21. Lust, K. et al. Single-cell analyses of axolotl telencephalon organization, neurogenesis, and regeneration. Science 377, eabp9262 (2022).

22. Hain, D. et al. Molecular diversity and evolution of neuron types in the amniote brain. Science 377, eabp8202 (2022).

23. Tosches, M. A. et al. Evolution of pallium, hippocampus, and cortical cell types revealed by single-cell transcriptomics in reptiles. Science 360, 881–888 (2018).

24. Colquitt, B. M., Merullo, D. P., Konopka, G., Roberts, T. F. & Brainard, M. S. Cellular transcriptomics reveals evolutionary identities of songbird vocal circuits. Science 371, eabd9704 (2021).

25. Hecker, N. et al. Enhancer-driven cell type comparison reveals similarities between the mammalian and bird pallium. Science 387, eadp3957 (2025).

26. Zeisel, A. et al. Molecular Architecture of the Mouse Nervous System. Cell 174, 999–1014.e22 (2018).

27. Wang, L. et al. Molecular and cellular dynamics of the developing human neocortex. Nature 647, 169– 178 (2025).

28. Maroteaux, L., Campanelli, J. & Scheller, R. Synuclein: a neuron-specific protein localized to the nucleus and presynaptic nerve terminal. J. Neurosci. 8, 2804–2815 (1988).

29. Ghiglieri, V., Calabrese, V. & Calabresi, P. Alpha-Synuclein: From Early Synaptic Dysfunction to Neurodegeneration. Front. Neurol. 9, 295 (2018).

30. Suzuki, C. et al. Direct evidence for ultrastructures of the α-synuclein-associated synaptic vesicle pool in presynaptic terminals. Biochim. Biophys. Acta BBA -Mol. Basis Dis. 1870, 167494 (2024).

31. Inoue, T., Ota, M., Ogawa, M., Mikoshiba, K. & Aruga, J. *Zic1* and *Zic3* Regulate Medial Forebrain Development through Expansion of Neuronal Progenitors. J. Neurosci. 27, 5461–5473 (2007).

32. Gheldof, A., Hulpiau, P., Van Roy, F., De Craene, B. & Berx, G. Evolutionary functional analysis and molecular regulation of the ZEB transcription factors. Cell. Mol. Life Sci. 69, 2527–2541 (2012).

33. Leszczyński, P. et al. Emerging Roles of PRDM Factors in Stem Cells and Neuronal System: Cofactor Dependent Regulation of PRDM3/16 and FOG1/2 (Novel PRDM Factors). Cells 9, 2603 (2020).

34. Du, H. et al. Transcription factors *Bcl11a* and *Bcl11b* are required for the production and differentiation of cortical projection neurons. Cereb. Cortex 32, 3611–3632 (2022).

35. Seigfried, F. A. & Britsch, S. The Role of Bcl11 Transcription Factors in Neurodevelopmental Disorders. Biology 13, 126 (2024).

36. Aruga, J. et al. An oligodendrocyte enhancer in a phylogenetically conserved intron region of the mammalian myelin gene *Opalin*. J. Neurochem. 102, 1533–1547 (2007).

37. Dann, E., Henderson, N. C., Teichmann, S. A., Morgan, M. D. & Marioni, J. C. Differential abundance testing on single-cell data using k-nearest neighbor graphs. Nat. Biotechnol. 40, 245–253 (2022).

38. Pachicano, M. et al. Laminar organization of pyramidal neuron cell types defines distinct CA1 hippocampal subregions. Nat. Commun. 16, 10604 (2025).

39. Danielson, N. B. et al. Sublayer-Specific Coding Dynamics during Spatial Navigation and Learning in Hippocampal Area CA1. Neuron 91, 652–665 (2016).

40. Arendt, D. et al. The origin and evolution of cell types. Nat. Rev. Genet. 17, 744–757 (2016).

41. Arendt, D., Bertucci, P. Y., Achim, K. & Musser, J. M. Evolution of neuronal types and families. Curr. Opin. Neurobiol. 56, 144–152 (2019).

42. Wang, J. et al. Tracing cell-type evolution by cross-species comparison of cell atlases. Cell Rep. 34, 108803 (2021).

43. Schweitzer, J., Becker, T., Schachner, M., Nave, K.-A. & Werner, H. Evolution of myelin proteolipid proteins: Gene duplication in teleosts and expression pattern divergence. Mol. Cell. Neurosci. 31, 161– 177 (2006).

44. Jeserich, G., Klempahn, K. & Pfeiffer, M. Features and Functions of Oligodendrocytes and Myelin Proteins of Lower Vertebrate Species. J. Mol.Neurosci. 35, 117–126 (2008).

45. Hu, H., Gao, T., Zhao, J. & Li, H. Oligodendrogenesis in Evolution, Development and Adulthood. Glia 73, 1770–1783 (2025).

46. Fjodorova, M., Noakes, Z. & Li, M. How to make striatal projection neurons. Neurogenesis 2, e1100227 (2015).

47. Van Velthoven, C. T. J. et al. Transcriptomic and spatial organization of telencephalic GABAergic neurons. Nature 647, 143–156 (2025).

48. Krienen, F. M. et al. Innovations present in the primate interneuron repertoire. Nature 586, 262–269 (2020).

49. Garg, V. et al. Mature interneuron subtypes arise from distinct spatial and temporal subdomains within the caudal ganglionic eminence. Preprint at 10.1101/2025.07.28.667082 (2025).

50. Flandin, P. et al. Lhx6 and Lhx8 Coordinately Induce Neuronal Expression of Shh that Controls the Generation of Interneuron Progenitors. Neuron 70, 939–950 (2011).

51. Arellano, J. I., Morozov, Y. M., Micali, N. & Rakic, P. Radial Glial Cells: New Views on Old Questions. Neurochem. Res. 46, 2512–2524 (2021).

52. Ganz, J. & Brand, M. Adult Neurogenesis in Fish. Cold Spring Harb. Perspect. Biol. 8, a019018 (2016).

53. Jurisch-Yaksi, N., Yaksi, E. & Kizil, C. Radial glia in the zebrafish brain: Functional, structural, and physiological comparison with the mammalian glia. Glia 68, 2451–2470 (2020).

54. Trapnell, C. et al. The dynamics and regulators of cell fate decisions are revealed by pseudotemporal ordering of single cells. Nat. Biotechnol. 32, 381–386 (2014).

55. Weiler, P., Lange, M., Klein, M., Pe’er, D. & Theis, F. CellRank 2: unified fate mapping in multiview single-cell data. Nat. Methods 21, 1196–1205 (2024).

56. Lampada, A. & Taylor, V. Notch signaling as a master regulator of adult neurogenesis. Front. Neurosci. 17, 1179011 (2023).

57. Killoy, K. M., Harlan, B. A., Pehar, M. & Vargas, M. R. FABP7 upregulation induces a neurotoxic phenotype in astrocytes. Glia 68, 2693–2704 (2020).

58. Harris, L. & Guillemot, F. HES1, two programs: promoting the quiescence and proliferation of adult neural stem cells. Genes Dev. 33, 479–481 (2019).

59. Wu, K. et al. The microglial innate immune receptors TREM-1 and TREM-2 in the anterior cingulate cortex (ACC) drive visceral hypersensitivity and depressive-like behaviors following DSS-induced colitis. Brain. Behav. Immun. 112, 96–117 (2023).

60. Tutukova, S., Tarabykin, V. & Hernandez-Miranda, L. R. The Role of Neurod Genes in Brain Development, Function, and Disease. Front. Mol. Neurosci. 14, 662774 (2021).

61. Miettinen, M., Clark, R. & Virtanen, I. Intermediate filament proteins in choroid plexus and ependyma and their tumors. Am. J. Pathol. 123, 231–240 (1986).

62. Harris, L., Genovesi, L. A., Gronostajski, R. M., Wainwright, B. J. & Piper, M. Nuclear factor one transcription factors: Divergent functions in developmental versus adult stem cell populations. Dev. Dyn. 244, 227–238 (2015).

63. McKinsey, G. L. et al. Radial glia integrin avb8 regulates cell autonomous microglial TGFβ1 signaling that is necessary for microglial identity. Nat. Commun. 16, 2840 (2025).

64. Cao, J. et al. The single-cell transcriptional landscape of mammalian organogenesis. Nature 566, 496– 502 (2019).

65. Kanton, S. et al. Organoid single-cell genomic atlas uncovers human-specific features of brain development. Nature 574, 418–422 (2019).

66. Chauvin, S. & Sobel, A. Neuronal stathmins: A family of phosphoproteins cooperating for neuronal development, plasticity and regeneration. Prog. Neurobiol. 126, 1–18 (2015).

67. Rocamonde, B., Herranz-Pérez, V., Garcia-Verdugo, J. M. & Huillard, E. ID4 Is Required for Normal Ependymal Cell Development. Front. Neurosci. 15, 668243 (2021).

68. Tzeng, S.-F. & De Vellis, J. Id1, Id2, and Id3 gene expression in neural cells during development. Glia 24, 372–381 (1998).

69. Hu, Z. et al. Involvement of LIN28A in Wnt-dependent regulation of hippocampal neurogenesis in the aging brain. Stem Cell Rep. 17, 1666–1682 (2022).

70. Zhang, J. et al. The m5C reader Ybx1 regulates embryonic cortical neurogenesis by promoting progenitor cell cycle progression. PLOS Biol. 23, e3003175 (2025).

71. Boucherie, C. et al. Auts2 enhances neurogenesis and promotes expansion of the cerebral cortex. J. Adv. Res. 72, 151–163 (2025).

72. Kantarci, H., Edlund, R. K., Groves, A. K. & Riley, B. B. Tfap2a Promotes Specification and Maturation of Neurons in the Inner Ear through Modulation of Bmp, Fgf and Notch Signaling. PLOS Genet. 11, e1005037 (2015).

73. Brainard, M. S. & Doupe, A. J. Translating Birdsong: Songbirds as a Model for Basic and Applied Medical Research. Annu. Rev. Neurosci. 36, 489–517 (2013).

74. Costa, F. V., et al. Current State of Modeling Human Psychiatric Disorders Using Zebrafish. Int. J. Mol. Sci. 24, 3187 (2023).

75. Sommer-Trembo, C. et al. The genetics of niche-specific behavioral tendencies in an adaptive radiation of cichlid fishes. Science 384, 470–475 (2024).

76. Major Depressive Disorder Working Group of the Psychiatric Genomics Consortium et al. Genetic identification of brain cell types underlying schizophrenia. Nat. Genet. 50, 825–833 (2018).

77. De Leeuw, C. A., Mooij, J. M., Heskes, T. & Posthuma, D. MAGMA: Generalized Gene-Set Analysis of GWAS Data. PLOS Comput. Biol. 11, e1004219 (2015).

78. Murphy, A. E., Schilder, B. M. & Skene, N. G. MungeSumstats: a Bioconductor package for the standardization and quality control of many GWAS summary statistics. Bioinformatics 37, 4593–4596 (2021).

79. Shafer, M. E. R., Sawh, A. N. & Schier, A. F. Gene family evolution underlies cell-type diversification in the hypothalamus of teleosts. *Nat*. Ecol. Evol. 6, 63–76 (2021).

80. Zhu, Y. et al. Whole-genome duplication shaped cell-type evolution in the vertebrate brain. Nature 656, 659–669 (2026).

81. Mah, J. L. & Dunn, C. W. Cell type evolution reconstruction across species through cell phylogenies of single-cell RNA sequencing data. *Nat*. Ecol. Evol. 8, 325–338 (2024).

82. Gerber, T. et al. Single-cell analysis uncovers convergence of cell identities during axolotl limb regeneration. Science 362, eaaq0681 (2018).

83. Cellular basis of accelerated whole-tooth regeneration. eLife 15, RP110584 (2026).

84. Bendesky, A. Behavioral evolution by diverging cell type composition. Curr. Opin. Genet. Dev. 95, 102397 (2025).

85. Niepoth, N. et al. Evolution of a novel adrenal cell type that promotes parental care. Nature 629, 1082– 1090 (2024).

86. Woych, J. et al. Cell-type profiling in salamanders identifies innovations in vertebrate forebrain evolution. Science 377, eabp9186 (2022).

87. Camacho, C. et al. BLAST+: architecture and applications. BMC Bioinformatics 10, 421 (2009).

88. Traag, V. A., Waltman, L. & Van Eck, N. J. From Louvain to Leiden: guaranteeing well-connected communities. Sci. Rep. 9, 5233 (2019).

89. Lamanna, F. Supplementary Data Files from: A lamprey neural cell type atlas illuminates the origins of the vertebrate brain. Zenodo 10.5281/ZENODO.5903844 (2023).

90. Schloissnig, S., et al. The giant axolotl genome uncovers the evolution, scaling, and transcriptional control of complex gene loci. Proc. Natl. Acad. Sci. 118, e2017176118 (2021).

91. Hao, Y. et al. Dictionary learning for integrative, multimodal and scalable single-cell analysis. Nat. Biotechnol. 42, 293–304 (2024).

92. Choudhary, S. & Satija, R. Comparison and evaluation of statistical error models for scRNA-seq. Genome Biol. 23, 27 (2022).

93. Korsunsky, I. et al. Fast, sensitive and accurate integration of single-cell data with Harmony. Nat. Methods 16, 1289–1296 (2019).

94. Chari, T. et al. Whole-animal multiplexed single-cell RNA-seq reveals transcriptional shifts across *Clytia* medusa cell types. Sci. Adv. 7, eabh1683 (2021).

95. Kaplan, H. S. et al. Sensory input, sex and function shape hypothalamic cell type development. Nature 647, 157–168 (2025).

96. Paradis, E. & Schliep, K. ape 5.0: an environment for modern phylogenetics and evolutionary analyses in R. Bioinformatics 35, 526–528 (2019).

97. Smith, M. R. Using Information Theory to Detect Rogue Taxa and Improve Consensus Trees. Syst. Biol. 71, 1088–1094 (2022).

98. Major Depressive Disorder Working Group of the Psychiatric Genomics Consortium et al. Genetic identification of brain cell types underlying schizophrenia. Nat. Genet. 50, 825–833 (2018).

99. De Leeuw, C. A., Mooij, J. M., Heskes, T. & Posthuma, D. MAGMA: Generalized Gene-Set Analysis of GWAS Data. PLOS Comput. Biol. 11, e1004219 (2015).

100. Karczewski, K. J. et al. Pan-UK Biobank genome-wide association analyses enhance discovery and resolution of ancestry-enriched effects. Nat. Genet. 57, 2408–2417 (2025).

101. Autism Spectrum Disorder Working Group of the Psychiatric Genomics Consortium et al. Identification of common genetic risk variants for autism spectrum disorder. Nat. Genet. 51, 431–444 (2019).

102. Howard, D. M. et al. Genome-wide meta-analysis of depression identifies 102 independent variants and highlights the importance of the prefrontal brain regions. Nat. Neurosci. 22, 343–352 (2019).

103. Sullivan, P. bip2021_noUKBB. 385000295 Bytes figshare 10.6084/M9.FIGSHARE.22564402.V1 (2023).

104. Demontis, D. et al. Genome-wide analyses of ADHD identify 27 risk loci, refine the genetic architecture and implicate several cognitive domains. Nat. Genet. 55, 198–208 (2023).

105. Ruderfer, D. M. et al. Genomic Dissection of Bipolar Disorder and Schizophrenia, Including 28 Subphenotypes. Cell 173, 1705–1715.e16 (2018).

106. Liu, Y. & Li, G. Empowering biologists to decode omics data: the Genekitr R package and web server. BMC Bioinformatics 24, 214 (2023).

