## Supplemental Data 1 for "Evolutionarily informed gene sets reveal conserved and lineage-modified transcriptional programs during vertebrate forebrain evolution"

**SUPPLEMENT**

**FIGURES**

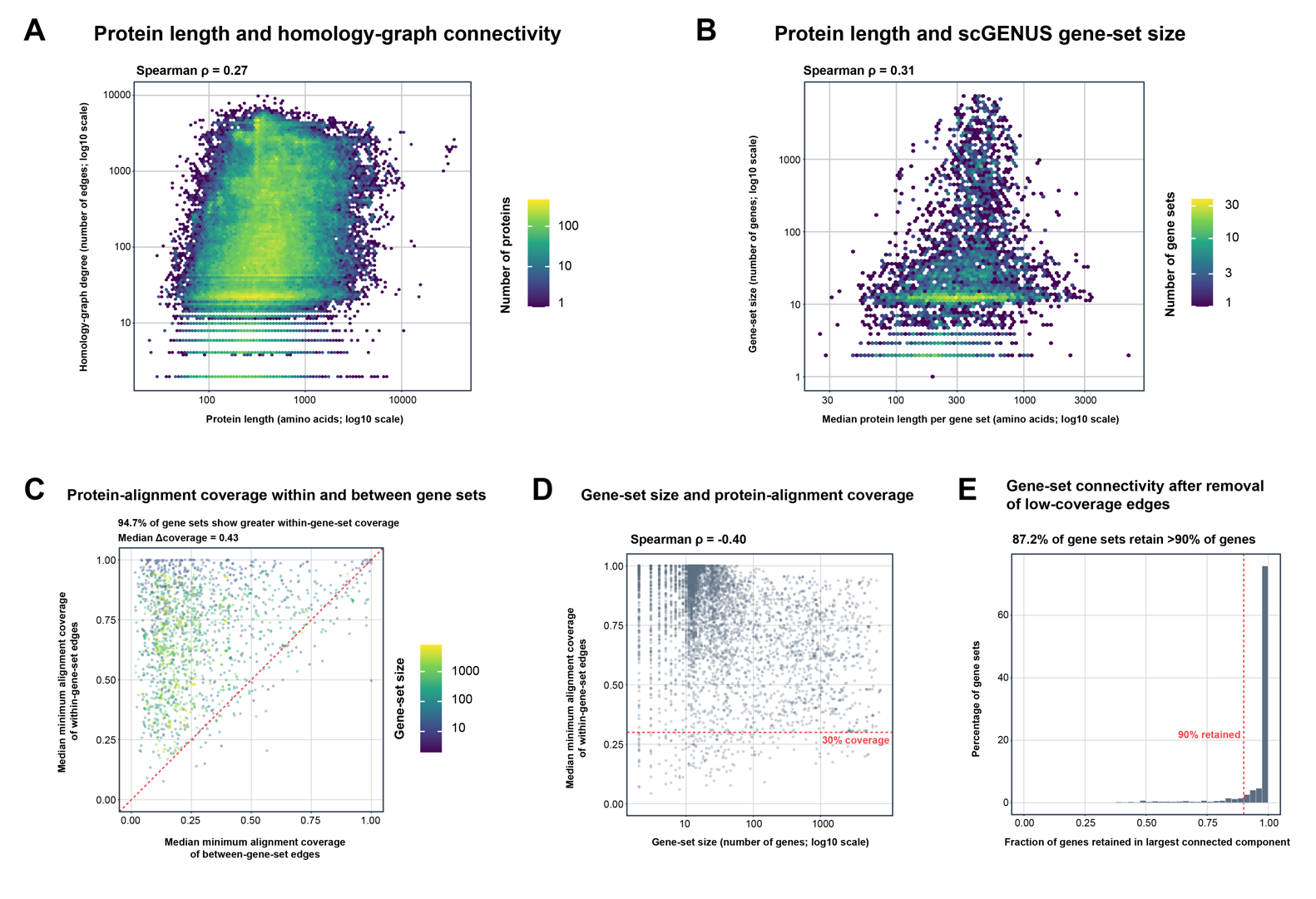

**Figure S1. Effects of protein sequence properties on scGENUS gene-set construction.** (A) Relationship between protein length and connectivity in the global protein-homology graph. Homology-graph degree represents the number of graph edges associated with each protein/gene, and both axes are displayed on logarithmic scales. Protein length showed a modest positive association with graph degree (Spearman $\rho=0.27$). Color indicates the local number of proteins. (B) Relationship between median protein length within each scGENUS gene set and gene-set size. Median protein length showed a modest positive association with the number of genes assigned to a gene set (Spearman $\rho=0.31$). Color indicates the local number of gene sets. (C) Comparison of protein-alignment coverage for edges connecting genes within the same scGENUS gene set versus edges connecting genes assigned to different gene sets. For each graph edge, overlapping BLASTp high-scoring-pair (HSP) intervals were merged separately for the two proteins, and alignment coverage was calculated as the fraction of each protein sequence covered by the merged HSPs. The smaller of the two coverage fractions was used as the edge-level minimum alignment coverage. For each gene set, median minimum coverage was calculated separately for within-gene-set and between-gene-set edges. A total of 94.7% of gene sets exhibited greater median alignment coverage for within-gene-set edges than for between-gene-set edges, with a median difference of 0.43. The dashed diagonal indicates equal within- and between-gene-set coverage. (D) Relationship between gene-set size and median minimum protein-alignment coverage among within-gene-set edges. Larger gene sets showed moderately lower alignment coverage (Spearman $\rho=-0.40$). The dashed horizontal line indicates the 30% minimum protein-coverage threshold used for the edge-filtering analysis in panel E. (E) Robustness of gene-set connectivity after removal of graph edges with minimum protein-alignment coverage below 30%. For each gene set, connectivity was quantified as the fraction of its genes remaining in the largest connected component after edge filtering relative to the number of genes in the original gene set. Overall, 87.2% of gene sets retained more than 90% of their genes within the largest connected component after removal of low-coverage edges. The dashed vertical line denotes the 90% retention threshold.

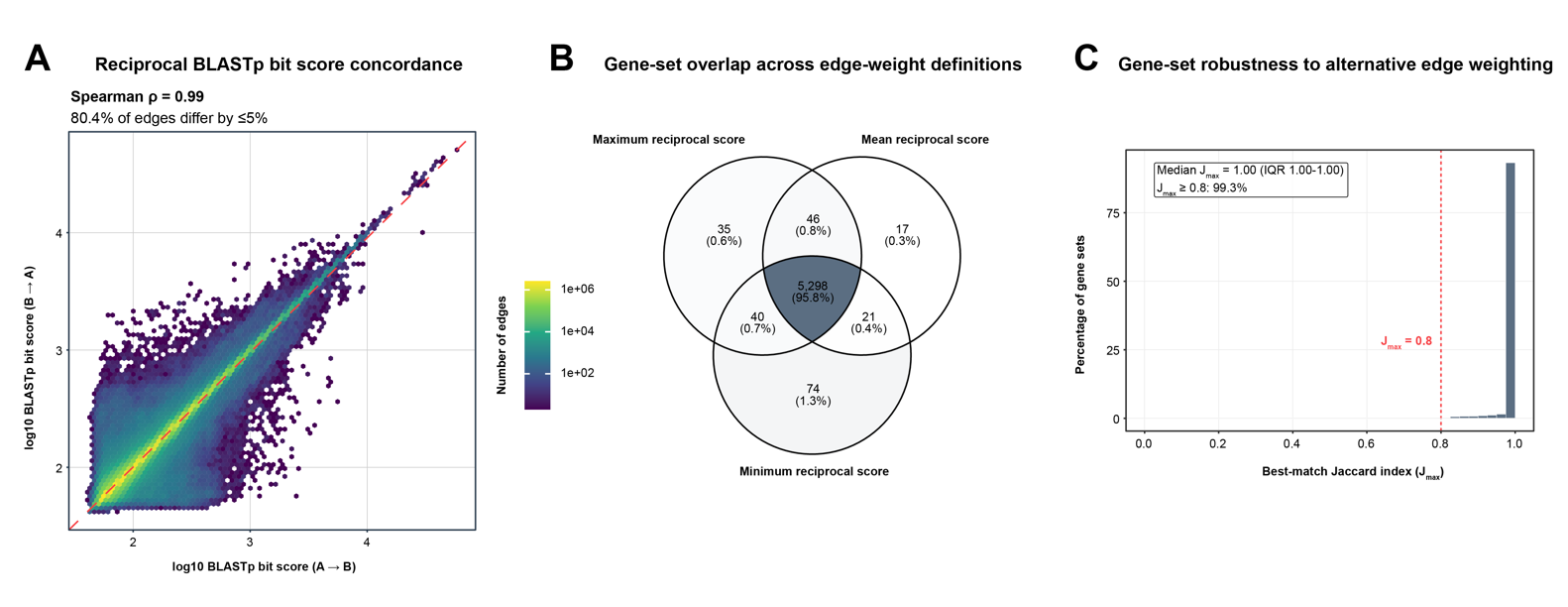

**Figure S2. Robustness of scGENUS gene sets to reciprocal BLASTp edge-weight definition.** (A) Concordance of reciprocal BLASTp bit scores for edges in the gene-homology graph. Each axis represents the log10-transformed BLASTp bit score obtained for the same gene pair in opposite search directions (A $\to$ B and B $\to$ A). Reciprocal scores were highly concordant (Spearman $\rho=0.99$), and 80.4% of graph edges differed by no more than 5% between the two directions. Color indicates the local number of graph edges, and the dashed diagonal denotes equal reciprocal bit scores. (B) Overlap among scGENUS gene-set collections reconstructed using three alternative symmetric edge-weight definitions: the maximum, mean, or minimum of the reciprocal BLASTp bit scores. Gene sets across reconstructions were considered matched when their Jaccard overlap was at least 0.8. A total of 5,298 gene sets (95.8%) were represented across all three edge-weight definitions, indicating extensive agreement among the resulting gene-set collections. Numbers and percentages indicate gene sets assigned to each region of the overlap diagram. (C) Gene-set similarity under alternative reciprocal-score weighting. For each gene set constructed using the primary maximum-reciprocal-score definition, the best-match Jaccard index ($J_{max}$) was defined as the highest Jaccard overlap with any gene set reconstructed using either the mean- or minimum-reciprocal-score weighting. The median $J_{max}$ was 1.00 (IQR, 1.00-1.00), and 99.3% of primary gene sets had a best-match Jaccard index of at least 0.8 (dashed vertical line).

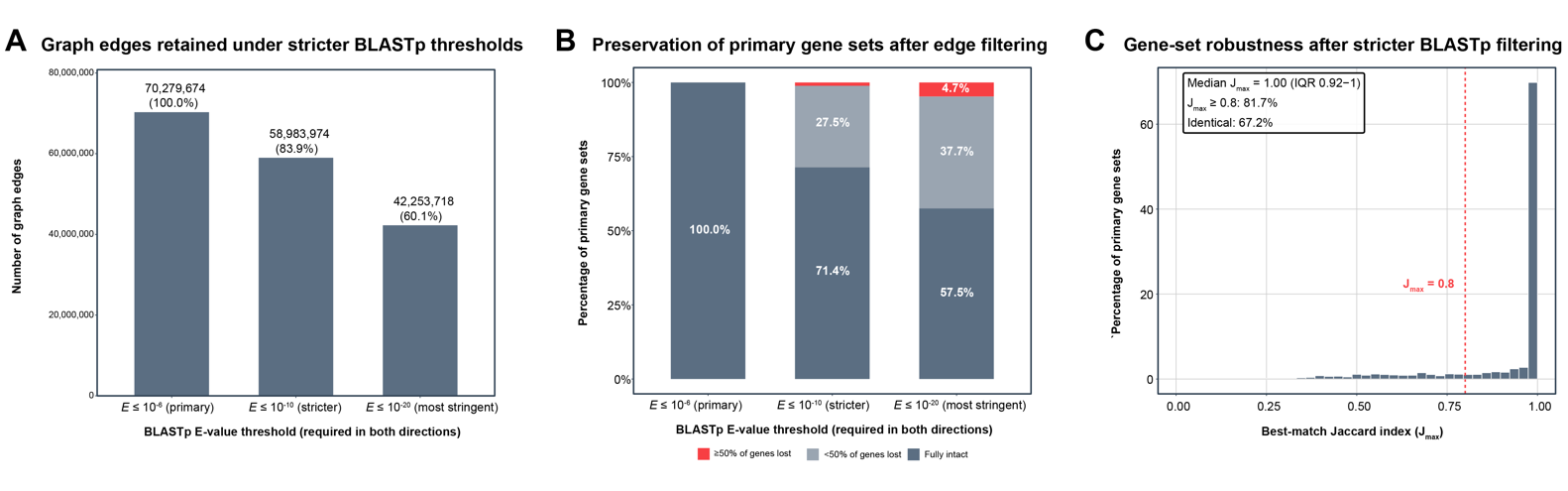

**Figure S3. Sensitivity of scGENUS gene-set construction to BLASTp E-value thresholds.** (A) Number of edges retained in the global gene-homology graph after progressively tightening the reciprocal BLASTp E-value threshold from the primary threshold of $E \leq{10}^{-6}$ to $E \leq{10}^{-10}$ and $E \leq{10}^{-20}$, with the threshold required in both search directions. Relative to the primary graph containing 70,279,674 edges, 83.9% and 60.1% of edges were retained at $E \leq{10}^{-10}$ and $E \leq{10}^{-20}$, respectively. (B) Preservation of primary scGENUS gene sets after applying the stricter E-value filters to the homology graph. For each primary gene set, we determined the proportion of its original member genes retained after applying the stricter $E \leq{10}^{-10}$ or $E \leq{10}^{-20}$ threshold. Gene sets were classified as fully intact, having lost <50% of their genes, or having lost ≥50% of their genes. At $E \leq{10}^{-10}$, 71.4% of primary gene sets remained fully intact and 27.5% lost fewer than 50% of their genes; at $E \leq{10}^{-20}$, 57.5% remained fully intact, 37.7% lost fewer than 50% of their genes, and 4.7% lost at least 50% of their genes. (C) Similarity between the primary gene sets and gene sets reconstructed de novo using the stricter $E \leq{10}^{-10}$ BLASTp threshold. For each primary gene set, the best-match Jaccard index ($J_{max}$) was defined as the maximum Jaccard overlap with any gene set reconstructed under the stricter threshold. The median $J_{max}$ was 1.00 (IQR, 0.92–1.00), 81.7% of primary gene sets had a best-match Jaccard index of at least 0.8, and 67.2% were identical to a reconstructed gene set. The dashed vertical line denotes $J_{max}=0.8$.

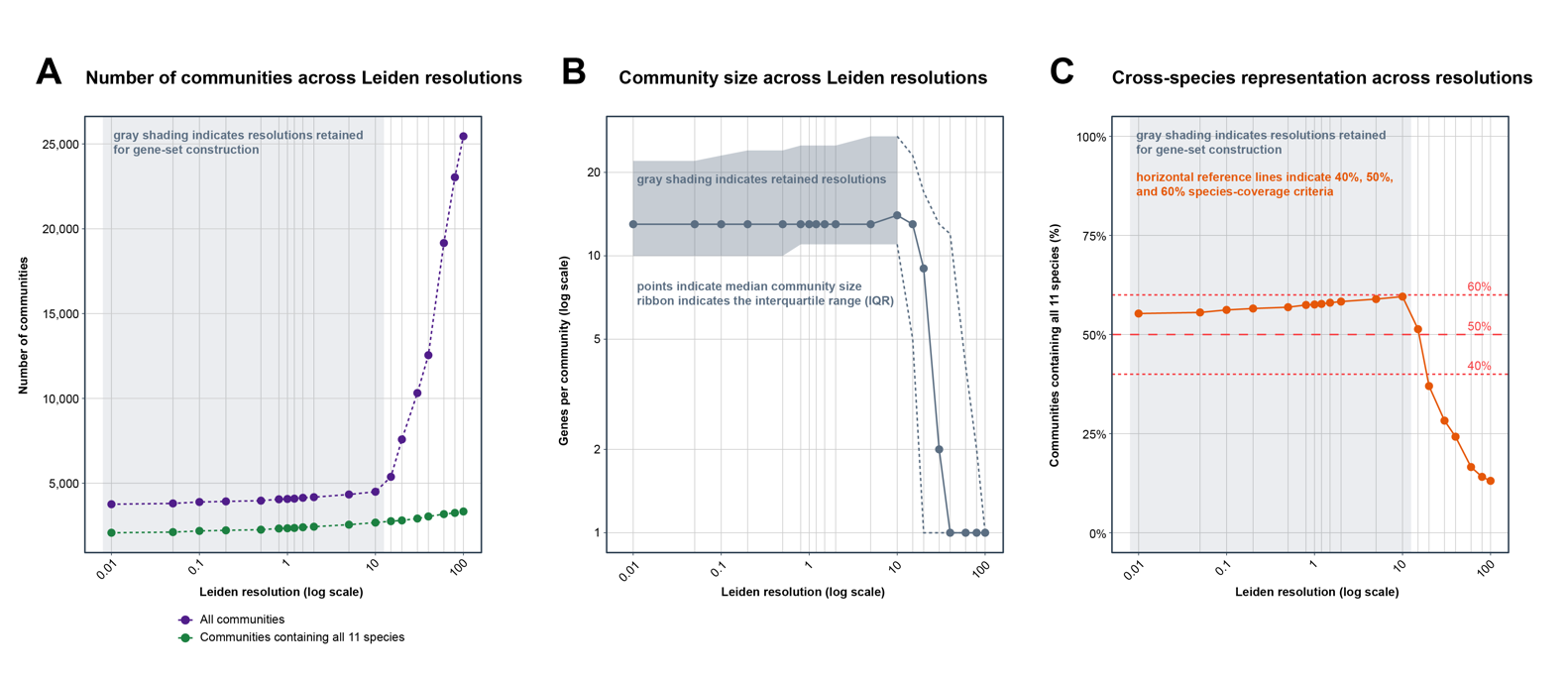

**Figure S4. Stability of Leiden partitioning across resolutions.** (A) Number of communities identified across the tested Leiden resolution parameters. Purple points indicate the total number of communities, and green points indicate the number of communities containing genes from all 11 species. The vertical gray background indicates the range of Leiden resolutions retained for scGENUS gene-set construction. (B) Community-size distributions across Leiden resolutions. Points indicate the median number of genes per community, and the gray ribbon indicates the interquartile range (IQR). Median community size remained relatively stable across the retained resolution range but decreased sharply at higher resolutions as the homology graph was partitioned into progressively smaller communities. The vertical gray background indicates the resolutions retained for gene-set construction. (C) Cross-species representation across Leiden resolutions, quantified as the percentage of communities containing genes from all 11 species. Horizontal reference lines indicate 40%, 50%, and 60% species-coverage criteria, with 50% representing the primary criterion used during gene-set construction. The vertical gray background indicates the final retained resolution range. Together, these analyses show a broad range of resolutions with relatively stable community number, size, and cross-species representation, supporting the resolution range used to construct the final scGENUS gene-set collection.

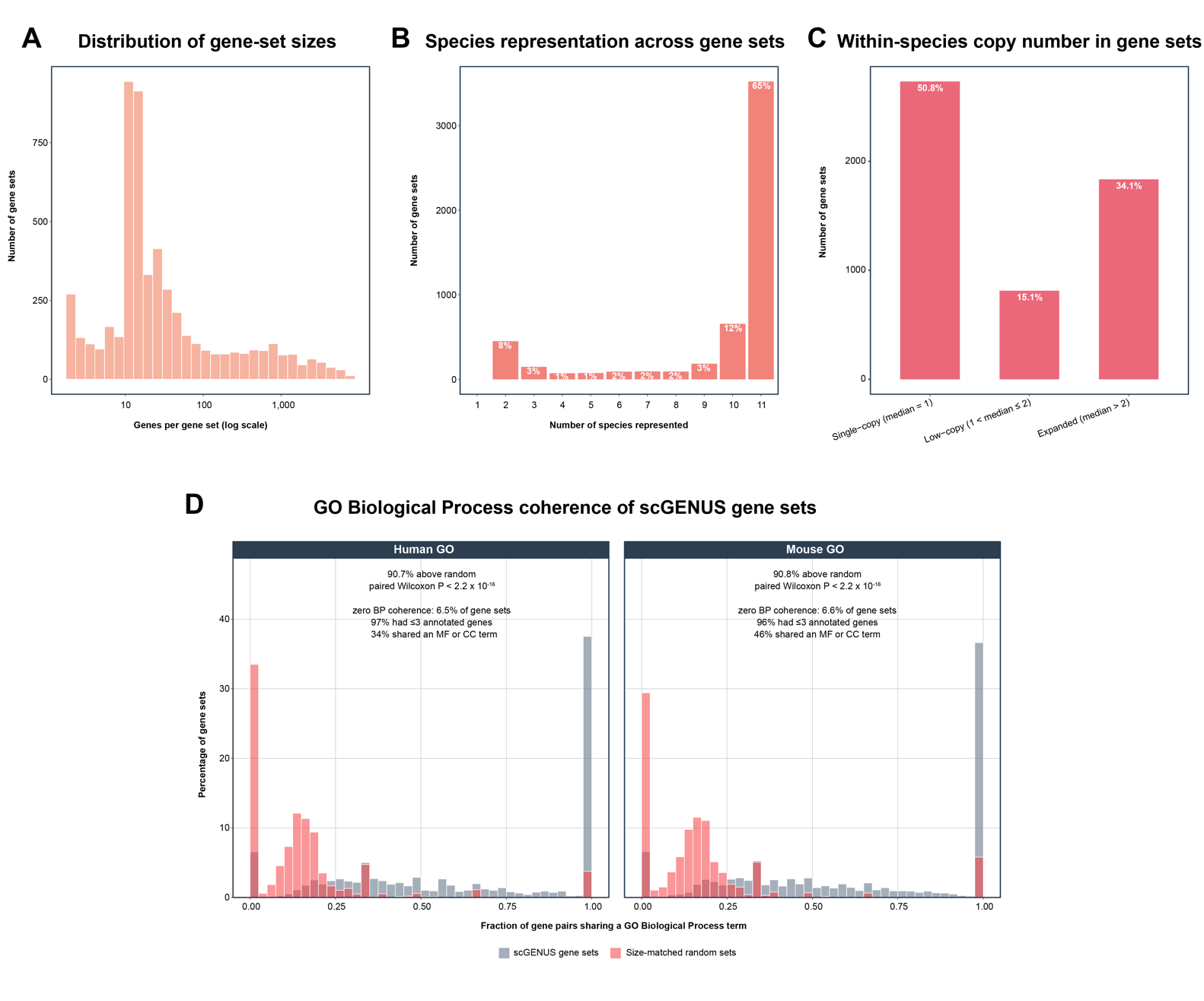

**Figure S5. Structural and functional characteristics of scGENUS gene sets.** (A) Distribution of gene-set sizes across the final 5,419 scGENUS gene sets. The x-axis indicates the number of genes assigned to each gene set on a logarithmic scale. (B) Species representation across scGENUS gene sets, quantified as the number of the 11 vertebrate species represented within each gene set. Percentages above bars indicate the proportion of gene sets in each category. Overall, 65% of gene sets contained genes from all 11 species, and an additional 12% contained genes from 10 species. (C) Within-species copy-number composition of scGENUS gene sets. Gene sets were classified according to the median number of member genes per represented species as single-copy (median = 1), low-copy (1 < median $\leq2$), or expanded (median > 2), comprising 50.8%, 15.1%, and 34.1% of gene sets, respectively. (D) Functional coherence of scGENUS gene sets assessed independently using human and mouse Gene Ontology (GO) Biological Process annotations. For each gene set, functional coherence was quantified as the fraction of gene pairs sharing at least one GO Biological Process term and compared with size-matched random gene sets from the corresponding species. scGENUS gene sets showed greater functional coherence than their size-matched random counterparts for 90.7% of human gene sets and 90.8% of mouse gene sets (paired Wilcoxon signed-rank test, P < 2.2 $\times$ 10^-16^ for both species). Zero Biological Process coherence was observed for 6.5% of human and 6.6% of mouse gene sets; among these, 97% and 96%, respectively, contained three or fewer GO-annotated genes, while 34% of human and 46% of mouse zero-coherence gene sets shared at least one GO Molecular Function or Cellular Component term.

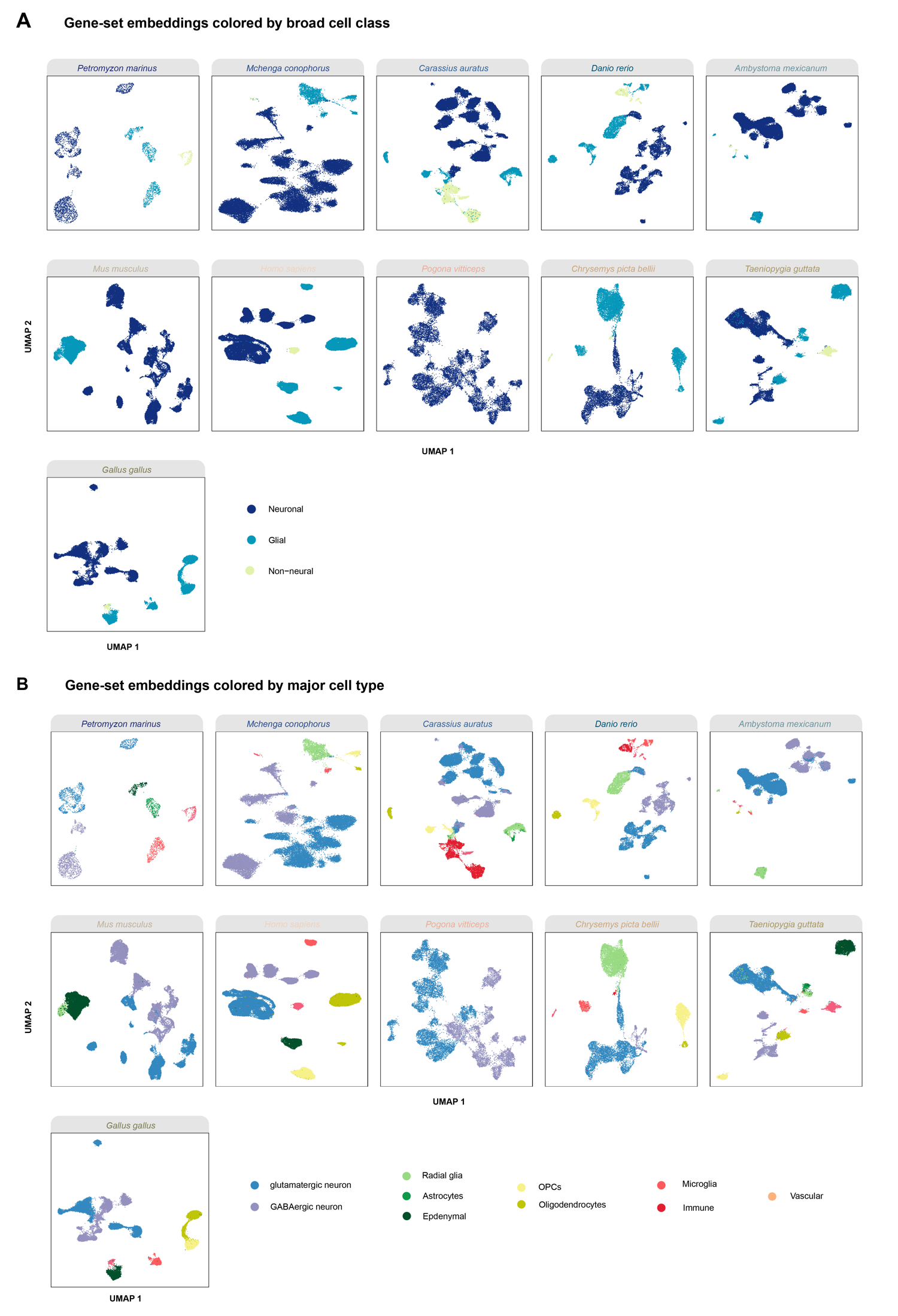

**Figure S6. Gene-set-based single-cell embeddings preserve cell-type structure within individual vertebrate datasets.** (A) Species-specific UMAP embeddings generated from scGENUS gene-set expression profiles, colored by broad cell class (neuronal, glial, and non-neural). Broad cellular classes occupy distinct regions of the gene-set-based embeddings across the individual vertebrate datasets. (B) The same species-specific UMAP embeddings colored by major cell type, including glutamatergic and GABAergic neurons, radial glia, astrocytes, ependymal cells, oligodendrocyte precursor cells (OPCs), oligodendrocytes, microglia, immune cells, and vascular cells, where represented in the source dataset. The separation of these annotated populations demonstrates that transformation from gene-level to scGENUS gene-set-level expression preserves biologically meaningful cellular heterogeneity within individual species prior to cross-species integration.

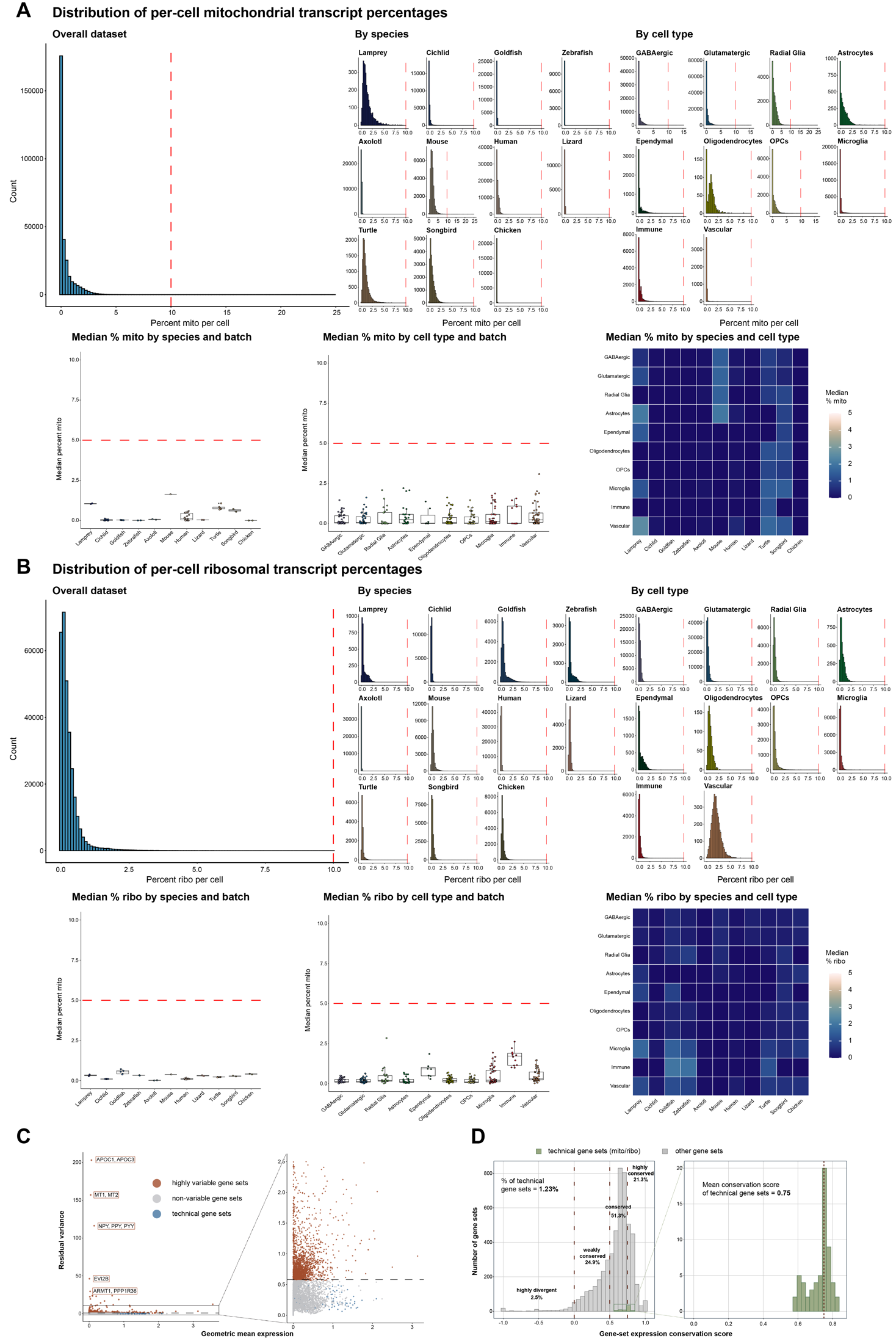

**Figure S7. Mitochondrial and ribosomal features have limited influence on scGENUS integration and conservation analyses.** (A) Distribution of mitochondrial transcript percentages calculated directly from the original gene-level expression matrices. Per-cell mitochondrial transcript percentages are shown for the complete dataset and separately by species and major cell type. The vertical dashed lines indicate 10% mitochondrial transcripts fraction. Lower panels summarize median mitochondrial transcript percentages across species and batches, across cell types and batches, and across species–cell-type combinations. Horizontal dashed lines indicate a 5% median mitochondrial transcript fraction. Mitochondrial transcript fractions were generally low across species, cell types, and batches. (B) Corresponding analysis of ribosomal transcript percentages calculated from the original gene-level expression matrices. Per-cell distributions are shown overall and stratified by species and cell type, with lower panels summarizing median ribosomal transcript percentages across species, batches, cell types, and species–cell-type combinations. Vertical dashed lines in the per-cell distributions indicate 10% ribosomal transcript fraction, whereas horizontal dashed lines in the median summaries indicate a 5% median ribosomal transcript fraction. Ribosomal transcript fractions were likewise generally low across datasets and cellular populations. (C) Assessment of mitochondrial and ribosomal scGENUS gene sets during variable feature selection. Residual variance is plotted against geometric mean expression following SCTransform-based feature selection; mitochondrial and ribosomal gene sets are highlighted relative to highly variable and non-variable gene sets. The expanded view highlights the region containing the technical gene sets. Of the 67 technical gene sets identified in the final scGENUS collection (15 mitochondrial and 52 ribosomal), only one was selected among the 4,000 variable gene sets used for PCA and subsequent Harmony integration. (D) Distribution of gene-set expression conservation scores for mitochondrial and ribosomal gene sets relative to all other scGENUS gene sets. Technical gene sets comprise 1.23% of the collection of 5,419 gene sets. Dashed vertical lines indicate the conservation score thresholds used to classify highly divergent, weakly conserved, conserved, and highly conserved gene sets. The right panel shows the conservation score distribution of mitochondrial and ribosomal gene sets alone, which exhibited a mean conservation score of approximately 0.75.

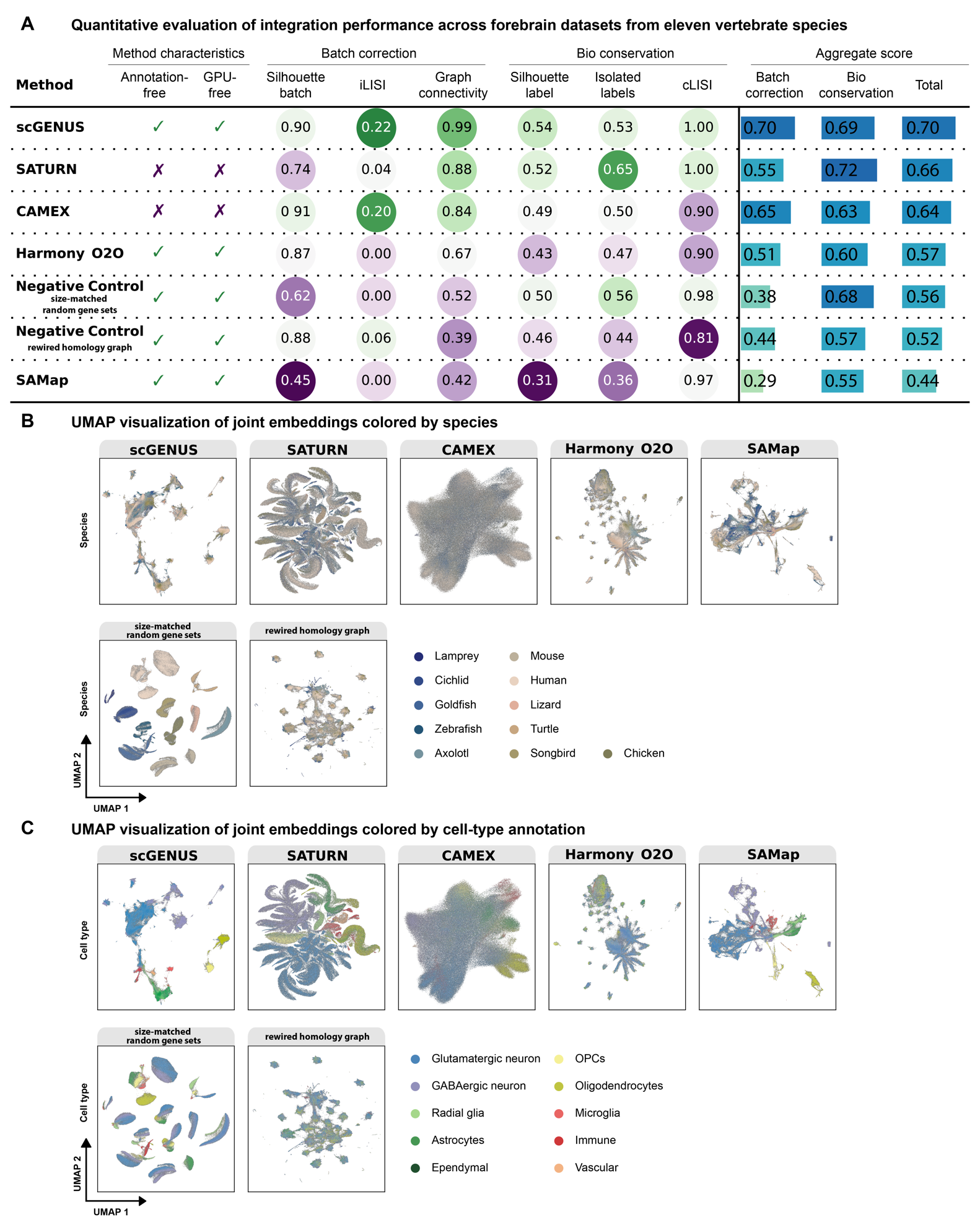

**Figure S8. Benchmarking scGENUS against cross-species integration methods across forebrain datasets from eleven vertebrate species.** (A) Quantitative comparison of scGENUS with alternative cross-species single-cell integration methods, including SATURN, CAMEX, Harmony using one-to-one ortholog features (Harmony O2O), and SAMap, together with two negative controls based on size-matched random gene sets and gene sets derived from a rewired homology graph. Method characteristics indicate whether integration can be performed without cell-type annotations and without GPU computation. Integration performance was evaluated using complementary metrics of batch correction and biological conservation. Batch-correction performance was assessed using the batch silhouette score, which measures mixing with respect to batch labels; integration local inverse Simpson’s index (iLISI), which quantifies local diversity of batch labels; and graph connectivity, which measures connectivity of corresponding biological populations across batches. Biological conservation was assessed using the cell-type-label silhouette score, which measures separation of annotated cell types; isolated-label preservation, which evaluates retention of cell populations represented in only a limited number of datasets; and cell-type local inverse Simpson’s index (cLISI), which quantifies local preservation of cell-type identity. All metrics were scaled such that higher values indicate better performance. Aggregate scores summarize batch correction, biological conservation, and overall integration performance. (B) UMAP visualizations of the joint cross-species embeddings generated by each integration method and negative control, with cells colored by species of origin. Effective integration is reflected by mixing of cells from different species rather than separation primarily by species identity. (C) The same joint embeddings colored by the original cell-type annotations, including glutamatergic and GABAergic neurons, radial glia, astrocytes, ependymal cells, oligodendrocyte precursor cells (OPCs), oligodendrocytes, microglia, immune cells, and vascular cells. Preservation of distinct cell-type structure while enabling cross-species mixing provides a complementary visual assessment of biological conservation and batch correction across integration approaches.

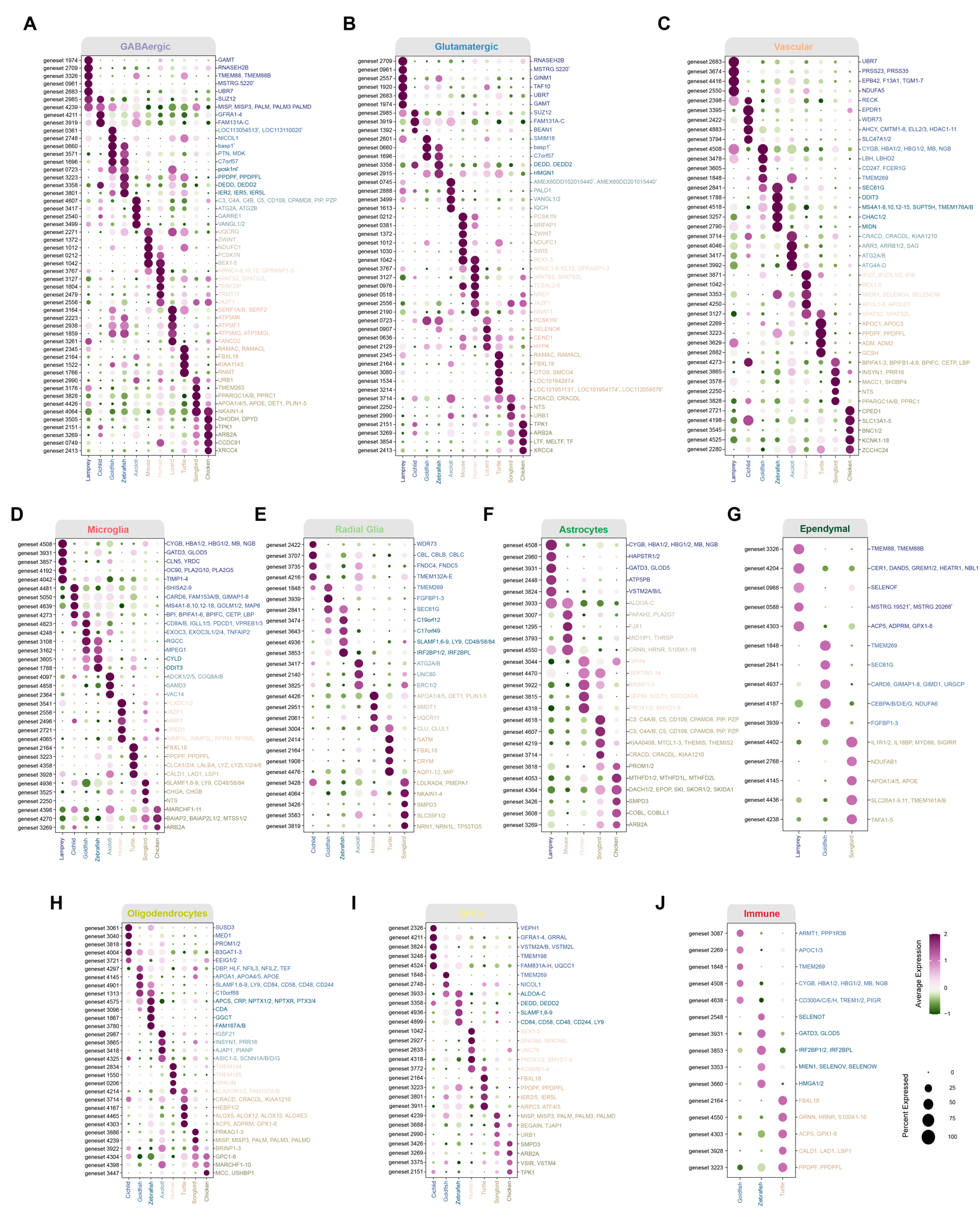

**Figure S9. Gene-set-level differential expression identifies species-enriched expression programs within vertebrate forebrain cell types.** Dot plots show scGENUS gene sets exhibiting species-enriched expression within (A) GABAergic neurons; (B) glutamatergic neurons; (C) vascular cells; (D) microglia; (E) radial glia; (F) astrocytes; (G) ependymal cells; (H) oligodendrocytes; (I) oligodendrocyte precursor cells (OPCs); (J) immune cells. Differential expression was evaluated separately within each cell type by comparing each species with the remaining species. Columns represent species, and rows represent differentially expressed scGENUS gene sets. Gene-set identifiers are listed on the left; representative human gene symbols within each gene set are shown on the right to facilitate annotation. Asterisks (*) denote gene sets lacking a human ortholog, in which case representative locus identifiers or species-specific gene names are displayed. Dot size indicates the fraction of cells expressing each gene set, and color represents scaled mean expression.

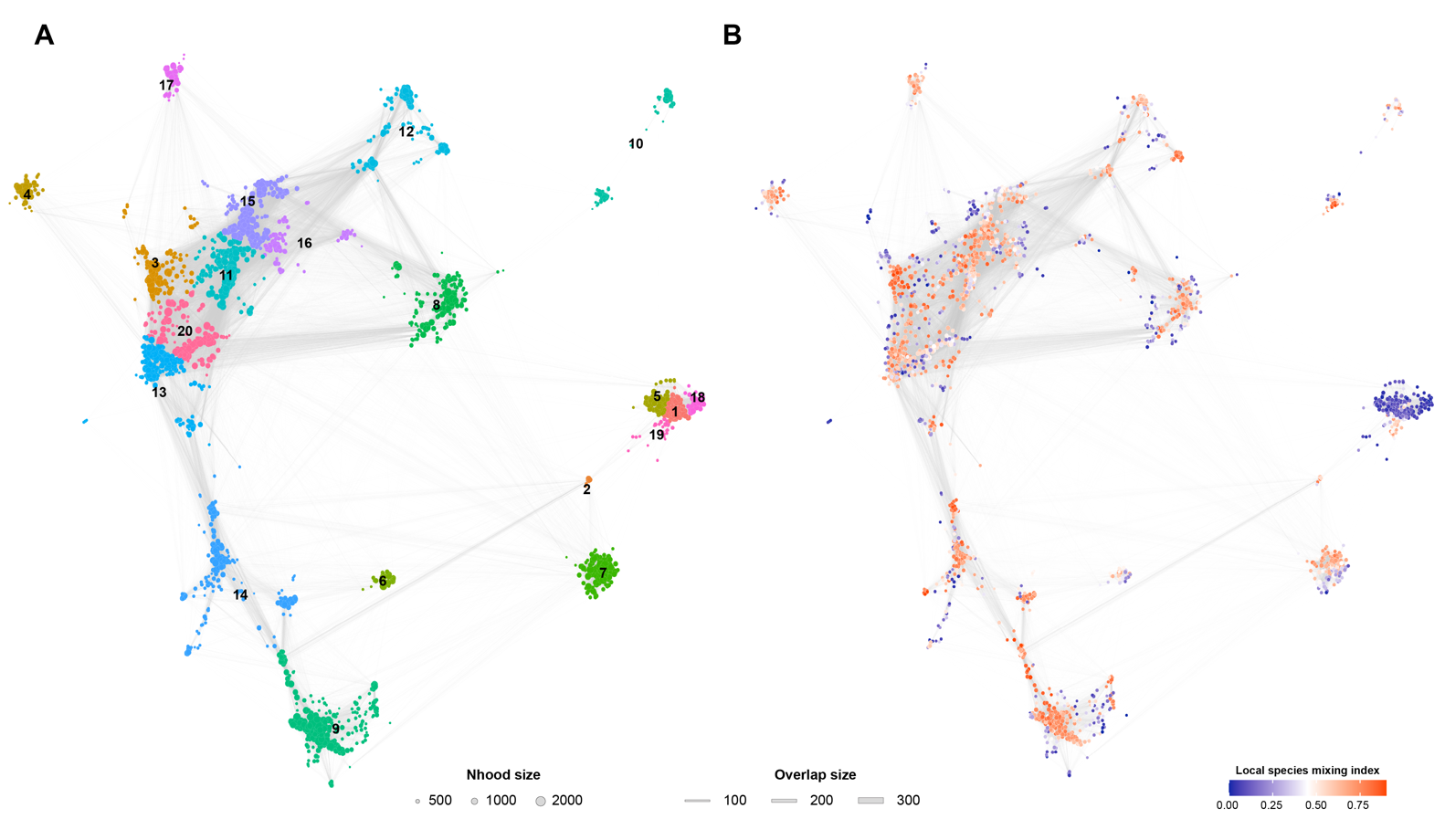

**Figure S10. Organization and local species mixing of Milo-defined transcriptional neighborhoods.** (A) Graph representation of transcriptional neighborhoods defined using Milo in the integrated scGENUS gene-set space. Each node represents a local neighborhood, node size indicates the number of cells in the neighborhood, and edge width reflects the number of cells shared between adjacent neighborhoods. Neighborhoods were grouped by k-means clustering (k = 20) of the neighborhood graph representation, with colors and numbers indicating the resulting neighborhood clusters. (B) The same neighborhood graph colored by the local species mixing index, calculated from species composition after normalization to global species frequencies. Lower values indicate greater dominance by a single species (0 indicating single-species dominance), whereas higher values indicate greater local mixing among species (1 indicating maximal mixing). Neighborhood clusters with reduced species mixing were used to identify the species-biased regions highlighted in Fig. 2E.

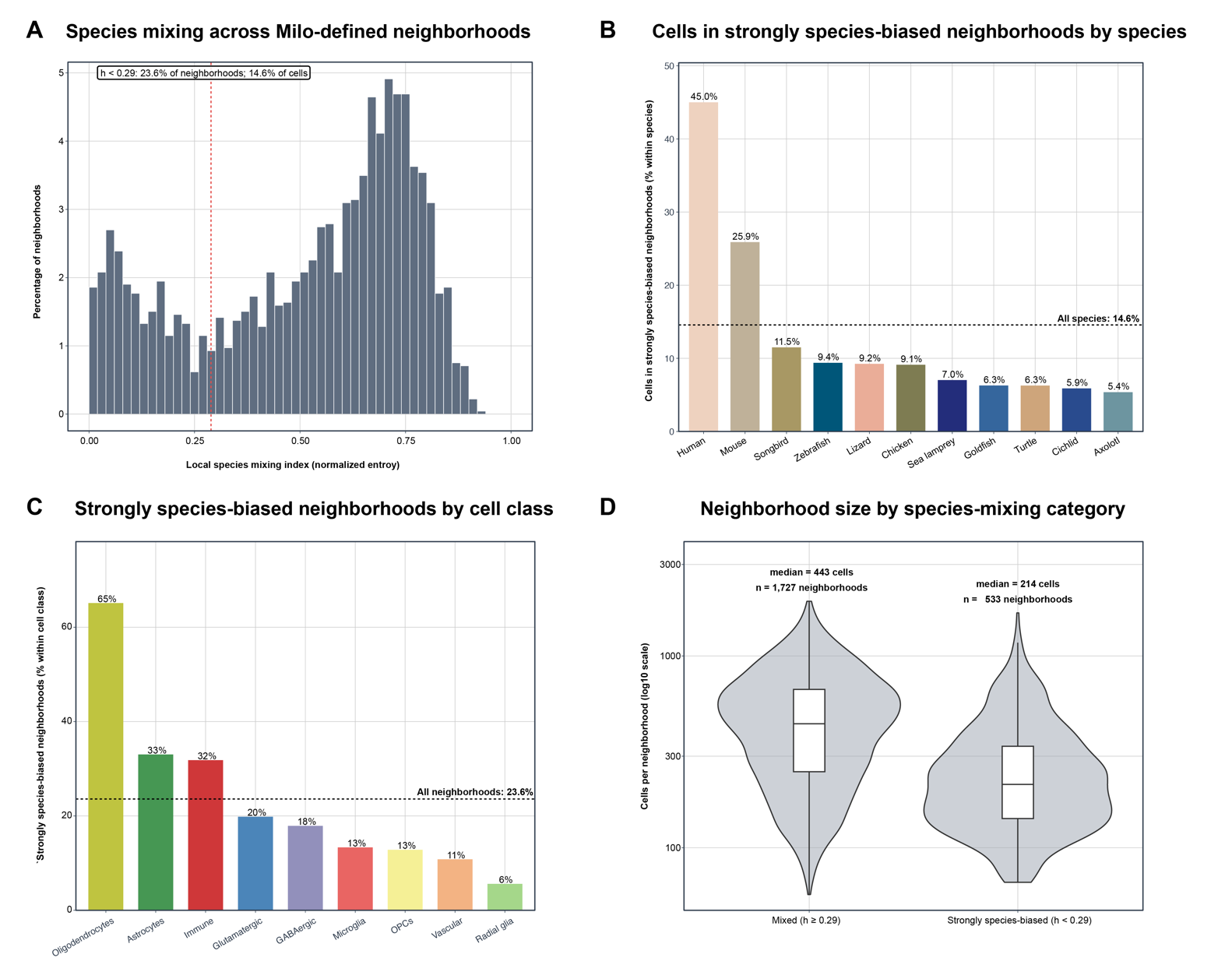

**Figure S11. Systematic characterization of strongly species-biased transcriptional neighborhoods in the integrated vertebrate forebrain atlas.** (A) Distribution of local species mixing across Milo-defined transcriptional neighborhoods in the integrated scGENUS gene-set space. Species mixing was quantified using normalized species entropy ($h$) calculated from species composition after normalization to global species frequencies, with values ranging from 0 (single-species dominance) to 1 (maximal cross-species mixing). The $h=0.29$ cutoff corresponds to the local minimum separating the low-entropy neighborhood mode from the broader distribution of mixed neighborhoods and was used to define strongly species-biased neighborhoods. Using this criterion, 23.6% of neighborhoods were strongly species-biased and collectively contained 14.6% of cells. The dashed vertical line indicates $h=0.29$. (B) Fraction of cells from each species located within strongly species-biased neighborhoods. For each species, bars indicate the percentage of all cells from that species assigned to neighborhoods with $h<0.29$. The horizontal dashed line indicates the corresponding fraction across all species combined (14.6%). (C) Frequency of strongly species-biased neighborhoods across major cell classes. Each neighborhood was assigned to its dominant cell class based on the original cell-type annotations, and bars indicate the percentage of neighborhoods within each cell class meeting the $h<0.29$ criterion. The horizontal dashed line indicates the overall proportion of strongly species-biased neighborhoods across all cell classes (23.6%). (D) Neighborhood-size distributions for mixed ($h\geq0.29$) and strongly species-biased ($h<0.29$) neighborhoods. Mixed neighborhoods contained a median of 443 cells (n = 1,727 neighborhoods), whereas strongly species-biased neighborhoods contained a median of 214 cells (n = 533 neighborhoods). Neighborhood size is displayed on a logarithmic scale.

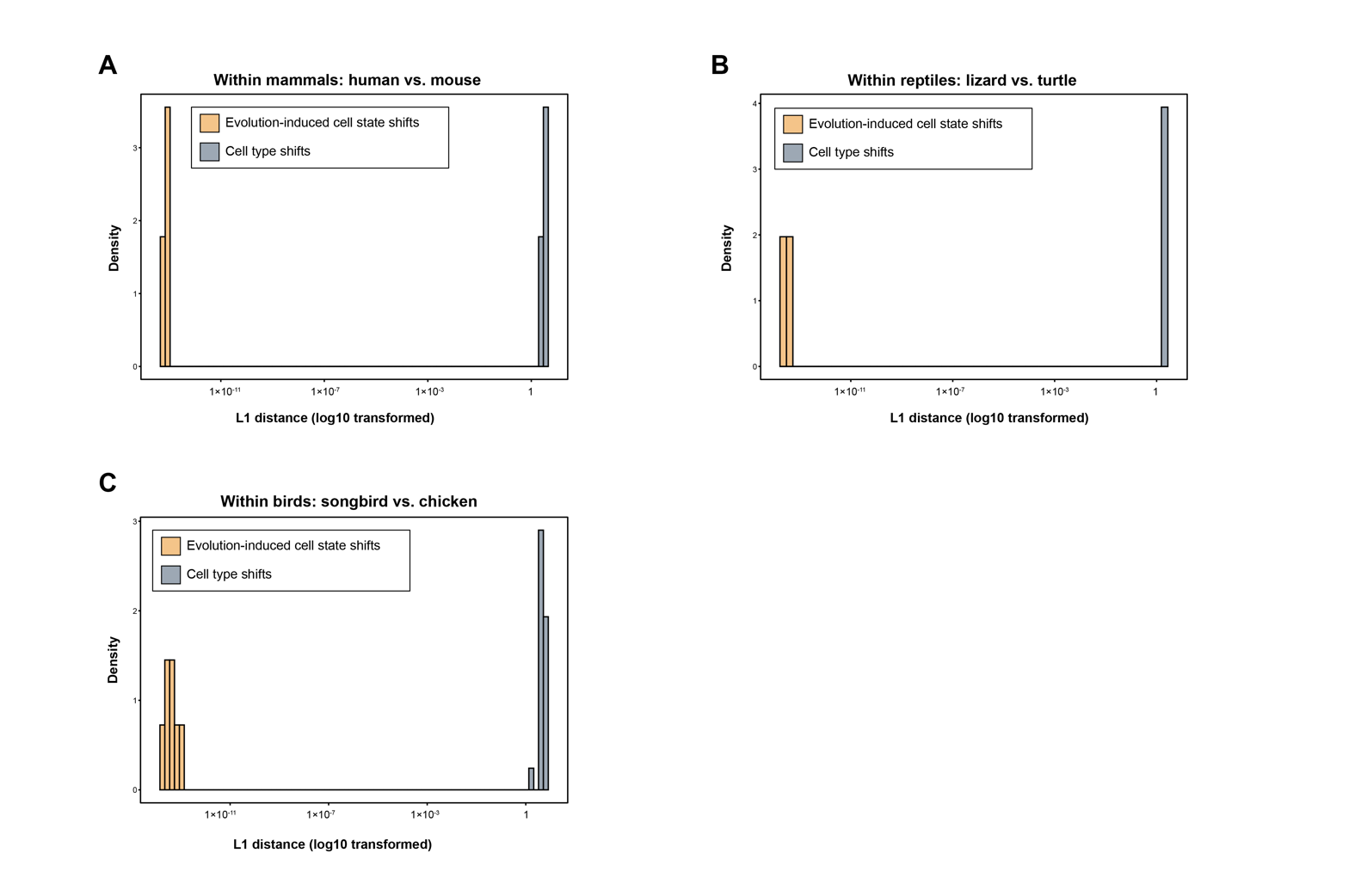

**Figure S12. Sublineage-restricted comparison of within- and between-cell-type transcriptional distances.** (A) Distribution of L1 distances between human and mouse species-specific centroids for the same cell type (within-cell-type interspecies distances) compared with L1 distances between centroids of distinct cell types (between-cell-type distances) within the mammalian comparison. (B) Corresponding comparison for reptiles using lizard and turtle. (C) Corresponding comparison for birds using songbird and chicken. Across each vertebrate lineage, within-cell-type interspecies distances were substantially smaller than distances between distinct cell types, supporting preservation of broad cell-type identity despite lineage-specific transcriptional divergence. Distances are displayed on a logarithmic scale.

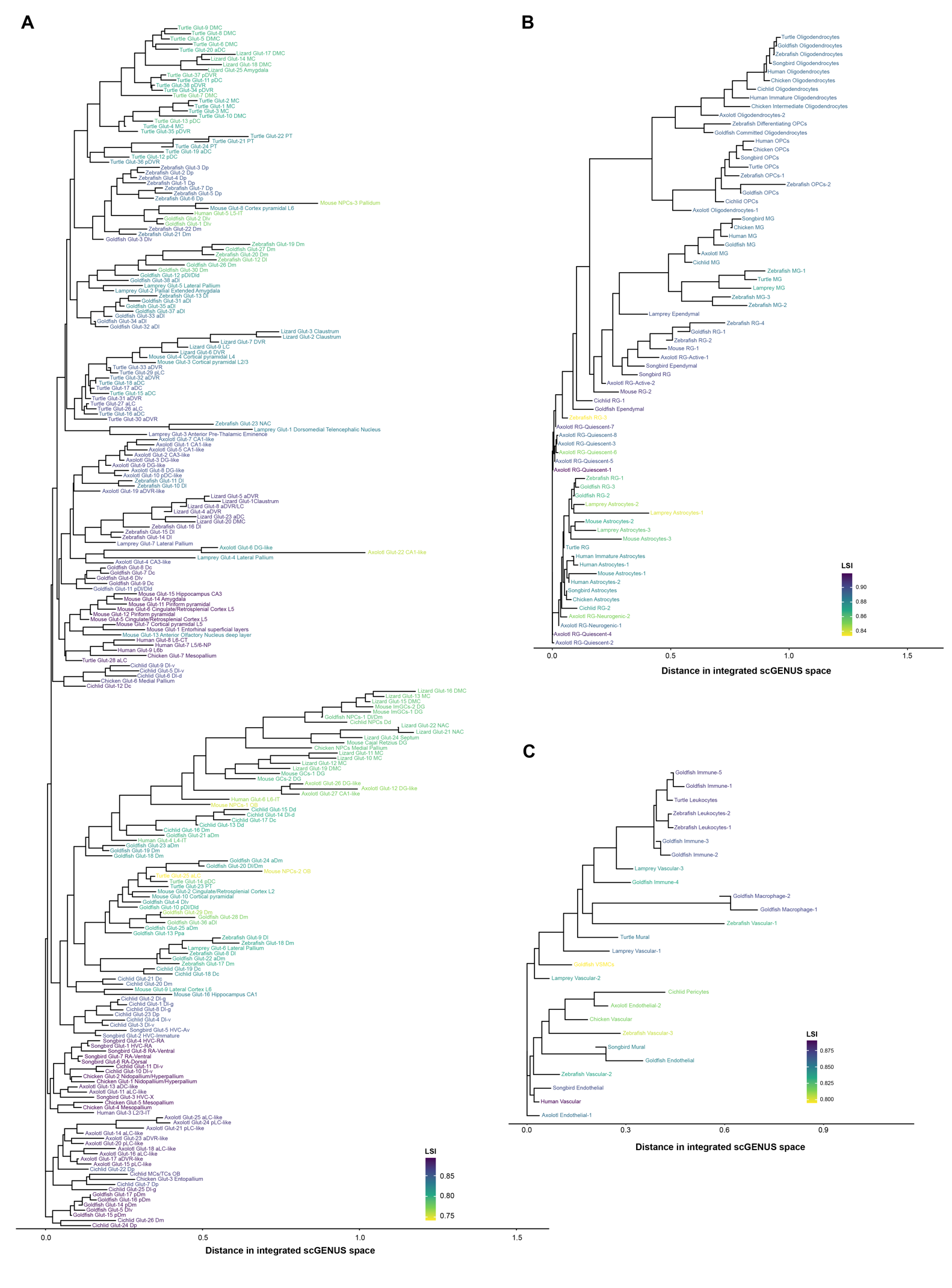

**Figure S13. Distance-based organization of glutamatergic, glial, and non-neural forebrain cell types in the integrated scGENUS space.** (A) Unrooted distance-based tree of glutamatergic neuronal subtypes constructed from subtype centroids in the first 50 Harmony-corrected principal components of the scGENUS gene-set representation. Subtypes from the eleven vertebrate species are shown at the tips. Branch lengths represent distances among subtype centroids in the integrated scGENUS space. Anatomical subdivisions are indicated in the subtype labels. Abbreviations: D, area dorsalis; V, area ventralis; Dc, large-celled subdivision of Dm; Vsst, ventral Sst; Ppa, nucleus preopticus parvocellularis anterioris; a, anterior; p, posterior; d, dorsal; v, ventral; m, medial; l, lateral; MC, medial cortex; DMC, dorsomedial cortex; pDC and aDC, posterior and anterior dorsal cortex; PT, pallial thickening; aLC and pLC, anterior and posterior lateral cortex; aDVR and pDVR, anterior and posterior dorsal ventricular ridge; DG, dentate gyrus; CA, cornu ammonis; OB, olfactory bulb; NAC, nucleus accumbens neurons. Tip color represents the leaf stability index (LSI) calculated from 1,000 bootstrap resampling iterations, with higher values indicating more stable tip placement across bootstrap trees. (B) Corresponding distance-based tree for glial populations, including radial glia, astrocytes, ependymal cells, oligodendrocyte precursor cells (OPCs), oligodendrocytes, and microglia. (C) Corresponding distance-based tree for non-neural populations, including vascular and peripheral immune cell types.

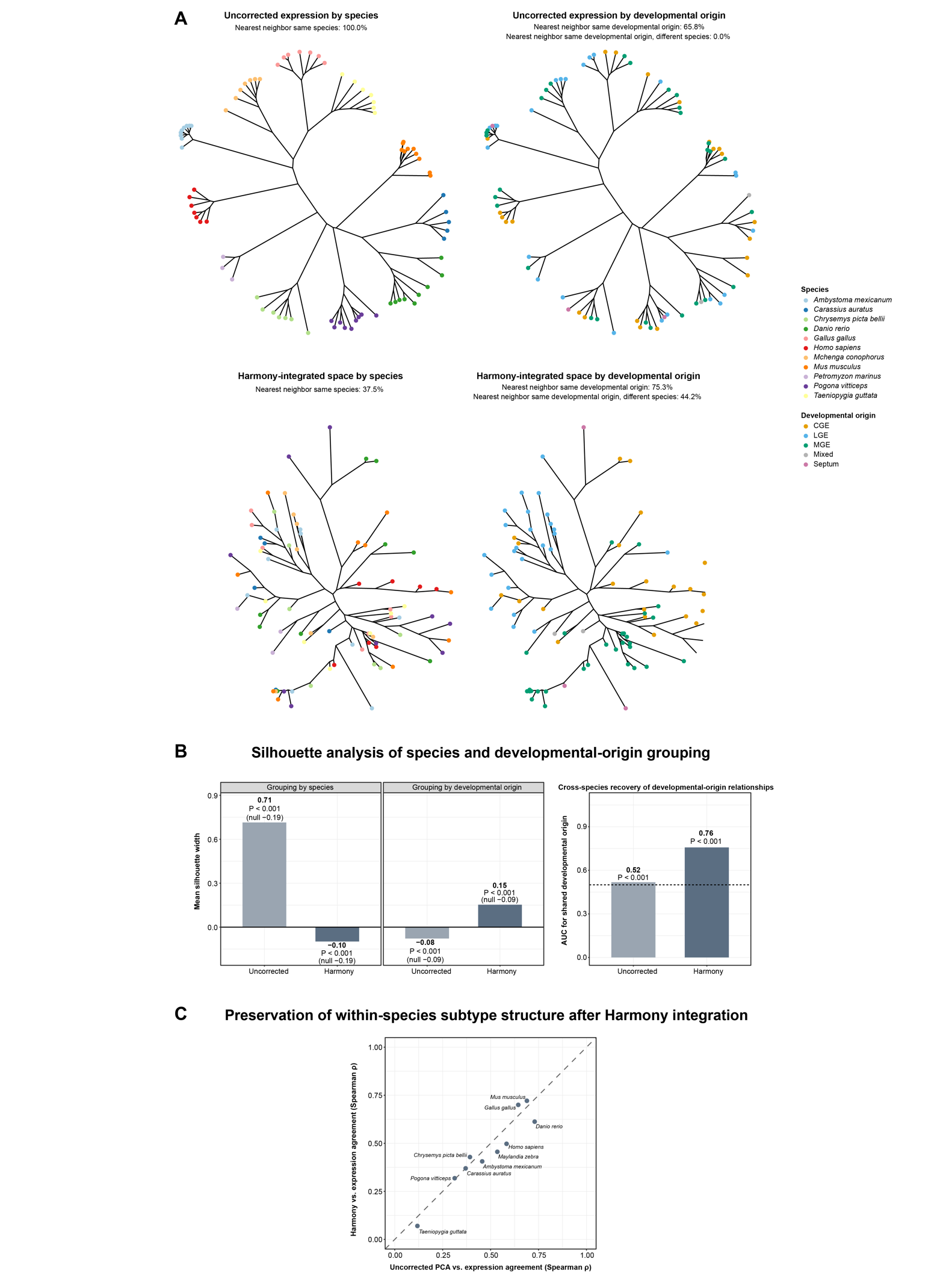

**Figure S14. Harmony integration reduces species-associated structure while revealing cross-species developmental organization of GABAergic subtypes.** (A) Comparison of GABAergic subtype relationships before and after Harmony integration. Unrooted distance-based trees were constructed from either uncorrected scGENUS gene-set expression or the Harmony-integrated scGENUS principal-component space and are shown with tips colored by species or developmental origin. In the uncorrected expression representation, nearest-neighbor relationships were strongly organized by species, with 100.0% occurring within the same species. Although 65.8% of nearest-neighbor relationships connected subtypes sharing developmental origin, none of these relationships occurred across species. In the Harmony-integrated space, the frequency of same-species nearest-neighbor relationships decreased to 37.5%, whereas 75.3% connected subtypes sharing developmental origin and 44.2% connected subtypes sharing developmental origin across different species. Developmental-origin categories include caudal ganglionic eminence (CGE), lateral ganglionic eminence (LGE), medial ganglionic eminence (MGE), mixed, and septal populations. (B) Quantitative assessment of species- and developmental-origin-associated organization before and after Harmony integration. Mean silhouette width was calculated using either species labels or developmental-origin labels, with higher positive values indicating stronger grouping by the corresponding annotation. Uncorrected expression showed strong species-associated organization (mean silhouette width = 0.71) but little developmental-origin organization (-0.08), whereas the Harmony-integrated representation reduced species-associated structure (-0.10) and increased grouping by developmental origin (0.15). Values in parentheses indicate the mean null expectation from 1,000 label permutations, and $P$ values were obtained from the corresponding permutation tests. Cross-species recovery of developmental-origin relationships was additionally quantified using the area under the receiver operating characteristic curve (AUC), where an AUC of 0.5 represents chance-level discrimination and higher values indicate greater similarity among cross-species subtype pairs sharing developmental origin than among pairs with different developmental origins. AUC increased from 0.52 for uncorrected expression to 0.76 after Harmony integration ($P<0.001$ for both comparisons). (C) Preservation of within-species GABAergic subtype structure after Harmony integration. For each species, pairwise relationships among GABAergic subtypes in uncorrected gene-set expression were compared with the corresponding relationships obtained from uncorrected PCA and Harmony-integrated space using Spearman correlation. Each point represents one species. The dashed diagonal denotes equal agreement with the uncorrected expression structure for PCA and Harmony; points close to this line indicate little additional distortion of within-species subtype structure after Harmony integration relative to dimensionality reduction alone.

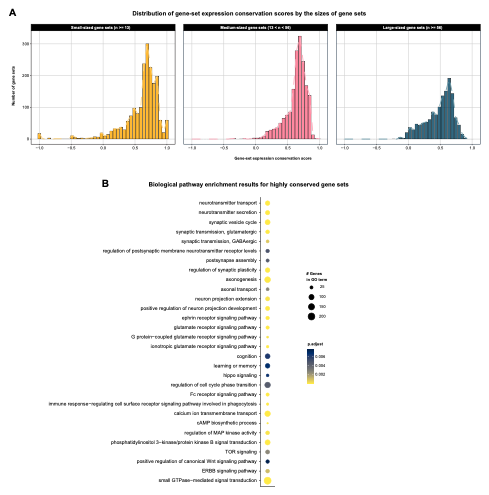

**Figure S15. Gene-set expression conservation is independent of gene-set size and functional enrichment of highly conserved gene sets.** (A) Distribution of gene-set expression conservation scores grouped by gene-set size. Conservation score distributions were broadly similar across gene-set size categories. (B) Gene Ontology (GO) Biological Process enrichment of the 1,086 highly conserved scGENUS gene sets, defined by a gene-set expression conservation score $>$ 0.75, using human orthologs for functional annotation. Circle size indicates the number of genes associated with each GO term, and color indicates the FDR adjusted $P$ value.

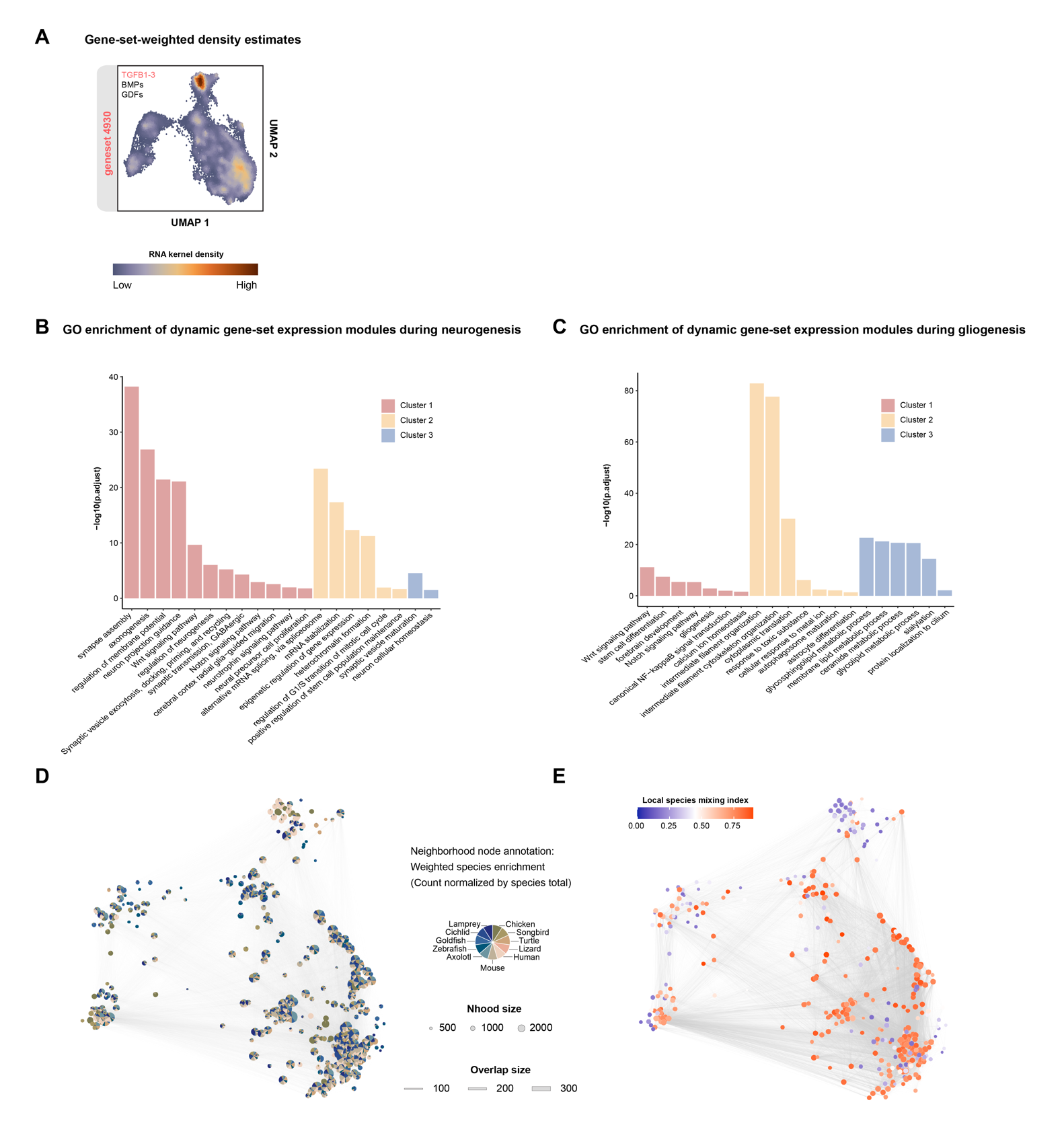

**Figure S16. TGFB-family gene-set density, functional enrichment of dynamic modules, and species mixing along radial glial differentiation trajectories.** (A) Gene-set-weighted density estimates for TGFB1-3 ligand-associated signaling projected onto the UMAP embedding. (B, C) Gene Ontology (GO) enrichment analysis of gene-set modules dynamically expressed along the neurogenic (B) and gliogenic (C) branches. Bars represent -log_10_(FDR adjusted $P$ values) and are colored according to the corresponding dynamic gene-set module. (D) Milo neighborhood graph of cells spanning the neurogenic and gliogenic branches. Each node represents a neighborhood, with node size proportional to neighborhood size and pie chart sectors showing species composition after normalization by the total number of cells from each species. Edges connect overlapping neighborhoods, with edge width proportional to the number of shared cells. (E) The same neighborhood graph colored by the local species mixing index, with higher values indicating greater mixing of species within neighborhoods. The median local species mixing index across neighborhoods was 0.67.

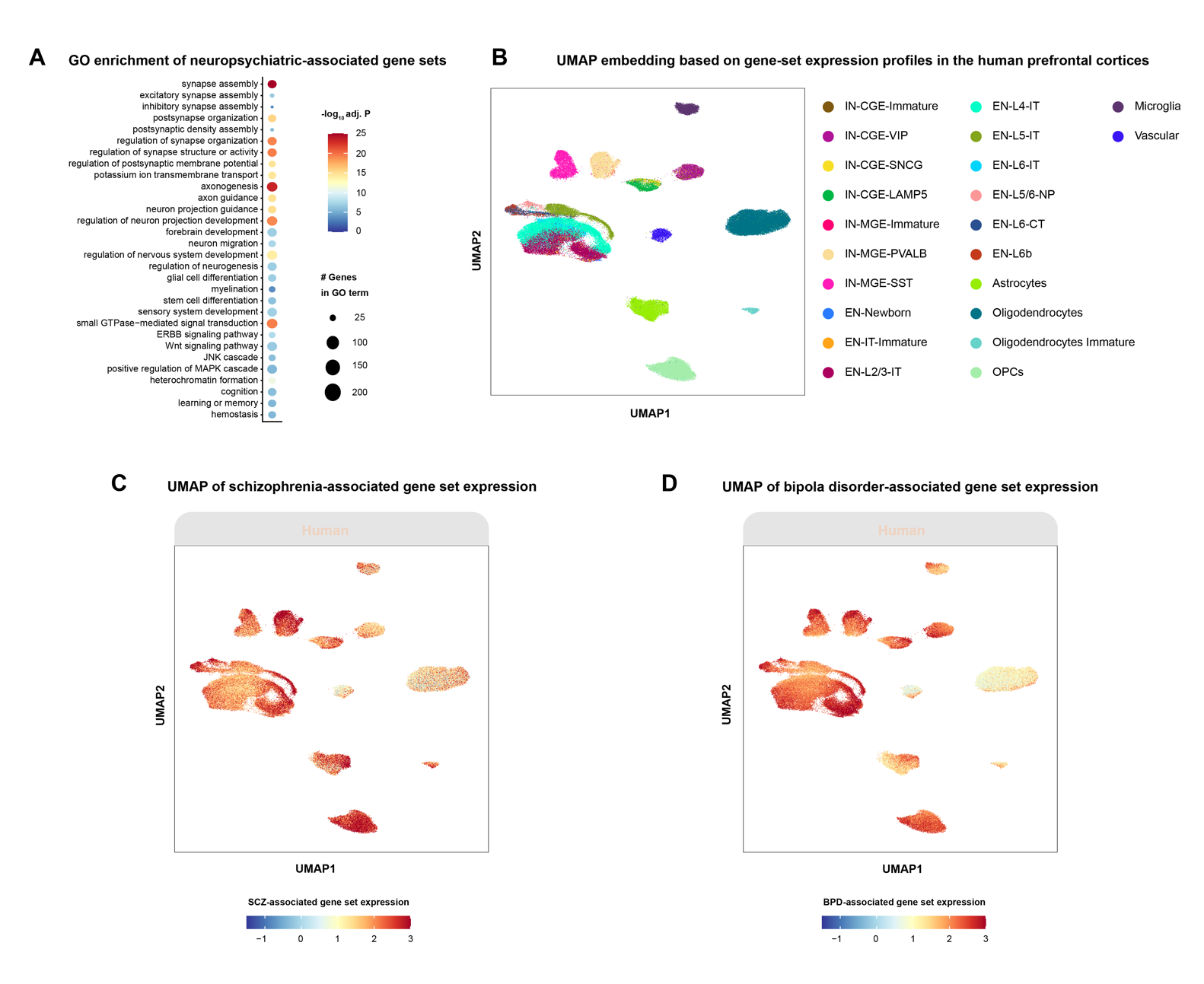

**Figure S17. Functional enrichment and human cell-type localization of neuropsychiatric-associated gene sets.** (A) Gene Ontology (GO) Biological Process enrichment of scGENUS gene sets significantly associated with neuropsychiatric traits by MAGMA. Circle size indicates the number of genes associated with each GO term, and color represents -log_10_(FDR adjusted $P$ values). (B) UMAP embedding of human prefrontal cortex cells generated from scGENUS gene-set expression profiles and colored by cell-type annotation. (C, D) Localization of schizophrenia (SCZ)- and bipolar disorder (BPD)-associated gene-set expression across the same human prefrontal cortex embedding. Module scores calculated from SCZ-associated gene sets are shown in (C), and module scores calculated from BPD-associated gene sets are shown in (D), with higher values indicating greater aggregate expression of the corresponding disease-associated gene sets.

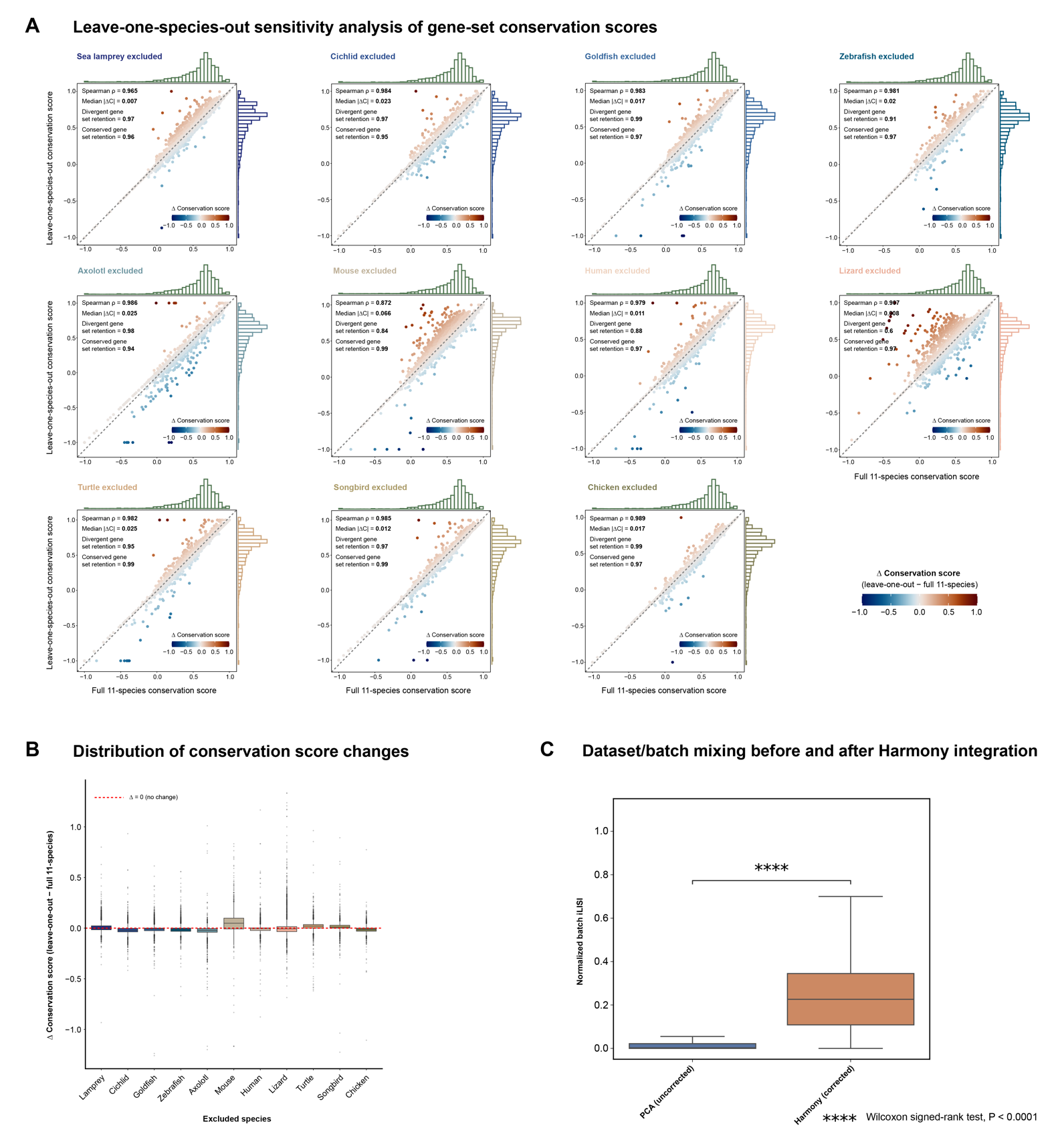

**Figure S18. Sensitivity of gene-set conservation and cross-species integration to dataset heterogeneity.** (A) Leave-one-species-out sensitivity analysis of scGENUS gene-set expression conservation scores. Conservation scores were recalculated after excluding each species in turn and compared with scores obtained from the full eleven-species analysis. Each point represents one gene set and is colored by the change in conservation score ($\Delta$conservation score = leave-one-out - full eleven-species score). Marginal distributions show conservation score distributions for the corresponding analyses. The dashed diagonal denotes equal conservation scores between the full and leave-one-out analyses. For each excluded species, annotations report the Spearman correlation between full and leave-one-out scores, the median absolute conservation score change, and the retention of gene sets classified as divergent or conserved in the full analysis. (B) Distribution of conservation score changes after exclusion of each species. Boxplots summarize $\Delta$conservation scores across gene sets for each leave-one-species-out analysis; the dashed horizontal line denotes no change ($\Delta=0$). Conservation scores remained centered near zero change across exclusions, indicating limited sensitivity to any single species dataset. (C) Local dataset/batch mixing before and after Harmony integration, quantified using normalized batch local inverse Simpson's index (iLISI) calculated for the same cells in the uncorrected PCA and Harmony-corrected representations. Higher normalized iLISI values indicate greater local mixing across dataset/batch labels. Harmony integration significantly increased batch mixing relative to the uncorrected PCA representation (paired Wilcoxon signed-rank test, $P<0.0001$).

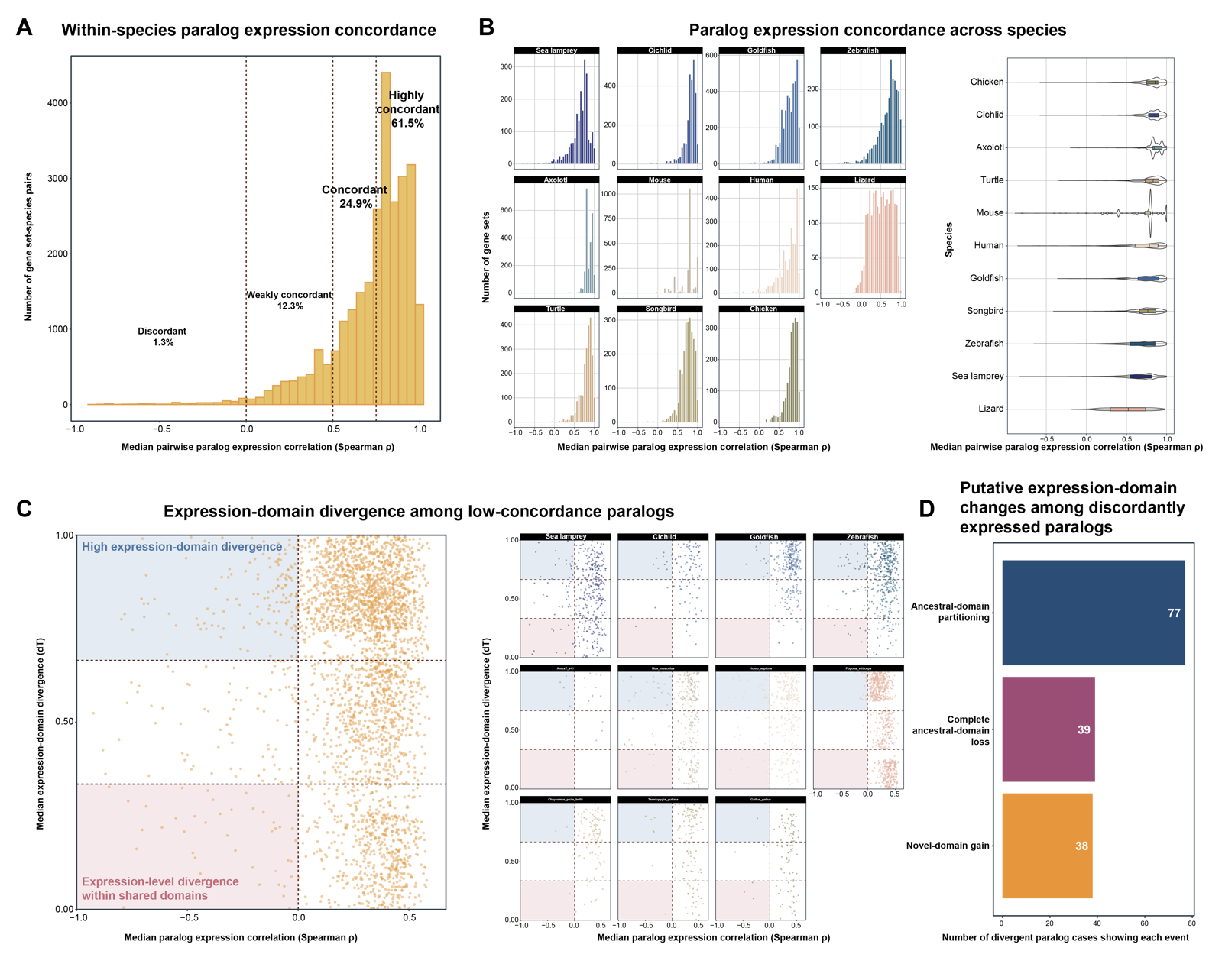

**Figure S19. Expression concordance and divergence among same-species paralogs within scGENUS gene sets.** (A) Distribution of within-species expression concordance among putative paralogs assigned to the same scGENUS gene set. For each gene set and species containing at least two same-species homologs, gene-level expression was aggregated across cell types and pairwise Spearman correlations were calculated between paralog expression profiles; the median pairwise correlation was used to summarize each gene-set-species combination. Gene-set-species pairs were classified as discordant ($\rho<0$), weakly concordant ($0\leq\rho<0.5$), concordant ($0.5\leq\rho<0.75$), or highly concordant ($\rho\geq0.75$). Overall, 61.5% were highly concordant, 24.9% concordant, 12.3% weakly concordant, and 1.3% discordant. (B) Distribution of median pairwise paralog-expression correlations across individual species. Histograms show the number of gene sets across the concordance range for each species, and violin plots summarize the corresponding species-level distributions. (C) Expression-domain divergence among low-concordance paralogs. For gene-set-species combinations with low expression concordance, expression-domain divergence was quantified as $dT=1-N_{both}/N_{either}$, where $N_{both}$ is the number of cell types in which both paralogs are expressed and $N_{either}$ is the number in which at least one paralog is expressed. Thus, $dT=0$ indicates identical expression domains and $dT=1$ indicates non-overlapping domains. The left panel shows the relationship between median expression correlation and median $dT$ across gene-set-species combinations, and the right panels show the corresponding distributions by species. Shaded regions highlight cases characterized by high expression-domain divergence or expression-level divergence within largely shared expression domains. (D) Putative expression-domain changes among discordantly expressed paralogs. For discordant cases with an appropriate single-copy outgroup, the closest phylogenetically relevant single-copy homolog was used as a proxy for the ancestral expression domain, and descendant paralog domains were compared with this reference. Among 108 classified paralog cases, 77 showed ancestral-domain partitioning, 39 showed complete ancestral-domain loss, and 38 showed novel-domain gain. These event categories were not mutually exclusive, and a single paralog case could exhibit more than one expression-domain change.

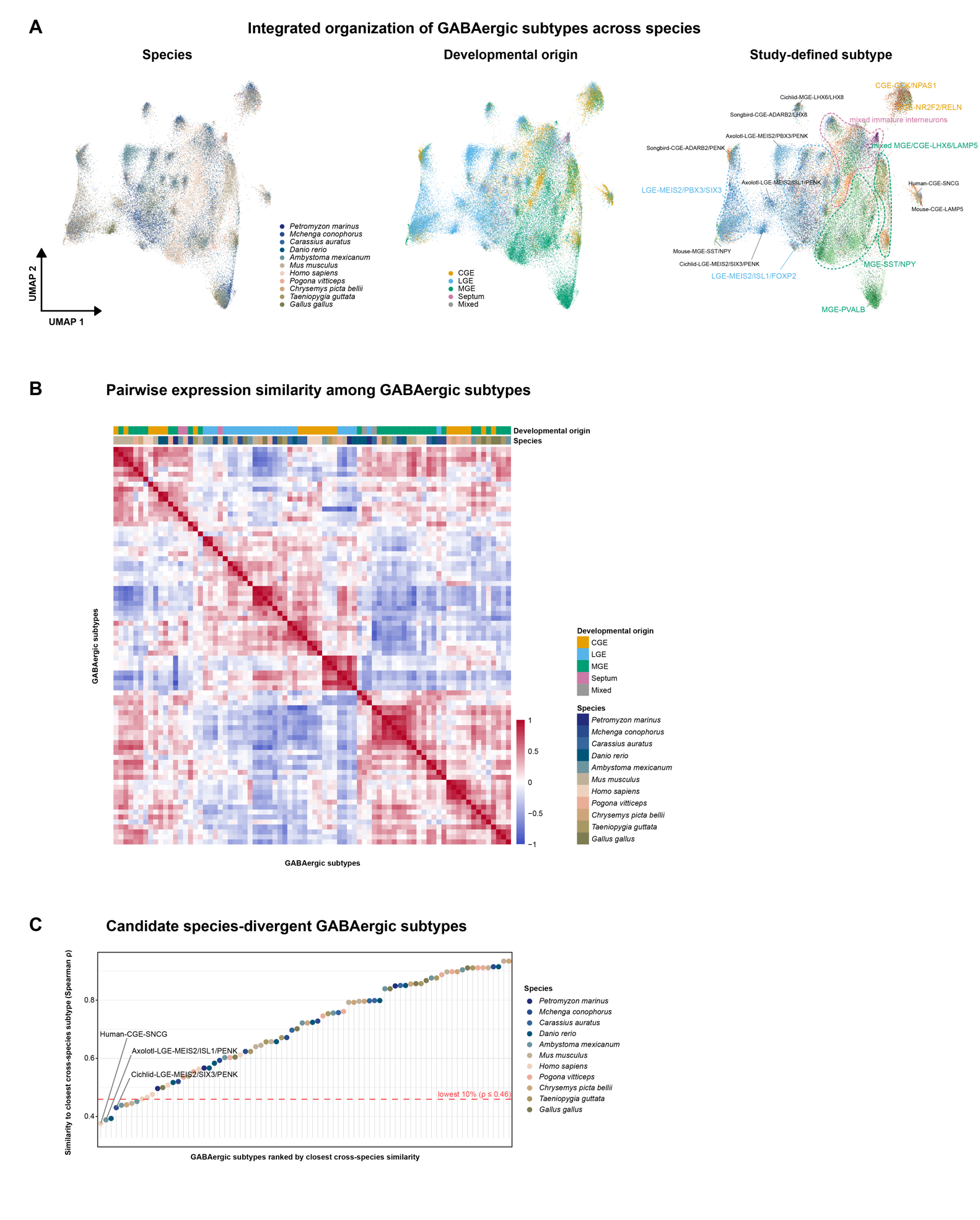

**Figure S20. Higher-resolution cross-species organization and divergence of GABAergic subtypes in the scGENUS representation.** (A) Integrated UMAP representation of GABAergic cells across the eleven vertebrate species, shown according to species, developmental origin, and study-defined subtype annotations. Developmental-origin categories include caudal ganglionic eminence (CGE), lateral ganglionic eminence (LGE), medial ganglionic eminence (MGE), septal, and mixed populations. The study-defined subtype view illustrates that finer-scale annotations from the individual source datasets occupy structured regions within the integrated GABAergic space. (B) Pairwise similarity among GABAergic subtypes based on Spearman correlation of subtype-level scGENUS gene-set expression profiles. Subtypes are ordered according to the distance-based subtype tree, and annotation bars indicate developmental origin and species. Higher positive correlations indicate greater similarity in gene-set expression profiles, whereas lower or negative correlations indicate greater transcriptional divergence. (C) Identification of candidate species-divergent GABAergic subtypes. For each subtype, similarity to its closest counterpart in another species was defined as the maximum Spearman correlation with any subtype from a different species. Subtypes are ranked by this closest cross-species similarity, with lower values indicating weaker correspondence to subtypes represented in other species. The dashed horizontal line marks the lowest 10% of the distribution ($\rho\leq0.46$), which were designated candidate species-divergent subtypes. Selected candidates are labeled for reference.

**SUPPLEMENTARY RESULTS**

**Robustness and functional characterization of scGENUS gene-set construction**

Because the downstream scGENUS analyses depend on gene sets derived from the multispecies protein-homology graph, we systematically evaluated whether gene-set construction was sensitive to protein sequence properties or major graph-construction and clustering parameters.

We first examined whether protein length disproportionately influenced homology-graph connectivity or final gene-set size. Protein length showed only modest positive associations with graph degree (Spearman $\rho=0.27$) and gene-set size ($\rho=0.31$; Fig. S1A,B). To assess whether gene-set membership could be disproportionately supported by short local sequence similarities, overlapping BLASTp high-scoring pairs (HSPs) were merged along each protein participating in an edge, and the smaller of the two resulting coverage fractions was used as a conservative measure of protein alignment coverage. In 94.7% of gene sets, edges connecting genes within the same gene set showed greater median alignment coverage than edges connecting genes assigned to different gene sets, with a median within-minus-between difference of 0.43 (Fig. S1C). Larger gene sets showed lower median within-set alignment coverage ($\rho=-0.40$; Fig. S1D), indicating a greater contribution from partial sequence similarities in larger homology communities. Nevertheless, after removing edges with minimum protein alignment coverage below 30%, 87.2% of evaluable gene sets retained more than 90% of their genes within the largest connected component (Fig. S1E), indicating that low-coverage sequence similarities were generally not the sole structural support for gene-set connectivity.

We next evaluated sensitivity to the definition of reciprocal BLASTp edge weights. Bit scores obtained in the two reciprocal search directions were highly concordant ($\rho=0.99$), and 80.4% of reciprocal edges differed by no more than 5% between directions (Fig. S2A). Gene sets were then reconstructed using the mean or minimum reciprocal bit score instead of the maximum used in the primary analysis. Across the maximum-, mean-, and minimum-weighted constructions, 5,298 gene sets (95.8%) were represented across all three collections under the matching criterion used for comparison (Fig. S2B). For each primary maximum-weighted gene set, we further calculated its highest Jaccard similarity to any gene set obtained under either alternative weighting definition. The median best-match Jaccard index was 1.00 (IQR, 1.00–1.00), and 99.3% of primary gene sets had a best-match Jaccard index of at least 0.8 (Fig. S2C). Thus, gene-set composition was largely insensitive to the particular reciprocal bit-score summary used to weight graph edges.

We further tested sensitivity to BLASTp stringency by applying progressively more restrictive reciprocal E-value thresholds. Relative to the primary $E\leq{10}^{-6}$ graph containing 70,279,674 edges, thresholds of $E\leq{10}^{-10}$ and $E\leq{10}^{-20}$ retained 58,983,974 (83.9%) and 42,253,718 (60.1%) edges, respectively (Fig. S3A). Despite this substantial edge removal, 71.4% of primary gene sets remained fully intact at ${10}^{-10}$, whereas 27.5% lost fewer than half of their genes. At ${10}^{-20}$, 57.5% remained fully intact, 37.7% lost fewer than half of their genes, and only 4.7% lost at least half of their original membership (Fig. S3B). We additionally reconstructed gene sets de novo from the $E\leq{10}^{-10}$ graph using the same clustering procedure as the primary analysis. These reconstructed gene sets showed a median best-match Jaccard index of 1.00 (IQR, 0.92–1.00) relative to the primary collection; 81.7% had $J_{max}\geq0.8$, and 67.2% were identical to their best-matching primary gene set (Fig. S3C). Among non-identical reconstructed gene sets, 93% had at least 90% of their genes contained within a single primary gene set, further indicating that increased BLASTp stringency generally produced modest boundary changes rather than wholesale reorganization of gene-set membership.

We next examined the effect of Leiden resolution on graph partitioning. Across the full resolution scan, increasing resolution progressively increased the number of communities while decreasing median community size (Fig. S4A,B). We evaluated the fraction of communities containing genes from all 11 species across resolutions and compared cross-species representation criteria of 40%, 50%, and 60% (Fig. S4C). The primary 50% criterion identified a broad range in which community number, community size, and cross-species representation remained comparatively stable. Relaxing the criterion to 40% extended the retained range only to an additional higher resolution at which community-size distributions became less stable, whereas a 60% requirement was overly restrictive and did not retain an acceptable resolution range. These analyses support the 50% criterion as a practical balance between cross-species representation and partition stability rather than as a uniquely optimal threshold.

Finally, we characterized the structural and functional properties of the final collection of 5,419 scGENUS gene sets. Gene sets varied substantially in size but generally showed broad phylogenetic representation (Fig. S5A,B): 65% contained genes from all 11 species, and an additional 12% contained genes from 10 species. Based on the median number of genes per represented species, 50.8% of gene sets were predominantly single-copy, 15.1% showed low-copy expansion, and 34.1% showed broader within-species expansion (Fig. S5C). To independently assess functional coherence, we quantified the fraction of within-set gene pairs sharing at least one Gene Ontology Biological Process annotation and compared each scGENUS gene set with size-matched random gene groups. scGENUS gene sets showed greater Biological Process coherence than their random counterparts in 90.7% of human comparisons and 90.8% of mouse comparisons (paired Wilcoxon signed-rank test, $P<2.2\times{10}^{-16}$ for both species; Fig. S5D). A small subset showed zero Biological Process coherence (6.5% in human and 6.6% in mouse), but these cases were strongly enriched for gene sets containing three or fewer genes with available Biological Process annotations. Among these zero-BP-coherence sets, 34% in human and 46% in mouse nevertheless shared at least one Molecular Function or Cellular Component annotation.

Together, these analyses indicate that the scGENUS gene-set representation is broadly robust to the principal sequence-similarity, edge-weighting, and graph-partitioning choices examined here, while also defining important limitations of the approach. scGENUS gene sets should be interpreted as sequence-homology communities derived from a multispecies sequence-homology graph rather than as uniquely resolved gene families, orthogroups, or functionally homogeneous pathways. Larger gene sets contain a greater contribution from partial sequence similarities, and individual community boundaries can shift under alternative parameterizations. Moreover, protein alignment coverage provides an indirect measure of the extent of sequence similarity but does not distinguish similarities arising specifically from shared protein domains or repetitive architectures from other forms of local homology. Nevertheless, the broad preservation of gene-set structure under graph perturbation, together with substantially greater functional coherence than size-matched random groups in two independently annotated species, supports the use of these communities as shared features for downstream comparative single-cell analyses.

**Expression concordance and divergence among same-species homologs within scGENUS gene sets**

To evaluate whether mean aggregation could obscure paralog-specific expression differences, we examined expression concordance among same-species homologs assigned to the same scGENUS gene set. Across gene-set–species combinations containing at least two same-species homologs, 61.5% were highly concordant ($\rho\geq0.75$), 24.9% were concordant ($0.5\leq\rho<0.75$), 12.3% were weakly concordant ($0\leq\rho<0.5$), and only 1.3% were discordant ($\rho<0$; Fig. S19A). Thus, 86.4% of evaluated combinations showed median pairwise expression correlations of at least 0.5. Species-level distributions were broadly similar, although the frequency of low-concordance cases varied among species (Fig. S19B).

Among low-concordance cases, reduced expression correlation was associated with varying degrees of expression-domain divergence across cell types (Fig. S19C). Some homolog pairs showed divergence primarily through separation of their cell-type expression domains, whereas others retained broadly overlapping domains but differed in relative expression levels. For discordant cases with an appropriate single-copy outgroup, comparison with the closest phylogenetically relevant single-copy homolog identified putative changes in expression-domain organization. Among 108 classified cases, 77 showed ancestral-domain partitioning, 39 showed complete ancestral-domain loss, and 38 showed novel-domain gain; these categories were not mutually exclusive (Fig. S19D).

Together, these results indicate that mean aggregation is appropriate for the majority of scGENUS gene sets, in which same-species homologs retain broadly concordant expression, while also identifying a minority of cases in which paralog-specific expression divergence may be attenuated by aggregation.

**Benchmarking scGENUS against alternative cross-species integration approaches**

To quantitatively evaluate cross-species integration performance, we compared scGENUS with SATURN, CAMEX, SAMap, and a one-to-one ortholog representation integrated using Harmony (Harmony O2O). We additionally evaluated two negative controls designed to test whether integration performance could be reproduced without the evolutionary organization of the scGENUS gene sets: size-matched random gene sets and gene sets derived from a rewired homology graph. Integration quality was assessed using complementary metrics adapted from the scIB framework, including batch silhouette score, graph iLISI, and graph connectivity for batch correction, and cell-type silhouette score, isolated-label preservation, and graph cLISI for biological conservation (Fig. S8A; Supplementary Data File 8). UMAP embeddings colored by species and cell type were additionally examined as qualitative representations of the resulting integrated spaces (Fig. S8B,C).

scGENUS achieved an aggregate batch-correction score of 0.70 and an aggregate biological-conservation score of 0.69, yielding the highest overall aggregate score (0.70) among the evaluated methods (Fig. S8A). SATURN achieved a slightly higher biological-conservation score (0.72) but lower batch correction (0.55), resulting in an overall score of 0.66, whereas CAMEX achieved batch-correction and biological-conservation scores of 0.65 and 0.63, respectively, with an overall score of 0.64. Harmony O2O and SAMap showed lower overall scores of 0.57 and 0.44, respectively. At the individual-metric level, scGENUS showed high batch silhouette (0.90), graph iLISI (0.22), and graph connectivity (0.99) while maintaining cell-type silhouette (0.54), isolated-label preservation (0.53), and graph cLISI (1.00). No single method performed best across all individual metrics; rather, scGENUS provided the strongest overall balance between cross-species mixing and preservation of cell-type structure.

The negative controls further distinguished evolutionarily informed feature construction from generic feature aggregation. Size-matched random gene sets retained relatively strong biological conservation (0.68) but showed substantially weaker batch correction (0.38), yielding an overall score of 0.56. Gene sets derived from the rewired homology graph showed reduced batch correction (0.44) and biological conservation (0.57), with an overall score of 0.52. Thus, neither random aggregation of genes nor disruption of the underlying homology-graph structure reproduced the performance of scGENUS. Together with the lower performance of the one-to-one ortholog baseline, these results indicate that the evolutionary organization of the scGENUS gene-set representation contributes to effective cross-species integration rather than the observed performance arising solely from feature aggregation or downstream batch correction.

**Sensitivity of cross-species analyses to dataset heterogeneity**

Because the eleven-species atlas integrates datasets generated across different studies, sequencing platforms, developmental stages, and anatomical sampling schemes, we evaluated whether major cross-species patterns were sensitive to dataset heterogeneity. Dataset- and sample-level characteristics, including source study, sequencing modality and platform/chemistry, biological replicates, anatomical region, developmental stage, sex where available, retained cell numbers, and major cell-type composition, are summarized in Supplementary Data File 7. Study, platform, anatomical sampling, and species were frequently nested or confounded, preventing their independent effects from being reliably estimated.

We first assessed whether Harmony reduced detectable within-species batch structure by comparing graph iLISI values between the pre-Harmony PCA and Harmony-corrected PCA representations for datasets containing multiple batches. Batch mixing increased after Harmony correction, indicating reduced batch-associated structure within species (Fig. S18A). Importantly, this analysis evaluates correction of within-species technical variation without requiring platform or study effects to be separated from species identity.

We next evaluated the robustness of gene-set conservation scores using a leave-one-species-out analysis. Conservation scores were recalculated eleven times, each time excluding one species, and compared with estimates obtained from the complete dataset. Conservation scores remained highly concordant with the full analysis across all leave-one-species-out comparisons, with generally small changes in individual gene-set scores (Fig. S18B,C). Exclusion of sea lamprey, the most phylogenetically distant species in the dataset, still produced a strong correlation with the full-data estimates ($\rho=0.965$) and a median absolute change of approximately 0.007. These results indicate that the overall gene-set conservation landscape is robust to the removal of individual species datasets, while the confounding among species, study, platform, and anatomical sampling remains an inherent limitation of the available comparative datasets.

**Species mixing and strongly species-biased neighborhoods**

Neighborhood-level analysis revealed extensive cross-species mixing across the integrated forebrain atlas while also identifying localized regions of lineage-associated divergence. Milo-defined neighborhoods were constructed in the integrated scGENUS space, and species composition was normalized relative to global species frequencies to account for unequal sampling across datasets (Fig. S10). Most neighborhoods contained cells from multiple species, although the degree of local species mixing varied across the atlas and among major cell classes.

To systematically characterize the strongest species-associated structure, we examined the distribution of normalized Shannon entropy across individual neighborhoods. The distribution contained a distinct low-entropy tail, with a local minimum at $h=0.29$; neighborhoods below this threshold were therefore classified as strongly species-biased (Fig. S11A). These comprised 533 of 2,260 neighborhoods (23.6%) and contained 14.6% of all cells, indicating that strongly biased neighborhoods represent a substantial but localized component of the integrated atlas rather than its predominant organization.

The prevalence of strongly species-biased neighborhoods varied markedly among species and cell classes. Approximately 45.0% of human cells and 25.9% of mouse cells occurred within strongly biased neighborhoods, compared with 5.4% of axolotl cells (Fig. S11B). At the cell-class level, approximately 65% of oligodendrocyte neighborhoods were strongly species-biased, whereas only approximately 6% of radial glial neighborhoods met this criterion (Fig. S11C). Strongly species-biased neighborhoods were also smaller than the remaining neighborhoods, with median sizes of 214 versus 443 cells, respectively (Fig. S11D).

Together, these analyses show that broad cross-species mixing coexists with localized species-associated transcriptional structure. Importantly, strongly species-biased neighborhoods are not interpreted as lineage-specific or novel cell types, because such patterns may reflect lineage-associated cell states, anatomical sampling differences, or residual dataset heterogeneity in addition to genuine cellular innovation.
