## Supplemental Data 2 for "Evolutionarily informed gene sets reveal conserved and lineage-modified transcriptional programs during vertebrate forebrain evolution": Description_of_Additional_Supplementary_Data_Files.docx

File Name: Supplementary Data File 1

Description: Annotation of scGENUS-derived gene sets. Each gene set is assigned a unique identifier (gene-set ID) and associated with the graph-partitioning resolution(s) at which it was identified, the number of genes within the gene set, and corresponding gene symbols. Gene membership is reported separately for each species, with one worksheet per species.

File Name: Supplementary Data File 2

Description: Results of differential gene-set expression testing used to identify cell-type-enriched gene sets. Each worksheet corresponds to one cell type and reports Wilcoxon rank-sum test statistics for each gene set, including $P$ value, average log2 fold change, percentage of cells expressing the gene set in the focal cell type and in all other cell types, adjusted $P$ value, and the difference in expression percentages between the two groups.

File Name: Supplementary Data File 3

Description: Cell-type metadata for each species included in the study. Each worksheet corresponds to one species and reports the original cell-cluster annotation (Original_annotation), primary cell class (Class), secondary cell classification (Subclass), harmonized cell-type annotation (Celltype), anatomical-region annotation (Anatomical_region), and developmental-origin classification of GABAergic interneuron subtypes (Interneuron_subtype).

File Name: Supplementary Data File 4

Description: Metadata for Milo-defined KNN graph neighborhoods used in cross-species compositional analyses. Each row corresponds to one neighborhood. The columns x and y report coordinates used to visualize the neighborhood graph. Species-specific columns report neighborhood species composition normalized relative to global species frequencies. Species abbreviations correspond to *Ambystoma mexicanum*, *Carassius auratus*, *Chrysemys picta bellii*, *Danio rerio*, *Gallus gallus*, *Homo sapiens*, *Mus musculus*, *Mchenga conophorus*, *Petromyzon marinus*, *Pogona vitticeps*, and *Taeniopygia guttata*. The column entropy_scaled reports normalized Shannon entropy of species composition within each neighborhood, with lower values indicating greater species bias and higher values indicating greater cross-species mixing. The column radius is proportional to the number of cells contained in each neighborhood, and km_cluster reports the k-means cluster assignment of each neighborhood based on the neighborhood-graph representation.

File Name: Supplementary Data File 5

Description: Gene sets exhibiting state- and species-dependent expression dynamics along radial glial developmental trajectories. Sheet 1 reports results of state-dependent analysis testing for differences in gene-set expression dynamics between neurogenic and gliogenic branches using natural-spline regression models. Sheets 2 and 3 report species-dependent analyses of gene-set expression dynamics along pseudotime within the neurogenic and gliogenic branches, respectively, using generalized linear models implemented in Monocle3, with species-by-pseudotime interaction terms capturing species-dependent differences in expression dynamics.

File Name: Supplementary Data File 6

Description: Results of MAGMA gene-set enrichment analyses testing associations between scGENUS-derived gene sets and human cognitive traits and neuropsychiatric disorders. Each row corresponds to one gene-set–trait pair and reports the associated trait, enrichment statistics, and standardized effect size.

File Name: Supplementary Data File 7

Description: Dataset- and sample-level metadata for the eleven vertebrate forebrain transcriptomic datasets included in the scGENUS analysis. The table reports species, source study and DOI, sequencing modality, library platform and chemistry, number of biological replicates, sample ID, anatomical region, developmental stage, sex, number of retained cells, and major cell types represented in each dataset. Sample-level entries are grouped according to their corresponding species and source study.

File Name: Supplementary Data File 8

Description: scIB benchmarking results comparing cross-species integration performance across scGENUS, SATURN, CAMEX, Harmony using one-to-one ortholog features (Harmony O2O), SAMap, and two negative controls based on size-matched random gene sets and a rewired homology graph. Batch-correction metrics include batch silhouette score, graph iLISI for batch mixing, and graph connectivity. Biological-conservation metrics include cell-type-label silhouette score, isolated-label score, and graph cLISI for preservation of cell-type structure. Reported benchmark scores are oriented such that higher values indicate better performance.
